# mrMLM v4.0: An R Platform for Multi-locus Genome-wide Association Studies

**DOI:** 10.1101/2020.03.04.976464

**Authors:** Ya-Wen Zhang, Cox Lwaka Tamba, Yang-Jun Wen, Pei Li, Wen-Long Ren, Yuan-Li Ni, Jun Gao, Yuan-Ming Zhang

## Abstract

Previous studies reported that some important loci are missed in single-locus genome-wide association studies (GWAS), especially because of the large phenotypic error in field experiments. To solve this issue, multi-locus GWAS methods have been recommended. However, only a few software packages are available. Therefore, an R software mrMLM, which includes our six multi-locus methods, was developed. mrMLM includes three components: dataset input, parameter setting and result output. The fread function in data.table is used to quickly read datasets, especially big datasets, and the doParallel package is used to conduct parallel computation using multiple CPUs. In addition, the graphical user interface software mrMLM.GUI v4.0, built upon Shiny, is also available. To confirm the correctness of the above programs, the same simulation datasets as used in previous studies, along with three real datasets, were re-analyzed by all the methods in mrMLM v4.0 and three widely-used methods. The results confirmed the advantages of our multi-locus methods over the current methods. The conclusion is also consistent with those in a Research Topic in Frontiers in Plant Science. Although a less stringent significance threshold is adopted, the false positive rates are effectively controlled. mrMLM is publicly available at https://cran.r-project.org/web/packages/mrMLM/index.html or https://bigd.big.ac.cn/biocode/tools/BT007077 as an open-source software.

## Introduction

Since the establishment of the mixed linear model (MLM) framework of genome-wide association studies (GWAS) [1,2], the MLM-based GWAS methodologies have been widely used to identify many important loci for complex traits in animals, plants and humans. With the technological advances in molecular biology, a huge number of markers is easily obtained. However, this brings new computational and analytic challenges. The MLM-based single-marker association in genome-wide scans proves its feasibility. To increase statistical power and decrease running time in quantitative trait nucleotide (QTN) detection, a series of additional MLM-based methods have been proposed. For example, Kang et al. [3] proposed an efficient mixed model association (EMMA), which was then extended to EMMAX [4] and GEMMA [5], while Zhang et al. [6] reported a compressed MLM (CMLM) method, which was then extended to ECMLM [7] and SUPER [8]. In addition, other methods have also been developed, i.e., GRAMMAR-Gamma [9], FaST-LMM [10], FaST-LMM-Select [11] and BOLT-LMM [12]. All the above methods have been subjected to multiple testing. To control the false positive rate in such tests, the Bonferroni correction is frequently adopted. However, this correction is often too conservative to detect many important loci.

To detect more QTNs with a low false positive rate, multi-locus methods have been recommended. This recommendation was implemented for the first time by Segura et al. [13]. Thereafter, Liu et al. [14] developed FarmCPU. Based on the advantages of the random model of QTN effect over the fixed model [15], recently, we have developed six multi-locus methods: mrMLM [16], FASTmrMLM [17], FASTmrEMMA [18], ISIS EM-BLASSO [19], pLARmEB [20] and pKWmEB [21] (Files S1 and S2). These methods include two stages. First, various algorithms are used to select all the potentially associated markers. Second, these selected markers are put in one model, all the effects in this model are obtained by empirical Bayes, and all the non-zero effects are further identified by likelihood ratio test for true QTNs. Although a less stringent criterion of significance is adopted, these methods have high power and accuracy and a low false positive rate.

In the GWAS software, there are many packages available, for example, PLINK [22], TASSEL [23], EMMA [3], EAMMAX [4], GEMMA [5] and GAPIT [24,25] (File S2). However, these packages are almost based on single-marker association in genome scans. To popularize our multi-locus GWAS methods, we integrated all the six multi-locus approaches into one R package, named mrMLM (Fig S1).

## Implementation

mrMLM v4.0 includes three parts (Fig 1): dataset input, parameter setting, and result output. In the dataset input module, users need to input trait phenotypes and marker genotypes, which are indicated by fileGen and filePhe, respectively. Two types of file formats, *.csv and *.txt, are available. Marker genotypes may be indicated by mrMLM numeric (or character) and Hapmap formats, and are used to calculate both kinship (using mrMLM or EMMA [3]) and population structure (using Structure [26] or fastSTRUCTURE [27]) matrices. This software has also an option to input kinship matrix, population structure matrix and covariate table, which are indicated by fileKin, filePS and fileCov, respectively. In the parameter setting module, users need to set eighteen parameters. Among these parameters, Likelihood, SearchRadius and SelectVariable are specific to method; eight may be default or set by users; fileGen, filePhe, Genformat, method, trait, CriLOD and dir must be set by users. In the result output module, intermediate and final results and ten plots (*.png, *.tiff, *.jpeg, *.pdf) are output to the path that users have previously set, i.e., dir=“D:/Users”. The software is started in a computer or server via the below codes (File S5) mrMLM(fileGen=“D:/Users/Genotype_num.csv”,filePhe=“D:/Users/Phenotype.csv”,f ileKin=“D:/Users/Kinship.csv”,filePS=“D:/Users/PopStr.csv”,PopStrType=“Q”,fileCo v=“D:/Users/Covariate.csv”,Genformat=“Num”,method=c(“mrMLM”,“FASTmrML M”,“FASTmrEMMA”,“pLARmEB”,“pKWmEB”,“ISIS EM-BLASSO”),Likelihood=“REML”,trait=1:3,SearchRadius=20,CriLOD=3,SelectVariable=50,Bootstrap=FALSE,DrawPlot=FALSE,Plotformat=“jpeg”,Resolution=“Low”,dir=“D:/users”)

**Figure 1.**
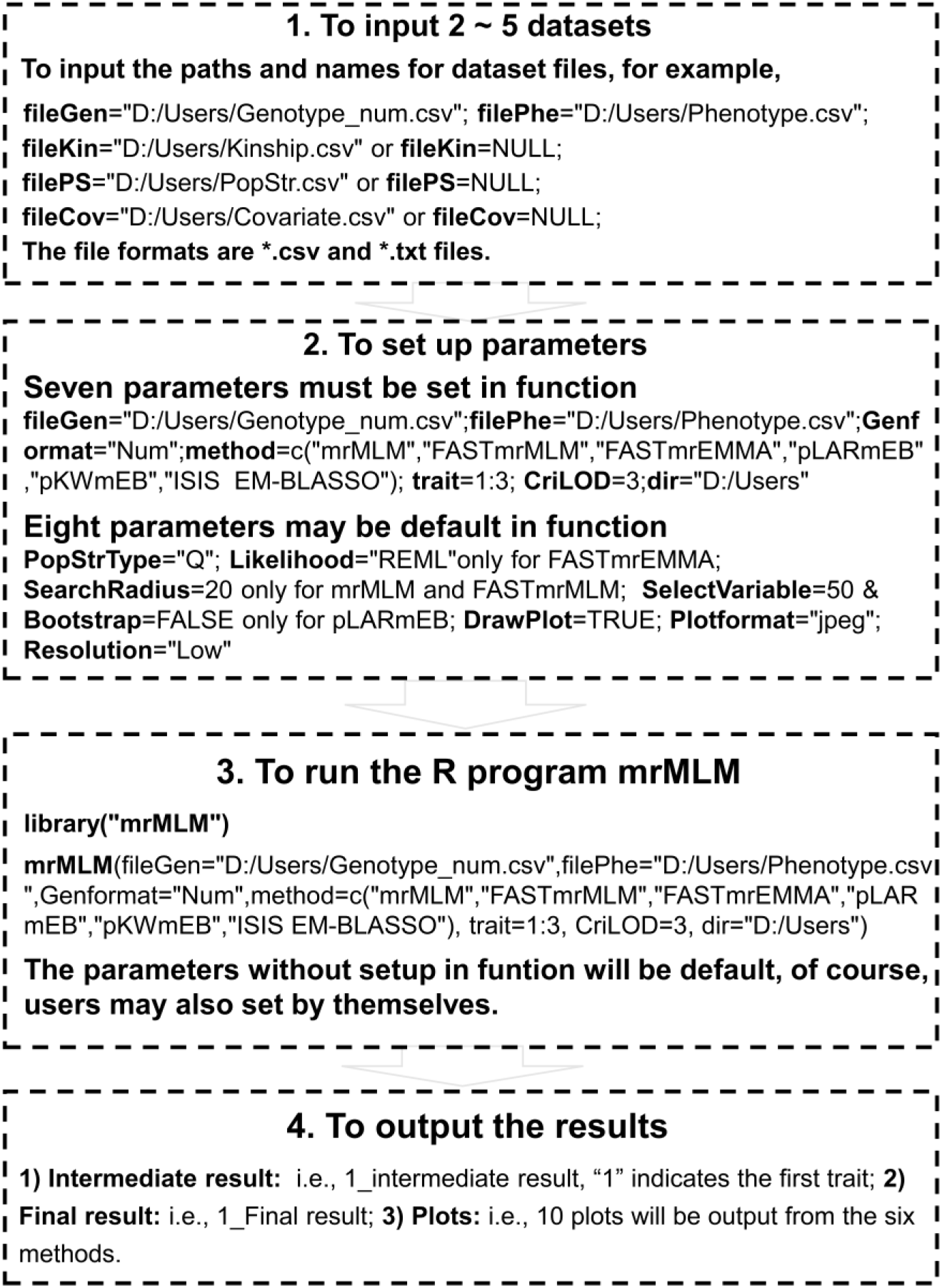
The framework of the R software package mrMLM v4.0.

R core is a single-threaded program and its computing mode limits its ability to handle large-scale data. In mrMLM v4.0, however, several packages in R were used to perform parallel calculation. First, detectCores() and makeCluster(cl.cores) in a parallel package were used, respectively, to detect the number of CPUs on the current host and create a set of copies of R running in parallel and communicating over sockets. Then, registerDoParallel(cl) in doParallel package was used to register the parallel backend with the foreach package. Third, ‘for’ loop was replaced by foreach(i=1:n,.combine=‘rbind’)%dopar%{…} in foreach package. Finally, stopCluster(cl) in parallel package was used to stop the above parallel calculation.

fread function in data.table was used to quickly read datasets, especially big datasets. For reading one genetic dataset with 500 individuals and one million markers, fread was three times faster (72.84 s) than read.csv does (201.45 s). Meanwhile, we utilized the advantages of package bigmemory, which can create, store, access and manipulate massive matrices, to define the huge genotypic matrix with the aid of the big.matrix() function. This largely saves the running time, especially for massive genetic matrix.

The graphical user interface (GUI) software mrMLM.GUI v4.0, built upon Shiny, is available as well. The interactive GUI is started via the two commands “library(mrMLM.GUI)” and “mrMLM.GUI()” (File S6). The next operation can be done through clicking the mouse conveniently.

## Results

To test the performances of the software package mrMLM v4.0, three real datasets in rice [28], maize [29] and Simmental beef cattle [30] were downloaded from the Rice SNP-Seek Database, the Maizego and the Dryad Digital Repository, respectively (File S3). In the above three datasets, the traits of interest were grain width, oil concentration and kidney weight, respectively; the numbers of phenotypic accessions were 2262, 362, and 1136, respectively; the numbers of markers were 1.01, 1.06, and 0.67 million, respectively (File S3).

### Influence of various factors on QTN detection using mrMLM v4.0

To investigate the effect of the number of markers on running time, four samples with various numbers of markers (0.2, 0.5, 0.8 and 1.01 million) and a fixed sample size (500) were sampled from the first real dataset [28]. As a result, it took 0.23, 0.66, 1.18 and 1.61 hours, respectively (Fig 2a), indicating the increase of running time with the increase of the number of markers. To investigate the effect of sample size on running time, four samples with various sizes (300, 600, 900, 1200 and 2262 accessions) and all the markers (1.01 million) were sampled from Wang et al. [28]. As a result, it took 0.37, 0.78, 1.30, 2.04 and 9.56 hours, respectively (Fig 2b). This indicates that large samples take much more running times than small samples.

**Figure 2.**
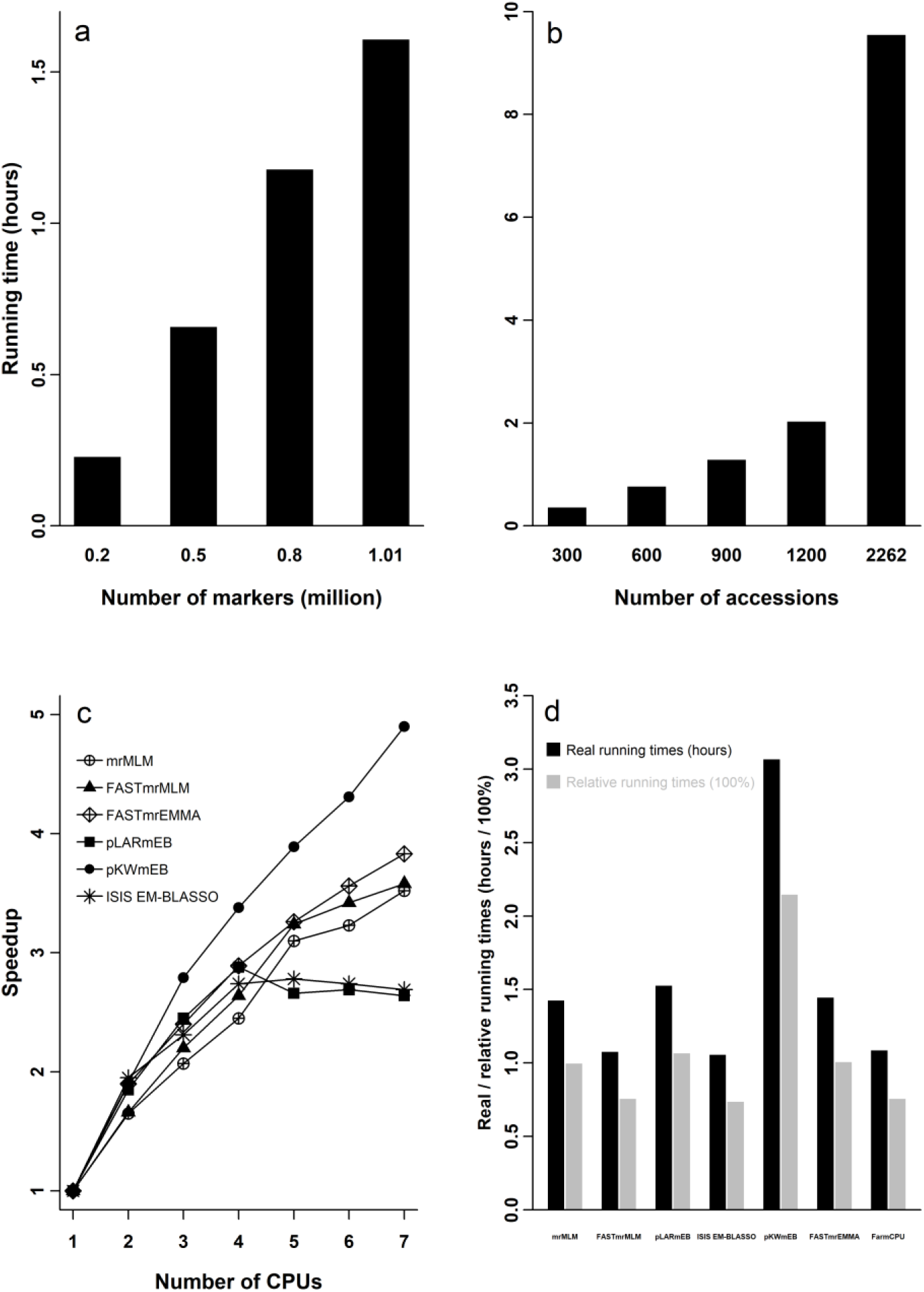
The performances of the R software package mrMLM v4.0 under various situations in the detection of QTNs for rice grain width. **a.** The numbers of markers. **b.** The number of individuals. **c.** The number of CPUs. **d.** Various GWAS approaches. The dataset was derived from Wang et al. [28].

To investigate the effect of the number of CPUs on speedup, one sample with 500 individuals and 1.01 million markers was analyzed by the mrMLM method under various numbers of CPUs (1 to 7). As a result, the speedup was 1.00, 1.65, 2.07, 2.45, 3.10, 3.23, 3.52, respectively (Fig 2c; Table S1), indicating the effectiveness of parallel computing under various numbers of CPUs. The relatively small speedups with 5 to 7 CPUs for pLARmEB and ISIS EM-BLASSO may be due to the fact that their potentially associated markers were determined at the chromosome and genome levels, respectively (Table S1). To compare the running times of various methods, one sample with 500 individuals and 1.01 million markers was analyzed by seven methods mrMLM, FASTmrMLM, FASTmrEMMA, pLARmEB, pKWmEB, ISIS EM-BLASSO and FarmCPU v2.0. As a result, it took 1.43, 1.08, 1.45, 1.53, 3.07, 1.06 and 1.09 hours, respectively (Fig 2d). This indicates that ISIS EM-BLASSO was the fastest one, and FASTmrMLM was equivalent with FarmCPU and faster than mrMLM. The first to fourth experiments were conducted on the first to fourth servers, respectively (File S4).

### Real data analyses in rice, maize and Simmental beef cattle

We re-analyzed the above three datasets in rice [28], maize [29] and Simmental beef cattle [30]. The details can be found in File S3.

The total running times (hours) for the rice dataset were 9.56, 3.37, 11.58, 5.09, 6.13 and 1.06 (hours) for the mrMLM, FASTmrMLM, FASTmrEMMA, pLARmEB, pKWmEB and ISIS EM-BLASSO methods, respectively. Clearly, ISIS EM-BLASSO is the least followed by FASTmrMLM, pLARmEB, pKWmEB, and mrMLM; the FASTmrEMMA is the maximum. The total numbers of QTNs, identified by the above six methods, for grain width in rice were 73, 77, 42, 59, 17, and 31, respectively (**Table S2**). Around these QTNs, some genes were found to be associated with grain width. Among these previously reported genes, two were identified commonly by mrMLM and in Wang et al. [28], and eleven were detected only by the mrMLM (Fig 3a; **Table S3**). In addition, two genes were predicted to be associated with grain width in this study (Fig 3a; **Table S4**).

**Figure 3.**
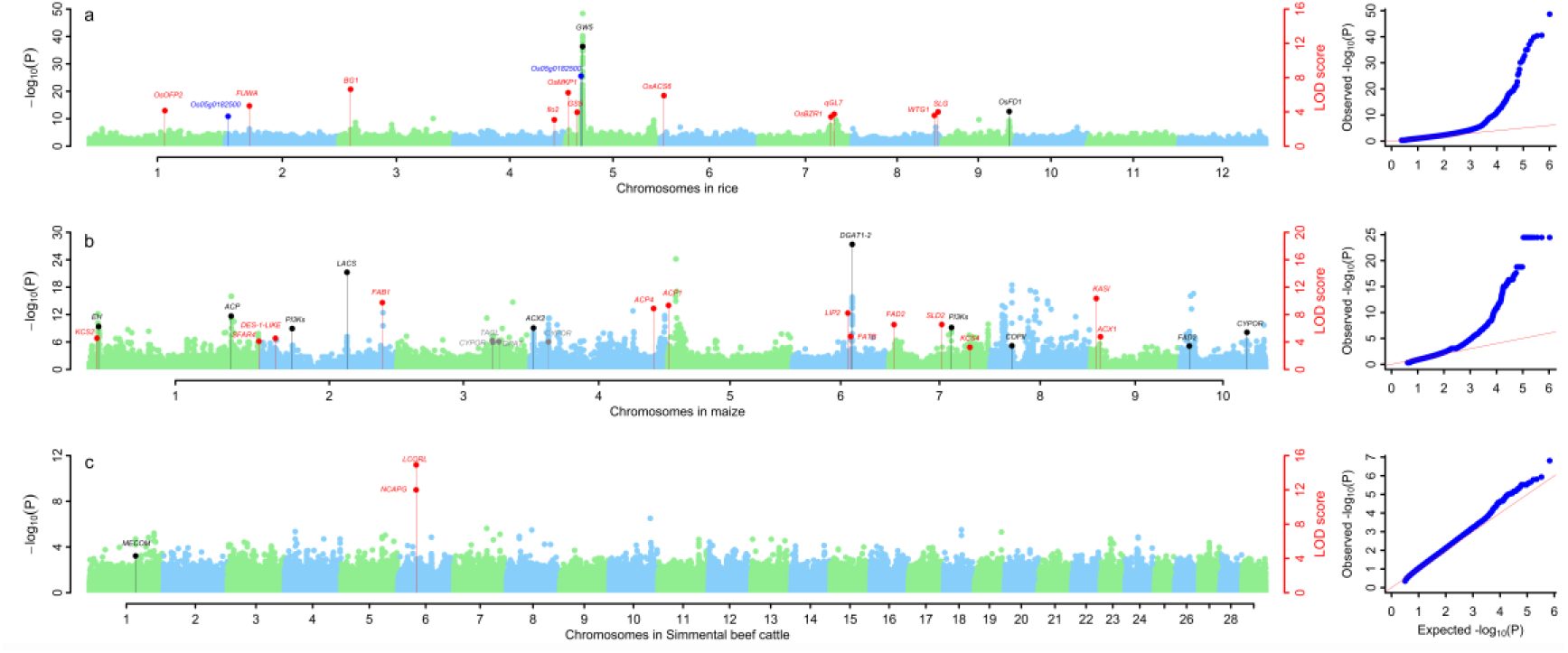
Manhattan (left) and QQ (right) plots in genome-wide association studies using the software mrMLM v4.0. **a.** Grain width in rice [28]. **b.** Oil concentration in maize [29]. **c.** Kidney weight in Simmental beef cattle [30]. The dots with black, red, ash and blue colours were used to indicate the known genes detected commonly by the mrMLM software and in original studies, only by the software mrMLM and only in original studies, and candidate genes around QTNs from the software mrMLM, respectively.

The total numbers of QTNs, detected by the above six methods, for oil concentration in maize were 42, 43, 31, 29, 17 and 6, respectively (**Table S5**). Around these QTNs, some genes were found to be associated with maize oil concentration. Among these previously reported genes, ten were identified commonly by mrMLM and in Li et al. [29], thirteen were detected only by the mrMLM, and four were presented only in Li et al. [29] (Fig 3b) (**Table S6**).

The total numbers of QTNs, identified by the above six methods, for kidney weight in Simmental beef cattle were 4, 55, 167, 117, 8 and 48, respectively (**Table S7**). Around these QTNs, some genes were found to be associated with kidney weight. Among the previously reported genes, *MECOM* was identified commonly by mrMLM and in An et al. [31], and *LCORL* and *NCAPG*, which are very important genes for kidney weight in cattle, were detected only by the mrMLM (Fig 3c; **Table S8**).

## Discussion

To confirm the correctness of our software mrMLM v4.0, the same simulation datasets in Zhang et al. [20] (File S4) were re-analyzed by the above six methods and three current methods (GEMMA [21], FarmCPU [14] and EMMAX [4]). As a result, our six methods were better than the three current methods (Tables S9 to S11; Figs. S3 to S5). The conclusion was also confirmed by the studies in Zhang et al. [32]. As compared with the original packages of our multi-locus GWAS methods, there have been some improvements in the new version. First, the FASTmrMLM algorithm is described for the first time in this study (File S1). Then, the new package is faster in reading datasets and efficient in parallel computing (Fig 2c). Even if the sample size is larger than 2000, FASTmrEMMA is fast as well. This is because at present it is unnecessary for FASTmrEMMA to solve eigenvector at genome scan. Finally, the option for continuous covariates has been set up in order to analyze animal and human GWAS datasets. The new package works well for continuous variables in plant, animal and human GWAS, although the current version doesn’t work for the case-control datasets in human genetics. In addition, we corrected one mistake in the determination of the potentially associated SNPs in the Monte Carlo simulation studies of Zhang et al. [20].

In Zhang et al. [33], several major concerns in GWAS have been discussed, for example, methodological selection, the critical probability value or LOD score, reliable candidate genes, and heritability missing. The details can be found in Zhang et al. [33].

Using mrMLM v4.0, individual parameters may be changed in order to obtain the best results (Files S5 and S6). For example, the number of potentially associated SNPs for each chromosome in Zhang et al. [20] is set at 50, and the search radius in mrMLM [16] and FASTmrMLM [17] is set at 20 kb in real data analysis. In addition, users should understand some parameter settings. For example, the maximum number of CPUs in parallel computation is set at 10. If users want to use more CPU cores, this parameter needs to be modified in the codes. Of course, the accuracy, size and color of the GWAS figures and the critical LOD score line of significant QTNs may be changed as well.

## Conclusion

To popularize our multi-locus GWAS methods, six multi- locus methods have been integrated into one R program, named mrMLM. In this package, three genotypic data formats are available, big dataset can be analyzed at server, parallel computation with multiple CPUs can be performed, and the parameters in the GWAS figures may be set. In addition, the graphical user interface software, mrMLM.GUI v4.0, built upon Shiny, is available as well. Real data analyses confirmed the advantages of our multi-locus GWAS methods.

## Authors’ contributions

YMZ conceived and supervised the study. JG assisted the supervision of the speedup calculation. YWZ, PL, WLR, and YLN wrote the R software. TCL and YMZ proposed the FASTmrMLM algorithm, and YJW and YMZ updated the FASTmrEMMA algorithm. YWZ and PL analysed the datasets. YMZ and YWZ wrote and revised the draft. All authors read and approved the final manuscript.

## Acknowledgements

The work was supported by National Natural Science Foundation of China (31571268, 31871242, U1602261, 31701071 and 21873034), Huazhong Agricultural University Scientific & Technological Self-innovation Foundation (2014RC020), and State Key Laboratory of Cotton Biology Open Fund (CB2019B01). We thank Prof. Jianbing Yan at Huazhong Agricultural University, Prof. Huijiang Gao at Chinese Academy of Agricultural Sciences and Prof. Jim M. Dunwell at University of Reading for providing maize dataset, Simmental beef cattle dataset and the help with improvements to the English text, respectively.

## References

[1] Zhang YM, Mao Y, Xie C, Smith H, Luo L, Xu S. Mapping quantitative trait loci using naturally occurring genetic variance among commercial inbred lines of maize (*Zea mays* L.). Genetics 2005;169:2267–75.

[2] Yu J, Pressoir G, Briggs WH, Vroh Bi I, Yamasaki M, Doebley JF, et al. A unified mixed-model method for association mapping that accounts for multiple levels of relatedness. Nat Genet 2006;38:203–8.

[3] Kang HM, Zaitlen NA, Wade CM, Kirby A, Heckerman D, Daly MJ, et al. Efficient control of population structure in model organism association mapping. Genetics 2008;178:1709–23.

[4] Kang HM, Sul JH, Service SK, Zaitlen NA, Kong SY, Freimer NB, et al. Variance component model to account for sample structure in genome-wide association studies. Nat Genet 2010;42:348–54.

[5] Zhou X, Stephens M. Genome-wide efficient mixed-model analysis for association studies. Nat Genet 2012; 44:821–6

[6] Zhang Z, Ersoz E, Lai CQ, Todhunter RJ, Tiwari HK, Gore MA, et al. Mixed linear model approach adapted for genome-wide association studies. Nat Genet 2010;42:355–60.

[7] Li M, Liu X, Bradbury P, Yu J, Zhang YM, Todhunter RJ, et al. Enrichment of statistical power for genome-wide association studies. BMC Biology 2014;12:73.

[8] Wang Q, Tian F, Pan Y, Buckler ES, Zhang Z. A SUPER powerful method for genome wide association study. PLoS ONE 2014;9:e107684.

[9] Svishcheva GR, Axenovich TI, Belonogova NM, van Duijn CM, Aulchenko YS. Rapid variance components-based method for whole-genome association analysis. Nat Genet 2012;44:1166–70

[10] Lippert C, Listgarten J, Liu Y, Kadie CM, Davidson RI, Heckerman D. FaST linear mixed models for genome-wide association studies. Nat Methods 2011;8:833–5.

[11] Listgarten J, Lippert C, Kadie CM, Davidson RI, Eskin E, Heckerman D. Improved linear mixed models for genome-wide association studies. Nat Methods 2012;9:525–6.

[12] Loh PR, Tucker G, Bulik-Sullivan BK, Vilhjálmsson BJ, Finucane HK, Salem RM, et al. Efficient Bayesian mixed-model analysis increases association power in large cohorts. Nat Genet 2015;47:284–90.

[13] Segura V, Vilhjálmsson B J, Platt A, Korte A, Seren Ü, Long Q, et al. An efficient multi-locus mixed-model approach for genome-wide association studies in structured populations. Nat Genet 2012;44:825–30.

[14] Liu X, Huang M, Fan B, Bucklers ES, Zhang Z. Iterative usage of fixed and random effect models for powerful and efficient genome-wide association studies. PLoS Genet 2016;12:e1005767.

[15] Goddard ME, Wray NR, Verbyla K, Visscher PM. Estimating effects and making predictions from genome-wide marker data. Stat Sci 2009; 24:517–29.

[16] Wang SB, Feng JY, Ren WL, Huang B, Zhou L, Wen YJ, et al. Improving power and accuracy of genome-wide association studies via a multi-locus mixed linear model methodology. Sci Rep 2016;6:19444.

[17] Tamba CL, Zhang YM. A fast mrMLM algorithm for multi-locus genome-wide association studies. bioRxiv, 2018. doi: 10.1101/341784.

[18] Wen YJ, Zhang H, Ni YL, Huang B, Zhang J, Feng JY, et al. Methodological implementation of mixed linear models in multi-locus genome-wide association studies. Brief Bioinform 2018;19:700–12.

[19] Tamba CL, Ni YL, Zhang YM. Iterative sure independence screening EM Bayesian LASSO algorithm for multi-locus genome-wide association studies. PLoS Comput Biol 2017;13:e1005357.

[20] Zhang J, Feng JY, Ni YL, Wen YJ, Niu Y, Tamba CL, et al. pLARmEB: integration of least angle regression with empirical Bayes for multilocus genome-wide association studies. Heredity 2017;118:517–24.

[21] Ren WL, Wen YJ, Dunwell JM, Zhang YM. pKWmEB: Integration of Kruskal-Wallis test with empirical Bayes under polygenic background control for multi-locus genome-wide association study. Heredity 2018; 120:208–18.

[22] Purcell S, Neale B, Todd-Brown K, Thomas L, Ferreira MAR, Bender D, et al. PLINK: a tool set for whole-genome association and population-based linkage analyses. Am J Hum Genet 2007;81:559–75.

[23] Bradbury PJ, Zhang Z, Kroon DE, Casstevens TM, Ramdoss Y, Buckler ES. TASSEL: software for association mapping of complex traits in diverse samples. Bioinformatics 2007;23:2633–35.

[24] Lipka AE, Tian F, Wang Q, Peiffer J, Li M, Bradbury PJ, et al. GAPIT: genome association and prediction integrated tool. Bioinformatics 2012;28:2397–99.

[25] Tang Y, Liu X, Wang J, Li M, Wang Q, Tian F, et al. GAPIT Version 2: An enhanced integrated tool for genomic association and prediction. Plant Genome 2016;9:doi:10.3835/plantgenome2015.11.0120

[26] Pritchard JK, Stephens M, Donnelly P. Inference of population structure using multilocus genotype data. Genetics 2000;155:945–59.

[27] Raj A, Stephens M, Pritchard JK. fastSTRUCTURE: variational inference of population structure in large SNP data sets. Genetics 2014;197:573–89.

[28] Wang W, Mauleon R, Hu Z, Chebotarov D, Tai S, Wu Z, et al. Genomic variation in 3,010 diverse accessions of Asian cultivated rice. Nature 2018; 557: 43–49.

[29] Li H, Peng Z, Yang X, Wang W, Fu J, Wang J, et al. Genome-wide association study dissects the genetic architecture of oil biosynthesis in maize kernels. Nat Genetics 2013;45:43–50.

[30] Zhu B, Zhu M, Jiang J, Niu H, Wang Y, Wu Y, et al. The impact of variable degrees of freedom and scale parameters in Bayesian methods for genomic prediction in Chinese Simmental beef cattle. PLoS ONE 2016; 11(5): e0154118.

[31] An B, Xia J, Chang T, Wang X, Miao J, Xu L, et al. Genome-wide association study identifies loci and candidate genes for internal organ weights in Simmental beef cattle. Physiol Genomics 2018;50:523–31.

[32] Zhang YM, Jia Z, Dunwell JM (eds). The applications of new multi-locus GWAS methodologies in the genetic dissection of complex traits. Lausanne: Frontiers Media. 2019. doi: 10.3389/978-2-88945-834-9

[33] Zhang YM, Jia Z, Dunwell JM. Editorial: The applications of new multi-locus GWAS methodologies in the genetic dissection of complex traits. Front Plant Sci 2019;10:100.

