## Supplementary Materials for "mrMLM v4.0: An R Platform for Multi-locus Genome-wide Association Studies"

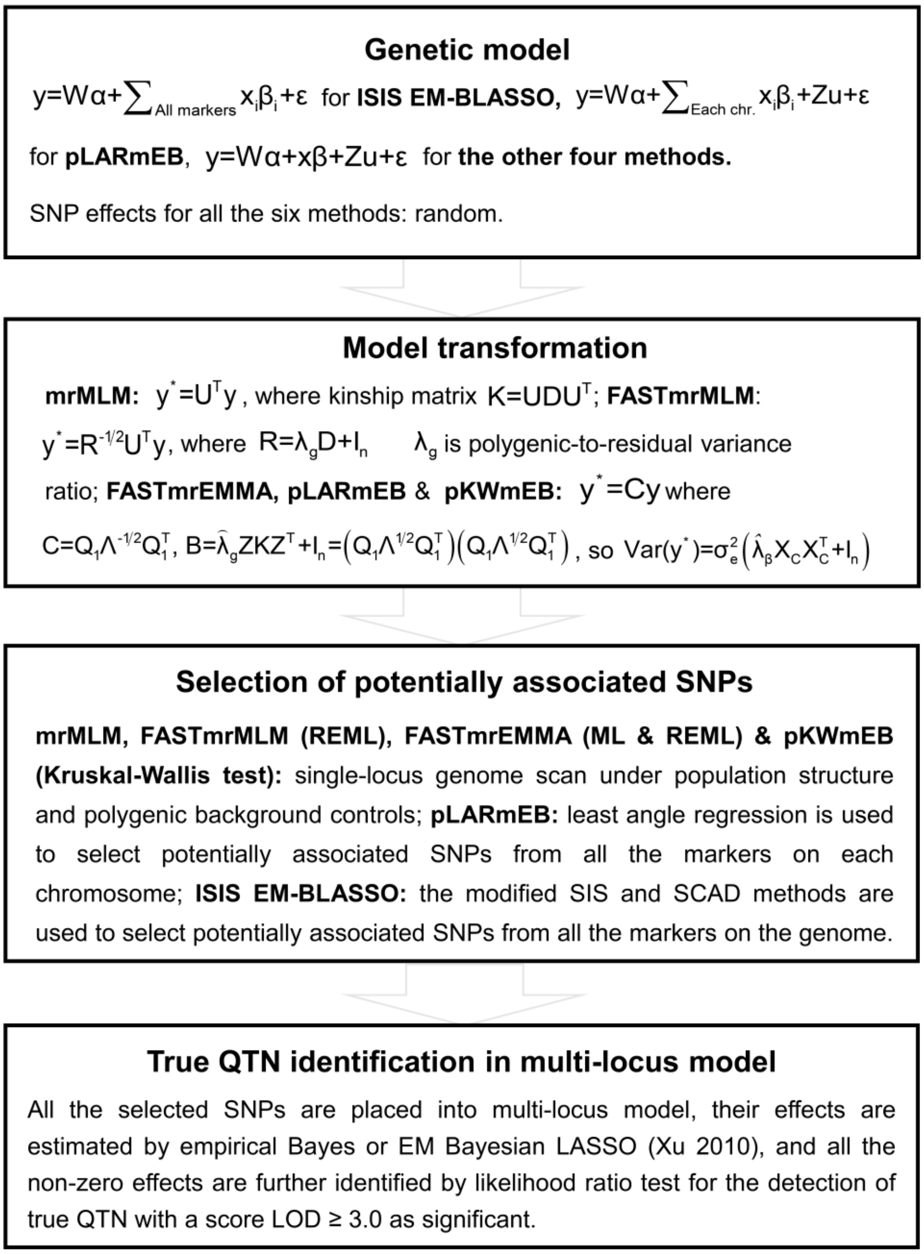


**Figure S1 Methodological comparison in the software package mrMLM v4.0**

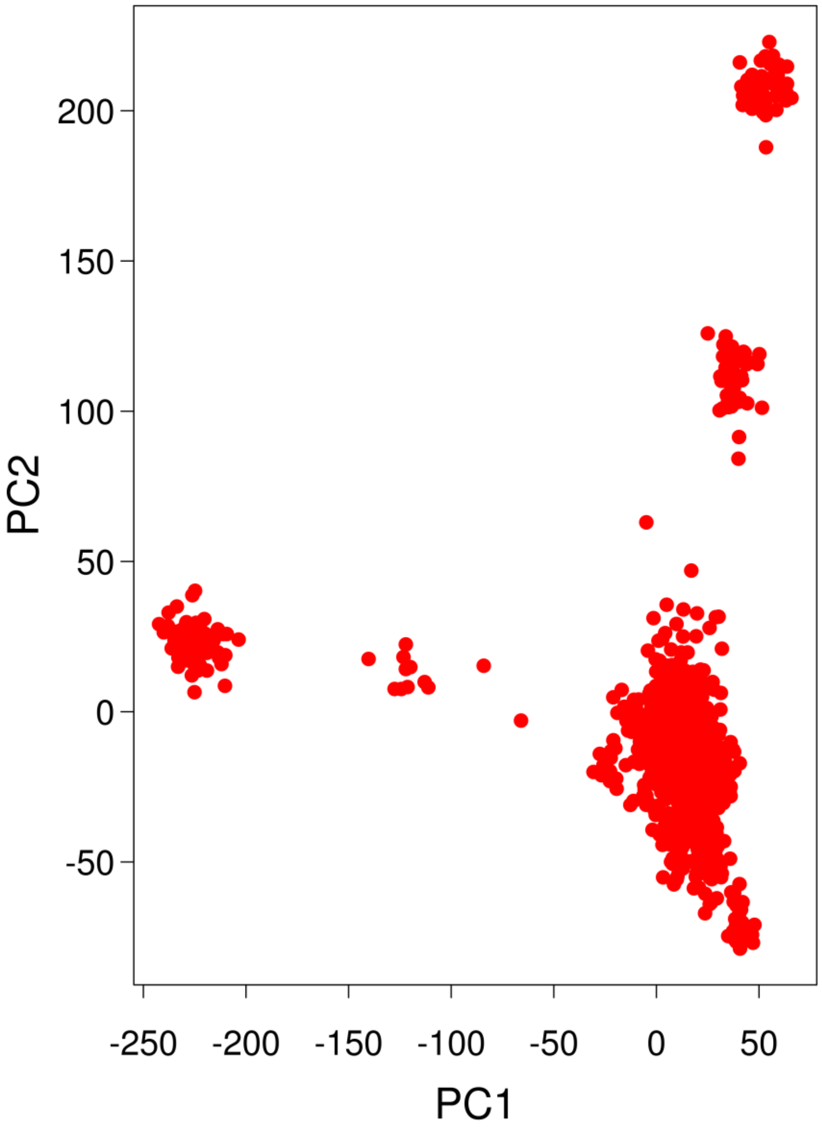


**Figure S2 Principal component analysis for the genotypes in Simmental beef cattle (**The dataset was derived from Zhu et al [30]**)**

#### Supplementary material A. The FASTmrMLM algorithm

##### Genetic model

We consider a mixed linear regression model,

$$\mathbf{y} = \mathbf{X}\boldsymbol{\alpha} + \mathbf{Z}_i\beta_i + \boldsymbol{\phi} + \boldsymbol{\varepsilon} \tag{A1}$$

where $\mathbf{y}$ is an $n\times 1$ phenotypic vector of quantitative trait for $n$ individuals, $\mathbf{X}$ is an $n\times q$ incident matrix of fixed effects $\boldsymbol{\alpha}$ including the overall mean, $\mathbf{Z_i}$ is an $n\times 1$ vector of the $i$th SNP, $\beta_i$ is a random effect of the $i$th marker, it is assumed to be a normal distribution with zero mean and each marker prior variance $\sigma_i^2$, $\boldsymbol{\phi} \sim \text{MVN}\left(\mathbf{0}, \mathbf{K}\sigma_g^2\right)$ is the polygenic effect with a multivariate normal distribution with zero mean and variance $\sigma_g^2$ described by a kinship matrix $\mathbf{K}$, and $\boldsymbol{\varepsilon}\sim\text{MVN}\left(\mathbf{0},\mathbf{I}\sigma_e^2\right)$ is residual error where $\mathbf{I}$ is an $n\times n$ identity matrix and $\sigma_e^2$ is residual variance. In this study, the kinship matrix $\mathbf{K}$ is marker inferred kinship matrix defined as $\mathbf{K} = \sum_{i=1}^{m}\mathbf{Z}_i\mathbf{Z}_i' \Big/ m$.

From equation (A1) we have that

$$\begin{aligned} &\text{E}(\mathbf{y}) = \mathbf{X}\boldsymbol{\alpha} \\ &\text{Var}(\mathbf{y}) = \mathbf{Z}_i\mathbf{Z}_i'\sigma_i^2 + \mathbf{K}\sigma_g^2 + \mathbf{I}\sigma_e^2 = \sigma_e^2\left(\mathbf{Z}_i\mathbf{Z}_i'\omega_i + \mathbf{K}\omega_g + \mathbf{I}\right) = \sigma_e^2\mathbf{H} \end{aligned} \tag{A2}$$

where $\omega_i = \sigma_i^2/\sigma_e^2$ and $\omega_g = \sigma_g^2/\sigma_e^2$ are the variance ratios and $\mathbf{H} = \mathbf{Z}_i\mathbf{Z}_i'\omega_i + \mathbf{K}\omega_g + \mathbf{I}$. In this model, we have three random effects: the $i$th marker effect, polygenic effect, and residual error. Under the pure polygenic model, we have that $\text{E}(\mathbf{y}) = \mathbf{X}\boldsymbol{\alpha}$ and $\text{Var}(\mathbf{y}) = \sigma_e^2\left(\mathbf{K}\omega_g + \mathbf{I}\right)$. We pre-estimate the value of $\omega_g = \sigma_g^2/\sigma_e^2$ under the pure polygenic model and fix it when testing each SNP effect in the genome-wide scanning. Using spectral decomposition, we can find a diagonal matrix $\mathbf{D} = \text{diag}\left(\delta_1,\cdots,\delta_m\right)$ and a matrix $\mathbf{U}$ such that $\mathbf{K}=\mathbf{UDU}'$. Notice that spectral decomposition can be performed on $\mathbf{K}$ since as defined it is a square, symmetric matrix. Transforming $\mathbf{y}$ in Equation (A1) by multiplying by $\mathbf{U}'$, we have,

$$\mathbf{U}'\mathbf{y}=\mathbf{U}'\mathbf{X}\boldsymbol{\alpha}+\mathbf{U}'\mathbf{Z}_i\beta_i+\mathbf{U}'(\boldsymbol{\phi}+\boldsymbol{\varepsilon}) \tag{A3}$$

Let $\mathbf{y}^*=\mathbf{U}'\mathbf{y},\mathbf{X}^*=\mathbf{U}'\mathbf{X},\mathbf{Z}_i^*=\mathbf{U}'\mathbf{Z}_i$, then Equation (3) becomes

$$\mathbf{y}^*=\mathbf{X}^*\boldsymbol{\alpha}+\mathbf{Z}_i^*\beta_i+\mathbf{U}'(\boldsymbol{\phi}+\boldsymbol{\varepsilon})$$

It follows that,

$$\begin{aligned}\mathrm{Var}(\mathbf{y}^*)&=\mathbf{Z}_i^*\mathbf{Z}_i^{*\prime}\sigma_i^2+\mathbf{U}'\mathbf{KU}\sigma_g^2+\mathbf{U}'\mathbf{IU}\sigma_e^2=\sigma_e^2\left(\mathbf{Z}_i^*\mathbf{Z}_i^{*\prime}\omega_i+\mathbf{D}\hat{\omega}_g+\mathbf{I}\right)\\&=\sigma_e^2\left(\mathbf{Z}_i^*\mathbf{Z}_i^{*\prime}\omega_i+\mathrm{diag}\left(\delta_1\hat{\omega}_g+1,\delta_2\hat{\omega}_g+1,\cdots,\delta_m\hat{\omega}_g+1\right)\right)\end{aligned} \tag{A4}$$

For simplicity, let $\mathbf{D}_1=\mathrm{diag}\left(\delta_1\hat{\omega}_g+1,\delta_2\hat{\omega}_g+1,\cdots,\delta_m\hat{\omega}_g+1\right)$. $\hat{\omega}_g$ is fixed, and $\mathbf{D}_1$, as defined, is a positive semi-definite matrix. Therefore, we can obtain $\mathbf{D}_1^{1/2}$. Further transforming $\mathbf{y}^*$ by multiplying equation (A3) through by $\mathbf{D}_1^{-1/2}$ and letting $\mathbf{y}_c=\mathbf{D}_1^{-1/2}\mathbf{y}^*,\mathbf{X}_c=\mathbf{D}_1^{-1/2}\mathbf{X}^*,\mathbf{Z}_c=\mathbf{D}_1^{-1/2}\mathbf{Z}_i^*$ and $(\boldsymbol{\phi}+\boldsymbol{\varepsilon})_c=\mathbf{D}_1^{-1/2}\mathbf{U}'(\boldsymbol{\phi}+\boldsymbol{\varepsilon})$. We have that,

$$\begin{aligned}\mathbf{D}_1^{-1/2}\mathbf{y}^*&=\mathbf{D}_1^{-1/2}\mathbf{X}^*\boldsymbol{\alpha}+\mathbf{D}_1^{-1/2}\mathbf{Z}_i^*\beta_i+\mathbf{D}_1^{-1/2}\mathbf{U}'(\boldsymbol{\phi}+\boldsymbol{\varepsilon})\\\mathbf{y}_c&=\mathbf{X}_c\boldsymbol{\alpha}+\mathbf{Z}_{ci}\beta_i+(\boldsymbol{\phi}+\boldsymbol{\varepsilon})_c\end{aligned} \tag{A5}$$

Notice that the transformation in Equation (A5) is equivalent to multiplying the original $\mathbf{y}$ by $\mathbf{U}'\mathbf{D}_1^{-\frac{1}{2}}$ i.e. $\mathbf{y}_c=\mathbf{U}'\mathbf{D}_1^{-1/2}(\mathbf{y})$. Now,

$$\begin{aligned}&\mathrm{E}(\mathbf{y}_c)=\mathbf{X}_c\boldsymbol{\alpha}\\&\mathrm{Var}(\mathbf{y}_c)=\sigma_e^2\left(\mathbf{Z}_{ci}\mathbf{Z}_{ci}'\omega_i+\mathbf{I}\right)=\sigma_e^2(\mathbf{A}+\mathbf{B})=\sigma_e^2\mathbf{H}\end{aligned} \tag{A6}$$

where $\mathbf{A}=\mathbf{Z}_{ci}\mathbf{Z}_{ci}'\omega_i$, $\mathbf{B}=\mathbf{I}$, and $\mathbf{H}=\mathbf{A}+\mathbf{B}$. Thus, the distribution of our transformed data vector $\mathbf{y}_c$ is normal with the mean $\mathbf{X}_c\boldsymbol{\alpha}$ and variance-covariance matrix $\sigma_e^2\mathbf{H}$. The parameters to be estimated are $\sigma_e^2$ and $\omega_i$ for $i=1,2,\cdots,m$.

**Parameter estimation**

The profiled residual log likelihood (REML) of $\mathbf{y}^*$ after absorbing other terms in the constant term $C$ is,

$$\ell_R = C - \frac{1}{2}\left\{(n-q)\log\sigma_e^2 + \log|\mathbf{H}| + \log\left|\mathbf{X}_c{}'\mathbf{H}^{-1}\mathbf{X}_c\right| + \frac{\mathbf{y}_c{}'\mathbf{P}\mathbf{y}_c}{\sigma_e^2}\right\} \quad (A7)$$

where, $\mathbf{P} = \mathbf{H}^{-1} - \mathbf{H}^{-1}\mathbf{X}_c\left(\mathbf{X}_c{}'\mathbf{H}^{-1}\mathbf{X}_c\right)^{-1}\mathbf{X}_c{}'\mathbf{H}^{-1}$, and $q = \text{rank}(\mathbf{X}_c)$. We differentiate equation (A7) to obtain REML estimates as shown below.

**Estimation of residual variance** $\sigma_e^2$**:** We have

$$\frac{\partial\ell_R}{\partial\sigma_e^2} = -\frac{1}{2}\left[\frac{n-q}{\sigma_e^2} - \frac{\mathbf{y}_c{}'\mathbf{P}\mathbf{y}_c}{\left(\sigma_e^2\right)^2}\right] \quad \Rightarrow \quad \hat{\sigma}_e^2 = \frac{\mathbf{y}_c{}'\mathbf{P}\mathbf{y}_c}{n-q}$$

This can be simplified to

$$\hat{\sigma}_e^2 = \frac{\mathbf{y}_c{}'\mathbf{P}\mathbf{y}_c}{n-q} = \frac{1}{n-q}\left[\left(\mathbf{y}_c{}'\mathbf{H}^{-1}\mathbf{y}_c\right) - \left(\mathbf{y}_c{}'\mathbf{H}^{-1}\mathbf{X}_c\right)\left(\mathbf{X}_c{}'\mathbf{H}^{-1}\mathbf{X}_c\right)^{-1}\left(\mathbf{X}_c{}'\mathbf{H}^{-1}\mathbf{y}_c\right)\right] \quad (A8)$$

**Estimation of the variance ratio** $\omega_i$**:** We have that

$$\frac{\partial\ell_R}{\partial\omega_i} = -\frac{1}{2}\left[\text{tr}(\mathbf{P}\mathbf{Z}_{ci}\mathbf{Z}_{ci}{}') - \frac{1}{\hat{\sigma}_e^2}\mathbf{y}_c{}'\mathbf{P}\mathbf{Z}_{ci}\mathbf{Z}'_{ci}\mathbf{P}\mathbf{y}_c\right] \quad (A9)$$

The second derivative of the residual log likelihood function with respect to $\omega_i$ is given below, and it is used to obtain the Hessian matrix.

$$\frac{\partial^2\ell_R}{\partial\omega_i^2} = -\frac{1}{2}\left\{-\text{tr}\left(\mathbf{P}\mathbf{Z}_{ci}\mathbf{Z}_{ci}{}'\mathbf{P}\mathbf{Z}_{ci}\mathbf{Z}_{ci}{}'\right) - (n-q)\frac{2\left(\mathbf{y}_c{}'\mathbf{P}\mathbf{Z}_{ci}\mathbf{Z}_{ci}{}'\mathbf{P}\mathbf{Z}_{ci}\mathbf{Z}_{ci}{}'\mathbf{P}\mathbf{y}_c\right)\left(\mathbf{y}_c{}'\mathbf{P}\mathbf{y}_c\right) - \left(\mathbf{y}_c{}'\mathbf{P}\mathbf{Z}_{ci}\mathbf{Z}_{ci}{}'\mathbf{P}\mathbf{y}_c\right)^2}{\left(\mathbf{y}_c{}'\mathbf{P}\mathbf{y}_c\right)^2}\right\} \quad (A10)$$

Evaluation of REML equation and its derivatives require $\mathbf{H}^{-1}$ and $|\mathbf{H}|$. $\mathbf{H}$ is the sum of the above two matrices. The inverse and determinant of $\mathbf{B}$ can easily be computed because it is an identity matrix, and $\mathbf{A}$ is a matrix of rank 1. In mrMLM of Wang et al. [1], $\mathbf{H}$ is also the sum of two matrices but in this new algorithm (FASTmrMLM) we have simplified the $\mathbf{H}$ further to be the sum of an identity matrix and a matrix of rank one. Therefore,

$$\mathbf{H}^{-1} = \mathbf{B}^{-1} - \frac{1}{1+g}\mathbf{B}^{-1}\mathbf{A}\mathbf{B}^{-1} = \mathbf{I} - \frac{1}{1+g}\mathbf{A} = \mathbf{I} - \frac{\omega_i}{1+g}\mathbf{Z}_{ci}\mathbf{Z}_{ci}' \tag{A11}$$

and $|\mathbf{H}| = (1+g)|\mathbf{B}| = 1+g$, where $g = \mathrm{tr}(\mathbf{A}\mathbf{B}^{-1}) = \mathrm{tr}(\mathbf{A}) = \omega_i\,\mathrm{tr}\left(\mathbf{Z}_{ci}\mathbf{Z}_{ci}'\right)$ [2]. REML requires many quadratic terms in the form $\boldsymbol{\eta}'\mathbf{H}^{-1}\boldsymbol{\tau}$, which can be expressed as

$$\boldsymbol{\eta}'\mathbf{H}^{-1}\boldsymbol{\tau} = \boldsymbol{\eta}'\boldsymbol{\tau} - \frac{\omega_i}{1+g}\boldsymbol{\eta}'\mathbf{Z}_{ci}\mathbf{Z}_{ci}'\boldsymbol{\tau} \tag{A12}$$

where $\boldsymbol{\eta}'$ and $\boldsymbol{\tau}$ can be any vectors or matrices with $\mathbf{H}^{-1}$ such as $\mathbf{X}_c'\mathbf{H}^{-1}\mathbf{X}_c$, $\mathbf{y}_c'\mathbf{P}\mathbf{y}_c$ or $\mathbf{X}_c'\mathbf{P}\mathbf{y}_c$. Note that $\boldsymbol{\eta}'\boldsymbol{\tau} = \sum_{j=1}^{n}\eta_j'\tau_j$, where $\eta_j$ corresponds to the *j*th row (element) of the matrix (vector) $\boldsymbol{\eta}$ and $\tau_j$ corresponds to the *j*th row (element) of the matrix (vector) $\boldsymbol{\tau}$ for $j = 1, 2, \cdots, n$. Thus, the value in Equation (A8) above can easily be estimated because with the help of Equation (A12) each term in the bracket in Equation (A8) is in the form $\boldsymbol{\eta}'\mathbf{H}^{-1}\boldsymbol{\tau}$. For example, $\mathbf{y}_c\mathbf{H}^{-1}\mathbf{y}_c = \mathbf{y}_c'\mathbf{y}_c - \frac{\omega_i}{1+g}\mathbf{y}_c'\mathbf{Z}_{ci}\mathbf{Z}_{ci}'\mathbf{y}_c$. Also, the gradient function / score function in Equation (A9) and Hessian matrix in Equation (A10) can also be expressed in the form $\boldsymbol{\eta}'\mathbf{H}^{-1}\boldsymbol{\tau}$. With these simplifications, the computation of the variance ratio $\omega_i$ for the *i*th SNP is less computationally intensive via the Newton-Raphson method. We estimate the variance ratio $\omega_i$ for the *i*th SNP by equating the gradient functions to be zero using the Newton-Raphson technique. With our simplifications,

$$\begin{aligned}\frac{\partial \ell_R}{\partial \omega_i} &= -\frac{1}{2}\left[\mathrm{tr}(\mathbf{P}\mathbf{Z}_{ci}\mathbf{Z}_{ci}') - \frac{1}{\hat{\sigma}_e^2}\mathbf{y}_c'\mathbf{P}\mathbf{Z}_{ci}\mathbf{Z}'_{ci}\mathbf{P}\mathbf{y}_c\right] \\ &= -\frac{1}{2}\left\{\mathrm{tr}\left(\mathbf{Z}_{ci}\mathbf{H}^{-1}\mathbf{Z}_{ci}' - \mathbf{Z}_{ci}\mathbf{H}^{-1}\mathbf{X}_c\left(\mathbf{X}_c'\mathbf{H}^{-1}\mathbf{X}_c\right)^{-1}\mathbf{X}_c'\mathbf{H}^{-1\prime}\mathbf{Z}_{ci}'\right) - \frac{1}{\hat{\sigma}_e^2}\left[\left(\left(\mathbf{y}_c'\mathbf{H}^{-1}\mathbf{Z}_{ci}\right) - \left(\mathbf{y}_c'\mathbf{H}^{-1}\mathbf{X}_c\right)\left(\mathbf{X}_c'\mathbf{H}^{-1}\mathbf{X}_c\right)^{-1}\left(\mathbf{X}_c'\mathbf{H}^{-1}\mathbf{Z}_{ci}\right)\right)\right.\right. \\ &\quad \left.\left.\times\left(\left(\mathbf{Z}'_{ci}\mathbf{H}^{-1}\mathbf{y}_c\right) - \left(\mathbf{Z}'_{ci}\mathbf{H}^{-1}\mathbf{X}_c\right)\left(\mathbf{X}_c'\mathbf{H}^{-1}\mathbf{X}_c\right)^{-1}\left(\mathbf{X}_c'\mathbf{H}^{-1}\mathbf{y}_c\right)\right)\right]\right\}\end{aligned} \tag{A13}$$

which is in the form $\boldsymbol{\eta}'\mathbf{H}^{-1}\boldsymbol{\tau}$, and therefore its computation is so fast. We can also express the second derivative expression in Equation (A10) in the form $\boldsymbol{\eta}'\mathbf{H}^{-1}\boldsymbol{\tau}$:

$$\frac{\partial^2 \ell_R}{\partial \omega_i^2} = -\frac{1}{2}\left\{-\mathrm{tr}(\mathbf{Z}_{ci}\mathbf{P}\mathbf{Z}_{ci}'\mathbf{Z}_{ci}\mathbf{P}\mathbf{Z}_{ci}') - (n-q)\frac{2\left(\mathbf{y}_c'\mathbf{P}\mathbf{Z}_{ci}\mathbf{Z}_{ci}'\mathbf{P}\mathbf{Z}_{ci}\mathbf{Z}_{ci}'\mathbf{P}\mathbf{y}_c\right)\left(\mathbf{y}_c'\mathbf{P}\mathbf{y}_c\right) - \left(\mathbf{y}_c'\mathbf{P}\mathbf{Z}_{ci}\mathbf{Z}_{ci}'\mathbf{P}\mathbf{y}_c\right)^2}{\left(\mathbf{y}_c'\mathbf{P}\mathbf{y}_c\right)^2}\right\}.$$

With these simplifications, the Newton-Raphson will converge smoothly to the estimate value of variance ratio $\omega_i$ for the *i*th SNP. The gradient function and the Hessian matrix are in a simplified form. Therefore, the computations in each simulation run are fast. We have implemented this algorithm in R software.

**Empirical Bayes estimate of** $\beta_i$ **:** The joint distribution of $\mathbf{y}$ and $\beta_i$ is a multivariate normal distribution

$$\begin{pmatrix}\mathbf{y}\\ \beta_i\end{pmatrix} \sim \mathrm{MVN}\left(\begin{bmatrix}\mathbf{X}\boldsymbol{\alpha}\\ 0\end{bmatrix}, \begin{bmatrix}\mathbf{Z}_i\mathbf{Z}_i'\omega_i + \mathbf{K}\omega_g + \mathbf{I} & \mathbf{Z}_i\sigma_i^2\\ \mathbf{Z}_i'\sigma_i^2 & \sigma_i^2\end{bmatrix}\right)$$

The conditional distribution of $\beta_i$ given $\mathbf{y}$ is

$$\beta_i \mid \mathbf{y} \sim \mathrm{N}\left[\mathbf{Z}_i'\hat{\sigma}_i^2\left(\mathbf{Z}_i\mathbf{Z}_i'\hat{\sigma}_i^2 + \mathbf{K}\hat{\sigma}_g^2 + \mathbf{I}\hat{\sigma}_e^2\right)^{-1}(\mathbf{y} - \mathbf{X}\hat{\boldsymbol{\alpha}}),\ \hat{\sigma}_i^2 - \hat{\sigma}_i^2\mathbf{Z}_i'\left(\mathbf{Z}_i\mathbf{Z}_i'\hat{\sigma}_i^2 + \mathbf{K}\hat{\sigma}_g^2 + \mathbf{I}\hat{\sigma}_e^2\right)^{-1}\mathbf{Z}_i\hat{\sigma}_i^2\right]$$

From a Bayesian analysis point of view, the conditional mean of $\beta_i$ given $\mathbf{y}$ is an empirical Bayes estimate of $\beta_i$. Based on this framework, we obtain the Wald test statistic $\frac{\left(\mathrm{E}(\beta_i \mid \mathbf{y})\right)^2}{\mathrm{Var}(\beta_i \mid \mathbf{y})} \sim \chi_1^2$ (Chi-square test with 1 degree of freedom) and using this distribution we obtain P-value for each marker effect. We test each marker effect at 1% level of significance. We do not perform multiple test correction because we intend to include markers that pass this initial test in a multi-locus model.

**Detection of true QTNs in multi-locus model**

If the number of markers passing the 1% level of significance test is more than $n$, we invoke the LARS algorithm [3] to select the $n-1$ variables that are most likely associated with the quantitative trait of interest. LARS is a flexible method for variable selection, which is conducted in lars package (http://cran.r-project.org/web/packages/lars/) in R language. The $n-1$ markers are then included in a multi-locus model. Note that if the number of markers passing the initial test is less than $n$, we skip the LARS step and proceed to include all the selected markers in a multi-locus model. We compared various multi-locus methods: SCAD [4], adaptive Lasso [5] and EM-Empirical Bayes [6]. EM-Empirical Bayes has the highest statistical power and accuracy of the estimated marker effects. EM-Empirical Bayes is a random model method given as,

$$\mathbf{y} = \mathbf{X}\boldsymbol{\alpha} + \sum_{i=1}^{s} \mathbf{Z}_i \beta_i + \boldsymbol{\varepsilon} \tag{A14}$$

where $\mathbf{y}$, $\mathbf{X}$ and $\boldsymbol{\alpha}$ are the same as in model in Equation (A1), $s$ is the number of potentially associated markers selected from the first step in FASTmrMLM, $\mathbf{Z}_i$ and $\beta_i$ are $n \times 1$ incident vector and the random effect of the $i$th SNP, respectively. The polygenic variance is not included in the model because the model included all the potentially associated QTNs. We assume a normal prior for $\beta_i$, $\beta_i \mid \sigma_i^2 \sim N\left(0, \sigma_i^2\right)$ and a scaled $\chi^2$ prior for $\sigma_i^2$, $f\left(\sigma_i^2 \mid \tau, \omega\right) \propto \left(\sigma_i^2\right)^{-\frac{1}{2}(\tau+2)} \exp\left(-\frac{\omega}{2\sigma_i^2}\right)$ and we set $(\tau, \omega) = (0, 0)$, which is Jeffrey's prior $f\left(\sigma_i^2 \mid \tau, \omega\right) = \sigma_i^2/2$ [6]. The procedure for parameter estimation in EM-Empirical Bayes is as follows:

a) Initial step: We set initial values as

$$\begin{aligned} &\sigma_i^2 = 1, \text{ for } i = 1, 2, \cdots, s \\ &\boldsymbol{\alpha} = \left(\mathbf{X}'\mathbf{X}\right)^{-1} \mathbf{X}'\mathbf{y} \\ &\sigma_e^2 = \frac{1}{2n} (\mathbf{y} - \mathbf{X}\boldsymbol{\alpha})' (\mathbf{y} - \mathbf{X}\boldsymbol{\alpha}) \end{aligned} \tag{A15}$$

b) E-step: QTN effect can be predicted by

$$\mathrm{E}(\beta_i) = \sigma_i^2 \mathbf{Z}_i' \mathbf{V}^{-1} (\mathbf{y} - \mathbf{X}\boldsymbol{\alpha}) \tag{A16}$$

where $\mathbf{V} = \sum_{i=1}^{s} \mathbf{Z}_i \mathbf{Z}_i' \sigma_i^2 + \mathbf{I}\sigma_e^2$ .

c) M-step: To update parameters $\sigma_i^2$, $\boldsymbol{\alpha}$ and $\sigma_e^2$

$$\begin{aligned} \sigma_i^2 &= \frac{\mathrm{E}(\beta_i'\beta_i) + \omega}{\tau + 3} \\ \boldsymbol{\alpha} &= \left(\mathbf{X}'\mathbf{V}^{-1}\mathbf{X}\right)^{-1} \mathbf{X}'\mathbf{V}^{-1}\mathbf{y} \\ \hat{\sigma}_e^2 &= \frac{1}{n}(\mathbf{y} - \mathbf{X}\boldsymbol{\alpha})' \left\{ (\mathbf{y} - \mathbf{X}\boldsymbol{\alpha}) - \sum_{i=1}^{s} \mathbf{Z}_i \mathrm{E}(\beta_i) \right\} \end{aligned} \tag{A17}$$

where $\mathrm{E}(\beta_i'\beta_i) = \mathrm{E}(\beta_i')\mathrm{E}(\beta_i) + \mathrm{tr}\left[\mathrm{Var}(\beta_i)\right]$, $\mathrm{Var}(\beta_i) = \mathbf{I}\sigma_i^2 - \sigma_i^2 \mathbf{Z}_i' \mathbf{V}^{-1} \mathbf{Z}_i \sigma_i^2$ and $(\tau, \omega) = (0, 0)$.

We repeat E-step and M-step until convergence is satisfied. We select all SNPs with a score $\mathrm{LOD} \geq 3$ (log of odds) and regard them as significant. We term our algorithm as a fast multi-locus random-SNP-effect mixed linear model (FASTmrMLM).

#### References


1 Wang SB, Feng JY, Ren WL, Huang B, Zhou L, Wen YJ, et al. Improving power and accuracy of genome-wide association studies via a multi-locus mixed linear model methodology. **Sci Rep** 2016;**6:**19444.

2 Miller K. On the inverse of the sum of matrices. **Mathematics Magazine** 1981;**54**:67–72.

3 Efron B, Hastie T, Johnstone I, Robert T. Least angle regression. **Ann Stat** 2004;**32**:407–99.

4 Fan J, Li R. Variable selection via nonconcave penalized likelihood and its oracle properties. **Journal of the American Statistical Association** 2001; **96** (456): 1348–60.

5 Zou H. The adaptive lasso and its oracle properties. **J Am Stat Assoc** 2006; 101(476): 1418–29.

6 Xu S. An expectation-maximization algorithm for the Lasso estimation of quantitative trait locus effects. **Heredity** 2010;**105**:483–94.

#### Supplementary material B. The GWAS methodologies and software packages

##### *Multi-locus random-SNP-effect mixed linear model (mrMLM) and FASTmrMLM*

In the mrMLM of Wang et al. [1], three techniques were used to reduce the running time in the first stage. First, the polygenic-to-residual variance ratio $\lambda$ was estimated under a pure polygenic model (the null model) and then treated as a constant ($\hat{\lambda}$) in single-marker association in genome-wide scans. Then, the residual variance at the single-marker association was estimated together with fixed effects. More importantly, eigen decomposition for kinship matrix **K** was carried out so that $\mathbf{K} = \mathbf{UDU}^T$ , where $\mathbf{D} = \mathrm{diag}\{\delta_1, \cdots, \delta_n\}$ is a diagonal matrix for the eigenvalues and **U** is an *n* × *n* matrix for the eigenvectors. Therefore, the special structure of $\mathrm{R}_k$ in restricted likelihood function allows us to implement the Woodbury matrix identities for calculating $|\mathrm{R}_k|$ and $\mathrm{R}_k^{-1}$, so running time is significantly reduced. Using the above three techniques, all the markers on the genome are scanned and some potentially associated markers are selected. In the second stage, all the potentially associated markers are placed into one model, their effects are estimated by empirical Bayes, and all the non-zero effects are further identified by likelihood ratio test for true QTNs.

To further reduce the running time, the above model transformation is changed from $\mathbf{y}^* = \mathbf{U}^T \mathbf{y}$ in Wang et al. [1] into $\mathbf{y}^* = \mathbf{R}^{-\frac{1}{2}} \mathbf{U}^T \mathbf{y}$ in Tamba & Zhang [2] or **Supplementary material A**, where $\mathbf{R} = \mathbf{D}\hat{\lambda} + \mathbf{I}$. Using the results of Miller [3], thus, $|\mathbf{H}|$, $\mathbf{H}^{-1}$ and the quadratic term with the form of $\boldsymbol{\eta}^T \mathbf{H}^{-1} \boldsymbol{\tau}$ can be quickly and easily calculated, where $\mathbf{H}=\mathbf{Z}_{ci}\mathbf{Z}_{ci}^T\omega_i + \mathbf{I}$, and $\mathbf{Z}_{ci}$ is a vector of genotype indicators for the *k*th marker. In addition, least angle regression [4] is used to select $n-1$ variables to be included in the multi-locus genetic model if the number of potentially associated SNPs at the first stage is more than the sample size *n*. This method named FASTmrMLM.

##### *ISIS EM-BLASSO*

Tamba et al. [5] developed an iterative modified-sure independence screening (ISIS) implemented by expectation-maximization (EM)-Bayesian LASSO (BLASSO), which is referred to as ISIS EM-BLASSO. In the first stage, the ISIS method was used to reduce the number of SNPs to a moderate size. In other words, we first reduce the number of SNPs by selecting only those that are significantly correlated with the trait at the 0.01 level of significance. At this case, slight correlations between SNPs and the trait can be captured. Then, the selected SNP effects are shrunk by SCAD in order to select relevant SNPs. The procedure is replicated twice, and all the selected SNPs are potentially associated with the trait. The second stage is the same as that in mrMLM. Note that both EM-BLASSO and empirical Bayes are derived from Xu [6], although different names are used in our studies.

***Fast multi-locus random-SNP-effect EMMA*** (***FASTmrEMMA***)

Wen et al. [7] proposed FASTmrEMMA, which is a multi-locus GWAS method. The first two techniques in mrMLM of Wang et al. [1] are also adopted in FASTmrEMMA. However, two new techniques are implemented. First, we do pre-multiplication $\mathbf{C}=\mathbf{Q}_1\mathbf{\Lambda}_r^{-\frac{1}{2}}\mathbf{Q}_1^T$ for the standard mixed linear model equation, where $\mathbf{B}=\hat{\lambda}\mathbf{Z}\mathbf{K}\mathbf{Z}^T+\mathbf{I}_n=\left(\mathbf{Q}_1\mathbf{\Lambda}_r^{\frac{1}{2}}\mathbf{Q}_1^T\right)\left(\mathbf{Q}_1\mathbf{\Lambda}_r^{\frac{1}{2}}\mathbf{Q}_1^T\right)$, $r=\mathrm{rank}(\mathbf{B})$ and $\mathbf{Z}$ was the design matrix for polygenic effect. Its purpose is to whiten the covariance of kinship matrix K and residual noise. Then, the nonzero eigen decomposition of matrix $\mathbf{x}_C\mathbf{x}_C^T$ is the same as that of $\mathbf{x}_C^T\mathbf{x}_C$ (a positive number), where $\mathbf{x}_C=\mathbf{C}\mathbf{x}$ and $\mathbf{x}$ was an vector of marker genotypes. This means that the number of nonzero eigenvalues is specified as one. As a result, all the formulae in the estimation of all the parameters can be simply and explicitly expressed, as described in EMMA. Using the above four techniques, all the markers on the genome are scanned and some potentially associated markers are obtained in the first stage. The second stage is the same as that in the mrMLM.

***pKWmEB***

Although non-parametric methods in GWAS are robust in QTN detection, the absence of a polygenic background control in single-marker association in genome-wide scans results in a high false positive rate. To overcome this shortcoming, the algorithm of Wen et al. [7] is used to whiten the covariance matrix of kinship matrix K (polygenic background) and residual noise. Using the transferred model, the Kruskal-Wallis test along with least angle regression can be used to select all the potentially associated markers. All the selected markers are further evaluated by empirical Bayes and likelihood ratio test for true QTN detection. This is the pKWmEB method proposed by Ren et al. [8].

***pLARmEB***

The first stages in all the above methods are involved in the one-dimensional genome-wide scan by testing one marker at a time. If we want to test all the markers on one chromosome at a time while the other markers are viewed as polygenic background, the model transformation of Wen et al. [7] can be used to whiten the covariance matrix of kinship matrix K and residual noise. At the transferred model, least angle regression is used to select the *t* most potentially associated SNPs from all the markers on each chromosome (*t*=198 in the Monte Carlo simulation studies, and 50 in real data analysis). The second stage is the same as that in the mrMLM. This is the pLARmEB method of Zhang et al. [9].

All the above six methods are implemented in the R software package mrMLM. The critical P-value of significance is set as 0.0002, which is converted from a LOD score of 3.0 in the test statistics using $p = \Pr(\chi^2_{df=1} > 3.0 \times 4.61) = 0.0002$ [1]. mrMLM v4.0 and mrMLM.GUI v4.0 are freely available online at https://cran.r-project.org/web/packages/mrMLM/index.html and https://cran.r-project.org/web/packages/mrMLM.GUI/index.html, respectively. The relationship among the above six methods was showed in Fig 1a.

***Genome-wide efficient mixed model association (GEMMA)***

This is an existing single-locus genome scan method, a fixed model version of the original MLM [10], and is used as the gold standard of single-locus model method for comparison. It is implemented in the C software GEMMA (http://www.xzlab.org/software.html) [10]. The P-value threshold of significance is set as 0.05/*m*, where *m* is the number of markers.

***Efficient mixed-model association eXpedited* (*EMMAX*)**

EMMAX [11] is an extension of EMMA [12], and both methods are existing single-locus genome scan GWAS methods. If each QTN explains only a small fraction of phenotypic variation for complex traits, this allows us to avoid the repetitive variance component estimation procedure. Although there is almost no improvement for statistical power in QTN detection, the running time is significantly reduced. The EMMAX software is available at http://genetics.cs.ucla.edu/emmax/. The P-value threshold of significance is the same as that for GEMMA.

***FarmCPU***

This is an existing multi-locus GWAS method [13] and is used as the gold standard of multi-locus model method for comparison. FarmCPU iteratively uses fixed and random effect models for GWAS. The P-value threshold of significance is the same as that for GEMMA. The method is implemented in the R software package MVP v2.0 (https://github.com/XiaoleiLiuBio/MVP).

#### References


1 Wang SB, Feng JY, Ren WL, Huang B, Zhou L, Wen YJ, et al. Improving power and accuracy of genome-wide association studies via a multi-locus mixed linear model methodology. **Sci Rep** 2016;**6:**19444.

2 Tamba CL, Zhang YM. A fast mrMLM algorithm for multi-locus genome-wide association studies. **bioRxiv**, 2018. doi: 10.1101/341784.

3 Miller K. On the inverse of the sum of matrices. **Mathematics Magazine** 1981;**54**:67–72.

4 Efron B, Hastie T, Johnstone I, Robert T. Least angle regression. **Ann Stat** 2004;**32**:407–99.

5 Tamba CL, Ni YL, Zhang YM. Iterative sure independence screening EM Bayesian LASSO algorithm for multi-locus genome-wide association studies. **PLoS Comput Biol** 2017;**13:**e1005357.

6 Xu S. An expectation-maximization algorithm for the Lasso estimation of quantitative trait locus effects. **Heredity** 2010;**105**:483–94.

7 Wen YJ, Zhang H, Ni YL, Huang B, Zhang J, Feng JY, et al. Methodological implementation of mixed linear models in multi-locus genome-wide association studies. **Brief Bioinform** 2018;**19:**700–12.

8 Ren WL, Wen YJ, Dunwell JM, Zhang YM. pKWmEB: Integration of Kruskal-Wallis test with empirical Bayes under polygenic background control for multi-locus genome-wide association study. **Heredity** 2018; **120:**208–18.

9 Zhang J, Feng JY, Ni YL, Wen YJ, Niu Y, Tamba CL, et al. pLARmEB: integration of least angle regression with empirical Bayes for multilocus genome-wide association studies. **Heredity** 2017;**118:**517–24.

10 Zhou X, Stephens M. Genome-wide efficient mixed-model analysis for association studies. **Nat Genet** 2012; **44:**821–6

11 Kang HM, Sul JH, Service SK, Zaitlen NA, Kong SY, Freimer NB, et al. Variance component model to account for sample structure in genome-wide association studies. **Nat Genet** 2010;**42:**348–54.

12 Kang HM, Zaitlen NA, Wade CM, Kirby A, Heckerman D, Daly MJ, et al. Efficient control of population structure in model organism association mapping. **Genetics** 2008;**178:**1709–23.

13 Liu X, Huang M, Fan B, Bucklers ES, Zhang Z. Iterative usage of fixed and random effect models for powerful and efficient genome-wide association studies. **PLoS Genet** 2016;**12:**e1005767.

#### Supplementary material C. Real data analyses in rice, maize and Simmental beef cattle

To test the performances of the software package mrMLM v4.0, three real datasets in rice [1], maize [2] and Simmental beef cattle [3] were downloaded from the Rice SNP-Seek Database (http://snp-seek.irri.org./index.zul), the Maizego (http://www.maizego.org/) and https://doi.org/10.5061/dryad.4qc06, respectively, and reanalyzed in this study. In the above three datasets, the traits of interest were grain width, oil concentration and kidney weight, respectively, the number of phenotypic accessions was 2262, 362, and 1136, respectively, and the number of markers was 1.01, 1.06, and 0.67 million, respectively.

##### *Real data analyses in rice*

We re-analyzed the above rice dataset of Wang et al. [1] on the second server (Intel(R) Xeon(R) Gold 6130 CPU @ 2.10GHz, 64 processors and 629G memory). Population structure was calculated by package ADMIXTURE from rice 3K-RG core SNP (version 0.4, http://snp-seek.irri.org./_download.zul) with K = 9 (Wang et al. [1]), the first column was deleted in the GWAS, and Kinship matrix was calculated by package mrMLM v4.0. This dataset was re-analyzed by mrMLM, FASTmrMLM, FASTmrEMMA, pLARmEB, pKWmEB and ISIS EM-BLASSO methods in the package mrMLM v4.0 (Supplementary material B). The default parameters in the above methods were used in this study.

Their total running times for the above six methods were 9.56, 3.37, 11.58, 5.09, 6.13 and 1.06 (hours), respectively. Clearly, ISIS EM-BLASSO is the least followed by FASTmrMLM, pLARmEB, pKWmEB, and mrMLM; the FASTmrEMMA is the maximum. The total numbers of QTNs significantly associated with rice grain width using the above six methods (mrMLM, FASTmrMLM, FASTmrEMMA, pLARmEB, pKWmEB, and ISIS EM-BLASSO) were 73, 77, 42, 59, 17 and 31, respectively (Table

S2), indicating quite a number of QTNs (mean: 49.8) to be identified by the above multi-locus GWAS methods.

If the purpose of users is to mine candidate genes, the previously reported genes may be viewed as true genes. All the previously reported genes around the above QTNs are listed in Table S3. In addition, more important thing is to predict new candidate genes. In this case, we need to obtain all the genes around all the above QTNs and their annotations in *Oryza sativa* (https://rapdb.dna.affrc.go.jp/download/irgsp1.html) and *Arabidopsis thaliana* (https://genome.jgi.doe.gov/portal/pages/dynamicOrganismDownload.jsf?organism=Osativa). As a result, 136 genes related to seed developments had been found. All the potentially candidate genes were used to conduct KEGG analysis. Fifteen genes were associated with seed development (Table S4). We further used the datasets of gene expression levels in Nipponbare (http://rice.plantbiology.msu.edu/index.shtml), Minghui 63 and Zhenshan 97 (https://www.ncbi.nlm.nih.gov/geo/query/acc.cgi?acc=GSE19024) to mine candidate genes. As a result, *Os02g0115900* and *Os05g0182500* were predicted to be the most likely candidate genes (Table S4).

***Real data analyses in maize***

We re-analyzed the above maize dataset in Li et al. [2] on the third server (Intel(R) Xeon(R) CPU E5-2680 v2 @ 2.80GHz, 40 processors and 504G memory). Population structure was calculated by package ADMIXTURE with K = 3 [2], the software automatically deleted the column whose sum is the smallest, and Kinship matrix was calculated by package mrMLM v4.0. This dataset was re-analyzed by mrMLM, FASTmrMLM, FASTmrEMMA, pLARmEB, pKWmEB and ISIS EM-BLASSO methods in the package mrMLM v4.0 (Supplementary material B). The default parameters in the above methods were used in this study.

The total numbers of QTNs significantly associated with maize oil concentration using the above six methods were 42, 43, 31, 29, 17, and 6, respectively (Table S5), indicating quite a number of QTNs (mean: 28) to be identified by the above multi-locus GWAS methods.

In order to mine candidate genes that are significantly associated with oil concentration in maize, we obtained all the genes around all the above QTNs and got their annotations in *Zea mays* (http://www.gramene.org/ and http://ensembl.gramene.org/Zea_mays/Info/Index) and *Arabidopsis thaliana* (https://www.arabidopsis.org/). All the candidate genes around the above QTNs are listed in Table S6. As a result, 23 genes were found to be associated with oil concentration. Among these genes, 13 were detected only by the software mrMLM, and 10 were detected commonly by the mrMLM software and in Li et al. [2].

***Real data analyses in Simmental beef cattle***

We re-analyzed the above Simmental beef cattle dataset in Zhu et al. [3] on the second server (Intel(R) Xeon(R) Gold 6130 CPU @ 2.10GHz, 64 processors and 629 G memory). R function prcomp() was used to conduct principal component analysis. As shown in the below Figure, all the maize lines were clustered into five groups, in other words, there were five sub-populations in this association population. The five principal components were used to correct the effect of population structure on GWAS for kidney weight in Simmental beef cattle.

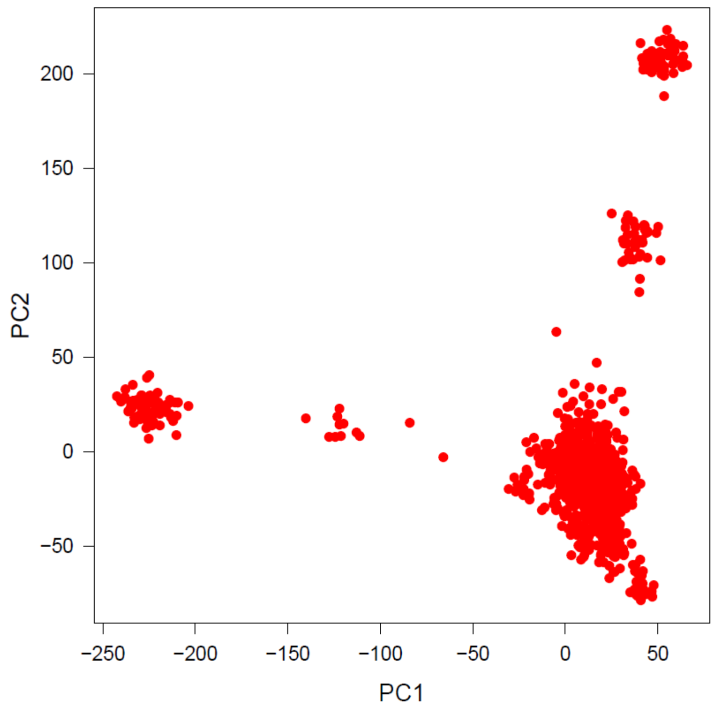


The software mrMLM v4.0 was used to calculate Kinship matrix. In this dataset, there were four covariates, including gender (categorical variable), birth year (categorical variable), body weight before experiment (continuous variable) and the number of days during fattening period (continuous variable). The last two continuous covariates were transferred into categorical variables by clustering them into five and six clusters, respectively. This dataset was re-analyzed by mrMLM, FASTmrMLM, FASTmrEMMA, pLARmEB, pKWmEB and ISIS EM-BLASSO in the software mrMLM v4.0 (Supplementary material B).

The total numbers of QTNs, detected by the above methods, for kidney weight in Simmental beef cattle were 4, 55, 167, 117, 8 and 48, respectively (Table S7), indicating quite a number of QTNs (mean: 66.5) to be identified by the above multi-locus GWAS methods.

In order to mine candidate genes that are significantly associated with kidney weight, we obtained all the genes around all the detected QTNs and got their annotations in *Bos taurus* (http://oct2018.archive.ensembl.org/Bos_taurus/Info/Index?db=core). All the candidate genes around the above QTNs are listed in Table S8. Among the previously reported genes, *MECOM* was identified commonly by mrMLM and in An et al. [4], and

*LCORL* and *NCAPG*, which are very important genes for kidney weight in cattle, were detected only by the mrMLM (Table S8).

#### Reference


1. Wang W, Mauleon R, Hu Z, Chebotarov D, Tai S, Wu Z, et al. Genomic variation in 3,010 diverse accessions of Asian cultivated rice. Nature 2018;557:43–49.
2. Li H, Peng Z, Yang X, Wang W, Fu J, Wang J, et al. Genome-wide association study dissects the genetic architecture of oil biosynthesis in maize kernels. Nat Genet 2013;45:43–50.
3. Zhu B, Zhu M, Jiang J, Niu H, Wang Y, Wu Y, et al. The impact of variable degrees of freedom and scale parameters in Bayesian methods for genomic prediction in Chinese Simmental beef cattle. PLoS ONE 2016; 11:e0154118.
4. An B, Xia J, Chang T, Wang X, Miao J, Xu L, et al. Genome-wide association study identifies loci and candidate genes for internal organ weights in Simmental beef cattle. Physiol Genomics 2018;**50**:523–31.

**Table S1**. The speedup in parallel computing under various numbers of CPUs and various GWAS approaches

| Number of CPUs | Multi-locus GWAS methods | | | | | |
|---|---|---|---|---|---|---|
| | mrMLM | FASTmrMLM | FASTmrEMMA | pLARmEB | pKWmEB | ISIS EM-BLASSO |
| 1 | 1.00 | 1.00 | 1.00 | 1.00 | 1.00 | 1.00 |
| 2 | 1.65 | 1.66 | 1.90 | 1.85 | 1.91 | 1.95 |
| 3 | 2.07 | 2.20 | 2.40 | 2.45 | 2.79 | 2.31 |
| 4 | 2.45 | 2.64 | 2.89 | 2.88 | 3.38 | 2.74 |
| 5 | 3.10 | 3.24 | 3.26 | 2.66 | 3.89 | 2.78 |
| 6 | 3.23 | 3.42 | 3.56 | 2.69 | 4.31 | 2.74 |
| 7 | 3.52 | 3.58 | 3.83 | 2.64 | 4.90 | 2.69 |

**Table S2**. All the QTNs for grain width in rice detected by mrMLM, FASTmrMLM, FASTmrEMMA, pLARmEB, pKWmEB and ISIS EBLASSO

| Chr | Position (bp) | QTN effect | LOD score | $-\log_{10}(P)$ | $r^2$ (%) | Method | Chr | Position (bp) | QTN effect | LOD score | $-\log_{10}(P)$ | $r^2$ (%) | Method |
|---|---|---|---|---|---|---|---|---|---|---|---|---|---|
| 1 | 6921155 | 0.0388 | 5.37 | 6.18 | 0.665 | mrMLM | 1 | 1831596 | -0.0354 | 3.19 | 3.90 | 0.1383 | FASTmrEMMA |
| 1 | 1831596 | -0.0331 | 4.79 | 5.57 | 0.5442 | mrMLM | 1 | 6491279 | -0.0798 | 5.38 | 6.20 | 0.836 | FASTmrEMMA |
| 1 | 23986872 | -0.0237 | 4.15 | 4.91 | 0.2814 | mrMLM | 1 | 10206576 | -0.0502 | 3.31 | 4.03 | 0.2181 | FASTmrEMMA |
| 1 | 41933338 | -0.0678 | 5.19 | 6.00 | 1.1884 | mrMLM | 1 | 39242548 | -0.0856 | 4.88 | 5.67 | 0.9135 | FASTmrEMMA |
| 1 | 19591075 | 0.0627 | 3.91 | 4.65 | 1.2254 | mrMLM | 1 | 41933338 | -6.38E-05 | 3.28 | 3.99 | 1.51E-07 | FASTmrEMMA |
| 2 | 33721702 | -0.0385 | 4.76 | 5.55 | 0.3686 | mrMLM | 2 | 9896989 | -0.093 | 6.57 | 7.42 | 0.6323 | FASTmrEMMA |
| 2 | 24640538 | 0.0269 | 3.45 | 4.17 | 0.2932 | mrMLM | 2 | 21908502 | -0.1073 | 5.95 | 6.78 | 1.4443 | FASTmrEMMA |
| 3 | 29849519 | 0.2325 | 13.02 | 14.02 | 1.5649 | mrMLM | 2 | 33773893 | -0.0859 | 3.15 | 3.86 | 1.0211 | FASTmrEMMA |
| 3 | 16733441 | -0.0475 | 12.00 | 12.98 | 1.2026 | mrMLM | 3 | 7230027 | -0.0679 | 4.21 | 4.97 | 0.5041 | FASTmrEMMA |
| 3 | 4919241 | 0.0512 | 3.70 | 4.43 | 0.9376 | mrMLM | 3 | 16733441 | -0.0603 | 5.01 | 5.81 | 0.4777 | FASTmrEMMA |
| 3 | 35769449 | 0.025 | 4.65 | 5.44 | 0.349 | mrMLM | 3 | 25332258 | 0.0628 | 4.70 | 5.48 | 0.478 | FASTmrEMMA |
| 3 | 7230027 | -0.0442 | 4.69 | 5.47 | 0.8871 | mrMLM | 3 | 35769449 | 0.0415 | 3.93 | 4.68 | 0.2303 | FASTmrEMMA |
| 3 | 26707321 | -0.0272 | 3.78 | 4.52 | 0.3492 | mrMLM | 4 | 1563119 | 0.0882 | 8.07 | 8.96 | 0.5889 | FASTmrEMMA |
| 4 | 837855 | 0.0538 | 4.99 | 5.78 | 0.7654 | mrMLM | 4 | 31705285 | -0.0662 | 4.20 | 4.96 | 0.414 | FASTmrEMMA |
| 4 | 32080864 | 0.0371 | 3.91 | 4.66 | 0.7407 | mrMLM | 4 | 32080864 | 0.0869 | 5.86 | 6.69 | 0.641 | FASTmrEMMA |
| 4 | 31705285 | -0.0335 | 4.03 | 4.78 | 0.4558 | mrMLM | 5 | 4833479 | -0.0724 | 4.25 | 5.02 | 0.5812 | FASTmrEMMA |
| 4 | 2404846 | 0.0392 | 3.56 | 4.29 | 0.4954 | mrMLM | 5 | 5361276 | -0.1108 | 3.29 | 4.00 | 1.595 | FASTmrEMMA |
| 4 | 10435637 | 0.0539 | 3.65 | 4.39 | 0.8709 | mrMLM | 5 | 5376243 | 0.2849 | 10.81 | 11.77 | 6.7119 | FASTmrEMMA |
| 4 | 22092126 | 0.0464 | 3.85 | 4.59 | 0.5928 | mrMLM | 5 | 5383914 | -0.1003 | 3.34 | 4.05 | 0.9649 | FASTmrEMMA |

| | | | | | | | | | | | | | |
|---|---|---|---|---|---|---|---|---|---|---|---|---|---|
| 5 | 5308999 | 0.0999 | 21.51 | 22.61 | 3.4461 | mrMLM | 5 | 27897307 | 0.0675 | 5.16 | 5.96 | 0.307 | FASTmrEMMA |
| 5 | 5007414 | -0.0662 | 12.60 | 13.59 | 2.011 | mrMLM | 6 | 1179404 | -0.0524 | 5.91 | 6.74 | 0.3495 | FASTmrEMMA |
| 5 | 5422688 | -0.0334 | 4.93 | 5.73 | 0.6056 | mrMLM | 6 | 27500790 | 0.0918 | 3.57 | 4.29 | 0.2549 | FASTmrEMMA |
| 5 | 27902614 | -0.1072 | 10.87 | 11.82 | 1.3452 | mrMLM | 7 | 16169282 | -0.0706 | 5.85 | 6.68 | 0.4404 | FASTmrEMMA |
| 5 | 5656974 | -0.0902 | 4.50 | 5.28 | 0.4653 | mrMLM | 7 | 19316334 | -0.0673 | 5.92 | 6.75 | 0.6084 | FASTmrEMMA |
| 5 | 5606561 | -0.0406 | 4.57 | 5.34 | 0.8252 | mrMLM | 7 | 21323336 | 0.0395 | 3.36 | 4.07 | 0.1957 | FASTmrEMMA |
| 5 | 762855 | -0.0453 | 4.03 | 4.78 | 0.7378 | mrMLM | 7 | 21666972 | 0.0822 | 4.78 | 5.57 | 0.9378 | FASTmrEMMA |
| 5 | 3627700 | 0.0376 | 3.95 | 4.70 | 0.6338 | mrMLM | 7 | 28321075 | 0.1372 | 8.42 | 9.32 | 0.9812 | FASTmrEMMA |
| 5 | 13136371 | -0.0563 | 7.23 | 8.10 | 1.4307 | mrMLM | 7 | 29606768 | 0.0523 | 3.34 | 4.06 | 0.2785 | FASTmrEMMA |
| 5 | 23667256 | -0.0454 | 3.66 | 4.39 | 0.5525 | mrMLM | 8 | 861782 | -0.058 | 4.86 | 5.65 | 0.3535 | FASTmrEMMA |
| 6 | 6346438 | 0.0883 | 4.78 | 5.57 | 0.3203 | mrMLM | 8 | 17843449 | -0.0618 | 4.42 | 5.19 | 0.5114 | FASTmrEMMA |
| 6 | 1179404 | -0.0273 | 4.56 | 5.34 | 0.4377 | mrMLM | 8 | 26217209 | 0.0521 | 4.85 | 5.64 | 0.2712 | FASTmrEMMA |
| 6 | 17265592 | -0.0384 | 5.06 | 5.86 | 0.8582 | mrMLM | 9 | 4595714 | -0.071 | 4.29 | 5.05 | 0.5662 | FASTmrEMMA |
| 6 | 30550157 | 0.0343 | 3.24 | 3.95 | 0.6941 | mrMLM | 9 | 6751029 | 0.0418 | 3.04 | 3.74 | 0.2434 | FASTmrEMMA |
| 7 | 29606768 | 0.0361 | 3.68 | 4.42 | 0.7402 | mrMLM | 9 | 8525240 | -0.048 | 4.07 | 4.83 | 0.2603 | FASTmrEMMA |
| 7 | 23624287 | 0.0368 | 5.21 | 6.01 | 0.4902 | mrMLM | 9 | 21359117 | -0.1137 | 9.10 | 10.02 | 1.2987 | FASTmrEMMA |
| 7 | 1579705 | -0.0523 | 4.70 | 5.48 | 0.7191 | mrMLM | 10 | 9280914 | -0.0837 | 4.70 | 5.48 | 0.447 | FASTmrEMMA |
| 7 | 21780181 | 0.0471 | 5.85 | 6.67 | 1.3002 | mrMLM | 10 | 19947905 | -0.1627 | 10.67 | 11.62 | 1.149 | FASTmrEMMA |
| 7 | 28456965 | 0.0412 | 3.85 | 4.60 | 0.8719 | mrMLM | 11 | 6915344 | 0.065 | 6.74 | 7.60 | 0.5872 | FASTmrEMMA |
| 8 | 26504638 | 0.1537 | 12.33 | 13.31 | 0.924 | mrMLM | 11 | 8857734 | 0.0529 | 3.57 | 4.30 | 0.1645 | FASTmrEMMA |
| 8 | 26249593 | 0.0654 | 3.03 | 3.73 | 0.2659 | mrMLM | 11 | 15133268 | -0.0885 | 6.48 | 7.33 | 0.4558 | FASTmrEMMA |
| 8 | 1660498 | 0.0526 | 3.05 | 3.75 | 1.6254 | mrMLM | 11 | 19526177 | -0.1052 | 6.69 | 7.54 | 0.4356 | FASTmrEMMA |

| | | | | | | | | | | | | | |
|---|---|---|---|---|---|---|---|---|---|---|---|---|---|
| 8 | 26019239 | -0.0414 | 3.58 | 4.31 | 0.5862 | mrMLM | 11 | 27020790 | 0.0506 | 4.59 | 5.37 | 0.2833 | FASTmrEMMA |
| 8 | 24755914 | -0.0297 | 3.50 | 4.22 | 0.4746 | mrMLM | 1 | 1831596 | -0.0183 | 3.26 | 3.97 | 0.1475 | pLARmEB |
| 8 | 5967659 | 0.0305 | 3.11 | 3.81 | 0.3142 | mrMLM | 1 | 26469542 | -0.0537 | 5.27 | 6.08 | 0.2973 | pLARmEB |
| 8 | 7641402 | -0.0295 | 3.07 | 3.77 | 0.496 | mrMLM | 1 | 31772747 | -0.0516 | 7.05 | 7.91 | 0.4934 | pLARmEB |
| 8 | 3881704 | -0.0577 | 6.20 | 7.04 | 1.3385 | mrMLM | 1 | 34597034 | 0.033 | 4.37 | 5.14 | 0.3379 | pLARmEB |
| 9 | 11745151 | -0.0688 | 3.50 | 4.23 | 0.2423 | mrMLM | 1 | 40027771 | 0.0589 | 4.10 | 4.85 | 0.258 | pLARmEB |
| 9 | 21393671 | -0.082 | 11.22 | 12.18 | 3.054 | mrMLM | 1 | 41933338 | -0.0451 | 3.90 | 4.64 | 0.4652 | pLARmEB |
| 9 | 15024826 | -0.0589 | 5.20 | 6.01 | 1.5083 | mrMLM | 2 | 11022186 | 0.0413 | 4.73 | 5.51 | 0.4998 | pLARmEB |
| 9 | 4905187 | 0.0424 | 4.31 | 5.08 | 0.8076 | mrMLM | 2 | 20311039 | -0.0175 | 4.03 | 4.78 | 0.1476 | pLARmEB |
| 9 | 10309830 | 0.0429 | 5.20 | 6.00 | 0.6999 | mrMLM | 2 | 33773893 | -0.0411 | 5.45 | 6.27 | 0.8569 | pLARmEB |
| 9 | 5861022 | 0.0551 | 4.91 | 5.71 | 0.7963 | mrMLM | 3 | 3676174 | -0.0705 | 5.25 | 6.05 | 0.5537 | pLARmEB |
| 9 | 21658026 | -0.0366 | 4.84 | 5.63 | 0.7688 | mrMLM | 3 | 5242456 | 0.0778 | 5.86 | 6.69 | 0.5017 | pLARmEB |
| 9 | 10501606 | -0.0474 | 4.35 | 5.12 | 0.9328 | mrMLM | 3 | 6865287 | 0.0251 | 5.98 | 6.81 | 0.2917 | pLARmEB |
| 9 | 6765802 | 0.0399 | 3.27 | 3.98 | 0.8269 | mrMLM | 3 | 16733441 | -0.0281 | 6.23 | 7.07 | 0.3728 | pLARmEB |
| 9 | 8525240 | -0.0294 | 3.52 | 4.25 | 0.4968 | mrMLM | 3 | 25456496 | 0.0311 | 3.17 | 3.87 | 0.1134 | pLARmEB |
| 9 | 7015224 | -0.0629 | 7.65 | 8.53 | 1.4863 | mrMLM | 3 | 35268655 | 0.0328 | 3.96 | 4.71 | 0.4187 | pLARmEB |
| 9 | 7648345 | 0.0305 | 3.02 | 3.71 | 0.5332 | mrMLM | 4 | 280312 | -0.065 | 5.06 | 5.86 | 0.8112 | pLARmEB |
| 10 | 19946757 | -0.0942 | 16.40 | 17.4 | 1.5924 | mrMLM | 4 | 5115500 | 0.0579 | 5.16 | 5.97 | 0.8066 | pLARmEB |
| 10 | 9282515 | -0.0368 | 3.90 | 4.64 | 0.514 | mrMLM | 4 | 13764697 | 0.0533 | 4.97 | 5.77 | 0.8885 | pLARmEB |
| 10 | 14652991 | 0.0433 | 3.51 | 4.24 | 1.0358 | mrMLM | 4 | 20536944 | -0.0231 | 3.08 | 3.78 | 0.244 | pLARmEB |
| 11 | 26988796 | 0.0284 | 5.10 | 5.90 | 0.397 | mrMLM | 4 | 31705285 | -0.0303 | 5.80 | 6.63 | 0.3291 | pLARmEB |
| 11 | 24873792 | -0.0287 | 4.65 | 5.43 | 0.4436 | mrMLM | 4 | 32238154 | 0.0425 | 4.56 | 5.34 | 0.4813 | pLARmEB |

| | | | | | | | | | | | | | |
|---|---|---|---|---|---|---|---|---|---|---|---|---|---|
| 11 | 19526177 | -0.0552 | 5.42 | 6.24 | 0.9821 | mrMLM | 5 | 972649 | -0.0431 | 3.32 | 4.04 | 0.8471 | pLARmEB |
| 11 | 693197 | -0.0452 | 5.57 | 6.39 | 0.9263 | mrMLM | 5 | 4859223 | -0.0402 | 8.18 | 9.08 | 0.7821 | pLARmEB |
| 11 | 8857734 | 0.0447 | 7.23 | 8.10 | 0.3815 | mrMLM | 5 | 5361276 | -0.0507 | 6.81 | 7.67 | 1.293 | pLARmEB |
| 11 | 10933886 | -0.0453 | 6.67 | 7.52 | 1.2151 | mrMLM | 5 | 5371949 | -0.0443 | 6.11 | 6.95 | 1.0271 | pLARmEB |
| 11 | 22303236 | -0.0223 | 3.72 | 4.45 | 0.2924 | mrMLM | 5 | 5376243 | 0.0771 | 11.60 | 12.57 | 1.8483 | pLARmEB |
| 11 | 22817547 | 0.0478 | 7.58 | 8.46 | 1.309 | mrMLM | 5 | 27902614 | -0.084 | 8.38 | 9.28 | 0.7336 | pLARmEB |
| 12 | 15690082 | -0.0502 | 4.19 | 4.95 | 0.9448 | mrMLM | 6 | 27500790 | 0.0593 | 3.36 | 4.08 | 1.2362 | pLARmEB |
| 12 | 9405542 | 0.0542 | 4.36 | 5.12 | 0.881 | mrMLM | 6 | 29387919 | -0.0352 | 3.20 | 3.90 | 0.3742 | pLARmEB |
| 12 | 19906987 | 0.0501 | 4.93 | 5.73 | 1.4848 | mrMLM | 7 | 9422683 | 0.0578 | 4.27 | 5.04 | 0.3554 | pLARmEB |
| 12 | 21802207 | -0.0547 | 6.83 | 7.69 | 1.0939 | mrMLM | 7 | 16169282 | -0.0324 | 5.53 | 6.35 | 0.4693 | pLARmEB |
| 1 | 2090441 | 0.0276 | 3.64 | 4.37 | 0.3867 | FASTmrMLM | 7 | 18257805 | 0.022 | 4.03 | 4.78 | 0.1868 | pLARmEB |
| 1 | 6921155 | 0.0323 | 5.67 | 6.49 | 0.4608 | FASTmrMLM | 7 | 21685804 | 0.0726 | 6.32 | 7.17 | 0.382 | pLARmEB |
| 1 | 10206576 | -0.0265 | 3.20 | 3.91 | 0.2369 | FASTmrMLM | 7 | 23624287 | 0.0323 | 4.73 | 5.51 | 0.3354 | pLARmEB |
| 1 | 29644733 | 0.0743 | 3.45 | 4.17 | 0.3047 | FASTmrMLM | 7 | 27677635 | -0.022 | 3.27 | 3.98 | 0.241 | pLARmEB |
| 1 | 39410519 | -0.0377 | 3.72 | 4.46 | 0.8305 | FASTmrMLM | 7 | 29606768 | 0.0289 | 5.35 | 6.16 | 0.4213 | pLARmEB |
| 1 | 40315934 | 0.0201 | 3.75 | 4.48 | 0.2392 | FASTmrMLM | 8 | 7641402 | -0.0222 | 3.28 | 3.99 | 0.2482 | pLARmEB |
| 1 | 41275767 | -0.0308 | 7.59 | 8.47 | 0.5615 | FASTmrMLM | 8 | 15609316 | 0.0381 | 6.52 | 7.37 | 0.5221 | pLARmEB |
| 1 | 41933338 | -0.058 | 5.34 | 6.15 | 0.868 | FASTmrMLM | 8 | 19624757 | 0.0539 | 3.26 | 3.97 | 0.2701 | pLARmEB |
| 2 | 762634 | 0.0153 | 3.48 | 4.21 | 0.1167 | FASTmrMLM | 8 | 26380813 | 0.1099 | 6.03 | 6.86 | 0.3428 | pLARmEB |
| 2 | 3174456 | -0.0347 | 3.37 | 4.08 | 0.519 | FASTmrMLM | 8 | 26504638 | 0.0986 | 4.25 | 5.01 | 0.3373 | pLARmEB |
| 2 | 10989478 | -0.0305 | 5.17 | 5.98 | 0.265 | FASTmrMLM | 8 | 27222965 | -0.0378 | 4.00 | 4.75 | 0.2868 | pLARmEB |
| 2 | 20946677 | -0.023 | 3.56 | 4.29 | 0.1157 | FASTmrMLM | 9 | 5823793 | 0.0453 | 4.18 | 4.94 | 0.2962 | pLARmEB |

| | | | | | | | | | | | | | |
|---|---|---|---|---|---|---|---|---|---|---|---|---|---|
| 2 | 21908502 | -0.0205 | 3.22 | 3.93 | 0.2358 | FASTmrMLM | 9 | 8525240 | -0.0172 | 3.03 | 3.72 | 0.151 | pLARmEB |
| 2 | 33725359 | -0.0404 | 4.17 | 4.93 | 0.464 | FASTmrMLM | 9 | 10309830 | 0.0349 | 3.69 | 4.42 | 0.4127 | pLARmEB |
| 2 | 35542100 | -0.0513 | 4.11 | 4.86 | 0.233 | FASTmrMLM | 9 | 21359117 | -0.0455 | 6.44 | 7.28 | 0.7894 | pLARmEB |
| 3 | 5242456 | 0.0864 | 7.04 | 7.90 | 0.6962 | FASTmrMLM | 9 | 21382154 | -0.0453 | 4.22 | 4.99 | 0.4309 | pLARmEB |
| 3 | 12488693 | 0.0351 | 3.46 | 4.19 | 0.2162 | FASTmrMLM | 9 | 21658026 | -0.0335 | 6.52 | 7.37 | 0.5733 | pLARmEB |
| 3 | 16733441 | -0.0322 | 7.06 | 7.93 | 0.5526 | FASTmrMLM | 10 | 9280914 | -0.0497 | 7.87 | 8.76 | 0.5155 | pLARmEB |
| 3 | 25320136 | 0.0246 | 5.54 | 6.36 | 0.3184 | FASTmrMLM | 10 | 19947905 | -0.0651 | 8.55 | 9.46 | 0.6941 | pLARmEB |
| 3 | 35141614 | -0.011 | 3.29 | 4.01 | 0.0571 | FASTmrMLM | 11 | 2557609 | -0.0254 | 4.38 | 5.15 | 0.3144 | pLARmEB |
| 3 | 35268655 | 0.0279 | 4.26 | 5.03 | 0.3424 | FASTmrMLM | 11 | 6920089 | 0.0302 | 4.35 | 5.11 | 0.3416 | pLARmEB |
| 4 | 315092 | 0.024 | 3.32 | 4.04 | 0.1306 | FASTmrMLM | 11 | 8050958 | -0.0486 | 4.43 | 5.20 | 0.7206 | pLARmEB |
| 4 | 4577211 | -0.0285 | 5.71 | 6.53 | 0.4701 | FASTmrMLM | 11 | 19526177 | -0.0451 | 5.37 | 6.18 | 0.5829 | pLARmEB |
| 4 | 19235815 | -0.032 | 4.88 | 5.67 | 0.6049 | FASTmrMLM | 11 | 19643504 | -0.028 | 4.11 | 4.87 | 0.2789 | pLARmEB |
| 4 | 31416786 | 0.019 | 3.46 | 4.18 | 0.1798 | FASTmrMLM | 11 | 21817990 | -0.0544 | 4.36 | 5.13 | 0.3093 | pLARmEB |
| 4 | 31705285 | -0.029 | 5.70 | 6.52 | 0.341 | FASTmrMLM | 11 | 24714951 | -0.0225 | 3.52 | 4.24 | 0.2382 | pLARmEB |
| 5 | 762855 | -0.0289 | 6.24 | 7.08 | 0.2999 | FASTmrMLM | 11 | 28796114 | -0.0197 | 3.05 | 3.75 | 0.1653 | pLARmEB |
| 5 | 913567 | -0.0437 | 4.83 | 5.62 | 0.914 | FASTmrMLM | 12 | 365399 | -0.044 | 4.68 | 5.46 | 0.3382 | pLARmEB |
| 5 | 4859223 | -0.0293 | 4.42 | 5.20 | 0.4699 | FASTmrMLM | 1 | 6491279 | -0.0316 | 3.41 | 4.13 | 3.8122 | pKWmEB |
| 5 | 5376243 | 0.1292 | 31.74 | 32.92 | 5.857 | FASTmrMLM | 2 | 7524480 | 0.0526 | 4.70 | 5.48 | 0.684 | pKWmEB |
| 5 | 14033709 | -0.0426 | 5.72 | 6.54 | 0.7804 | FASTmrMLM | 3 | 5242456 | 0.0797 | 3.60 | 4.33 | 2.4457 | pKWmEB |
| 5 | 23667256 | -0.0309 | 3.68 | 4.41 | 0.2553 | FASTmrMLM | 3 | 35769449 | 0.0299 | 4.11 | 4.87 | 1.1918 | pKWmEB |
| 6 | 1179404 | -0.0202 | 4.56 | 5.33 | 0.2405 | FASTmrMLM | 4 | 13764697 | 0.0482 | 3.84 | 4.59 | 2.2151 | pKWmEB |
| 6 | 27500790 | 0.0606 | 5.38 | 6.19 | 1.4533 | FASTmrMLM | 4 | 22310001 | -0.0435 | 3.28 | 3.99 | 2.2751 | pKWmEB |

| | | | | | | | | | | | | | |
|---|---|---|---|---|---|---|---|---|---|---|---|---|---|
| 6 | 29387919 | -0.0574 | 6.14 | 6.97 | 1.1207 | FASTmrMLM | 5 | 5371949 | -0.0662 | 11.63 | 12.59 | 5.2432 | pKWmEB |
| 7 | 19316334 | -0.0276 | 4.33 | 5.10 | 0.4483 | FASTmrMLM | 5 | 5361276 | -0.0477 | 3.74 | 4.48 | 5.0705 | pKWmEB |
| 7 | 21323336 | 0.0257 | 6.96 | 7.82 | 0.3887 | FASTmrMLM | 5 | 5343770 | 0.0827 | 3.45 | 4.17 | 6.6359 | pKWmEB |
| 7 | 23624287 | 0.0361 | 4.91 | 5.70 | 0.4704 | FASTmrMLM | 5 | 5291557 | -0.0747 | 5.09 | 5.89 | 0.2877 | pKWmEB |
| 7 | 27595785 | -0.0363 | 4.14 | 4.90 | 0.2785 | FASTmrMLM | 7 | 28334910 | 0.0454 | 6.25 | 7.09 | 1.503 | pKWmEB |
| 7 | 28321075 | 0.0475 | 5.03 | 5.83 | 0.535 | FASTmrMLM | 7 | 22895807 | 0.0484 | 3.42 | 4.14 | 2.7726 | pKWmEB |
| 7 | 29606768 | 0.0334 | 6.92 | 7.78 | 0.6319 | FASTmrMLM | 8 | 26504638 | 0.1378 | 4.42 | 5.19 | 1.3213 | pKWmEB |
| 8 | 1660498 | 0.04 | 3.63 | 4.36 | 0.9417 | FASTmrMLM | 9 | 21353073 | -0.0818 | 6.74 | 7.60 | 1.9548 | pKWmEB |
| 8 | 2410918 | -0.0218 | 4.59 | 5.37 | 0.2472 | FASTmrMLM | 10 | 19947905 | -0.0707 | 7.00 | 7.87 | 2.5975 | pKWmEB |
| 8 | 3881704 | -0.0258 | 3.25 | 3.96 | 0.2679 | FASTmrMLM | 11 | 27020790 | 0.0279 | 4.25 | 5.02 | 0.9692 | pKWmEB |
| 8 | 4880887 | 0.0179 | 3.49 | 4.21 | 0.1133 | FASTmrMLM | 12 | 21802207 | -0.0324 | 3.28 | 4.00 | 1.5278 | pKWmEB |
| 8 | 7568998 | 0.0453 | 6.31 | 7.15 | 0.5046 | FASTmrMLM | 1 | 3696298 | 0.0579 | 5.27 | 6.07 | 0.8724 | ISIS EM-BLASSO |
| 8 | 17392114 | 0.0417 | 6.16 | 7.00 | 0.7368 | FASTmrMLM | 1 | 6919215 | 0.0367 | 3.40 | 4.12 | 0.5815 | ISIS EM-BLASSO |
| 8 | 26249593 | 0.0471 | 3.42 | 4.14 | 0.1375 | FASTmrMLM | 3 | 3676174 | -0.0916 | 6.64 | 7.49 | 1.0518 | ISIS EM-BLASSO |
| 8 | 26292835 | 0.0573 | 3.80 | 4.54 | 0.8933 | FASTmrMLM | 3 | 5242456 | 0.0871 | 4.65 | 5.43 | 0.7074 | ISIS EM-BLASSO |
| 9 | 5861022 | 0.0537 | 5.72 | 6.54 | 0.7553 | FASTmrMLM | 3 | 25332258 | 0.0247 | 3.88 | 4.62 | 0.3156 | ISIS EM-BLASSO |
| 9 | 7015224 | -0.0398 | 5.39 | 6.20 | 0.5951 | FASTmrMLM | 4 | 8669122 | 0.0876 | 4.71 | 5.50 | 1.201 | ISIS EM-BLASSO |
| 9 | 7648345 | 0.0321 | 5.16 | 5.97 | 0.5894 | FASTmrMLM | 4 | 22310001 | -0.0411 | 3.64 | 4.37 | 0.7156 | ISIS EM-BLASSO |
| 9 | 8525240 | -0.02 | 4.54 | 5.32 | 0.2305 | FASTmrMLM | 4 | 31880075 | -0.0522 | 3.08 | 3.78 | 0.3618 | ISIS EM-BLASSO |
| 9 | 8900138 | -0.028 | 5.93 | 6.77 | 0.4167 | FASTmrMLM | 4 | 32080864 | 0.0341 | 3.19 | 3.90 | 0.6282 | ISIS EM-BLASSO |
| 9 | 10213296 | 0.023 | 4.57 | 5.35 | 0.2864 | FASTmrMLM | 5 | 4831052 | -0.0367 | 5.49 | 6.30 | 0.7348 | ISIS EM-BLASSO |
| 9 | 15024826 | -0.0412 | 4.93 | 5.72 | 0.7395 | FASTmrMLM | 5 | 5186176 | 0.0572 | 3.07 | 3.77 | 1.1965 | ISIS EM-BLASSO |

| | | | | | | | | | | | | | |
|---|---|---|---|---|---|---|---|---|---|---|---|---|---|
| 9 | 20251821 | 0.0412 | 5.91 | 6.74 | 0.3481 | FASTmrMLM | 5 | 5300234 | -0.0694 | 4.64 | 5.42 | 0.4589 | ISIS EM-BLASSO |
| 9 | 20559656 | -0.028 | 4.10 | 4.85 | 0.2477 | FASTmrMLM | 5 | 5361276 | -0.0678 | 11.63 | 12.60 | 2.6106 | ISIS EM-BLASSO |
| 9 | 21338733 | -0.0411 | 4.05 | 4.81 | 0.7589 | FASTmrMLM | 5 | 5371949 | -0.0607 | 9.68 | 10.62 | 2.1738 | ISIS EM-BLASSO |
| 9 | 22359906 | -0.029 | 3.65 | 4.38 | 0.4688 | FASTmrMLM | 5 | 5376243 | 0.07 | 7.01 | 7.88 | 1.7167 | ISIS EM-BLASSO |
| 10 | 3943372 | -0.0195 | 3.47 | 4.19 | 0.2111 | FASTmrMLM | 6 | 21315943 | 0.0297 | 3.09 | 3.80 | 0.3887 | ISIS EM-BLASSO |
| 10 | 4649756 | -0.0451 | 4.61 | 5.39 | 0.5017 | FASTmrMLM | 6 | 27500790 | 0.0499 | 3.74 | 4.48 | 0.9871 | ISIS EM-BLASSO |
| 10 | 19451104 | -0.0482 | 5.42 | 6.24 | 1.0079 | FASTmrMLM | 6 | 29387919 | -0.0442 | 3.13 | 3.83 | 0.6646 | ISIS EM-BLASSO |
| 10 | 19947905 | -0.0769 | 10.37 | 11.32 | 1.0915 | FASTmrMLM | 7 | 16169282 | -0.0308 | 4.29 | 5.05 | 0.4787 | ISIS EM-BLASSO |
| 11 | 693197 | -0.0271 | 3.65 | 4.38 | 0.3327 | FASTmrMLM | 7 | 21285184 | -0.051 | 3.96 | 4.71 | 1.0014 | ISIS EM-BLASSO |
| 11 | 7058901 | 0.0338 | 6.82 | 7.68 | 0.5245 | FASTmrMLM | 7 | 21666972 | 0.0375 | 5.20 | 6.01 | 0.8275 | ISIS EM-BLASSO |
| 11 | 8050958 | -0.0441 | 4.63 | 5.41 | 0.6697 | FASTmrMLM | 7 | 23951267 | -0.0477 | 3.74 | 4.48 | 0.6759 | ISIS EM-BLASSO |
| 11 | 8857734 | 0.0244 | 3.87 | 4.62 | 0.1139 | FASTmrMLM | 7 | 28321075 | 0.0493 | 5.81 | 6.63 | 0.5757 | ISIS EM-BLASSO |
| 11 | 10933886 | -0.0376 | 9.62 | 10.55 | 0.8381 | FASTmrMLM | 8 | 2410918 | -0.0218 | 3.06 | 3.76 | 0.2477 | ISIS EM-BLASSO |
| 11 | 19456239 | -0.0433 | 7.38 | 8.25 | 0.7129 | FASTmrMLM | 8 | 26504638 | 0.113 | 4.63 | 5.41 | 0.4993 | ISIS EM-BLASSO |
| 11 | 19643504 | -0.034 | 7.22 | 8.09 | 0.4641 | FASTmrMLM | 9 | 21393671 | -0.0579 | 4.87 | 5.66 | 1.5211 | ISIS EM-BLASSO |
| 11 | 21815738 | 0.0322 | 3.10 | 3.81 | 0.3844 | FASTmrMLM | 10 | 19946757 | -0.0609 | 6.19 | 7.03 | 0.6668 | ISIS EM-BLASSO |
| 11 | 23360520 | 0.036 | 3.30 | 4.01 | 0.4149 | FASTmrMLM | 11 | 6920089 | 0.0373 | 5.86 | 6.69 | 0.5874 | ISIS EM-BLASSO |
| 12 | 9405542 | 0.0397 | 3.78 | 4.52 | 0.4726 | FASTmrMLM | 11 | 11034491 | -0.0282 | 3.35 | 4.07 | 0.2883 | ISIS EM-BLASSO |
| 12 | 18270626 | -0.0318 | 3.59 | 4.32 | 0.4891 | FASTmrMLM | 12 | 365399 | -0.0431 | 3.73 | 4.47 | 0.365 | ISIS EM-BLASSO |
| 12 | 19906987 | 0.0279 | 3.55 | 4.28 | 0.4592 | FASTmrMLM | 12 | 21802207 | -0.0364 | 3.69 | 4.42 | 0.4837 | ISIS EM-BLASSO |
| 12 | 21802207 | -0.0434 | 7.89 | 8.78 | 0.6892 | FASTmrMLM | | | | | | | |

**Table S3.** Previously reported genes for grain width in rice around the QTNs identified by our multi-locus GWAS methods

| Chr | Marker position (bp) | Multi-locus GWAS | | | | Comparative genomics analysis | | | |
|---|---|---|---|---|---|---|---|---|---|
| | | QTN effect | LOD score | $r^2$ (%) | Method | MSU_locus | Gene | Functional annotation | Reference |
| Candidate genes associated with grain width | | | | | | | | | |
| 1 | 23986872 | -0.0237 | 4.15 | 0.28 | 1 | Os01g0625900 | *OsOFP2* | seed size, grain width | Schmitz et al. [6] |
| 2 | 7524480 | 0.0526 | 4.70 | 0.68 | 5 | Os02g0234200 | *FUWA* | grain shape, grain width | Chen et al. [7] |
| 3 | 3676174 | -0.0916~-0.0705 | 5.25~6.64 | 0.55~1.05 | 4, 6 | Os03g0175800 | *BG1* | grain size, grain width | Liu et al. [8] |
| 4 | 31880075 | -0.0522 | 3.08 | 0.36 | 6 | Os04g0645100 | *flo2* | grain size, grain width | She et al. [9] |
| 5 | 762855 | -0.0453~-0.0289 | 4.03~6.24 | 0.30~0.74 | 1, 2 | Os05g0115800 | *OsMKP1* | grain size, grain width | Guo et al. [10] |
| 5 | 3627700 | 0.0376 | 3.95 | 0.63 | 1 | Os05g0158500 | *GS5* | grain size, grain width | Xu et al. [11] |
| **5** | **5361276** | **-0.1108~-0.0477** | **3.29~11.63** | **1.29~5.07** | **3, 4, 5, 6** | **Os05g0187500** | ***GW5*** | **grain size, grain width** | **Liu et al. [12]** |
| 6 | 1179404 | -0.0524~-0.0202 | 4.56~5.90 | 0.24~0.44 | 1, 2, 3 | Os06g0130400 | *OsACS6* | grain size, grain width | Matsushima et al. [13] |
| 7 | 22895807 | 0.0484 | 3.42 | 2.77 | 5 | Os07g0580500 | *OsBZR1* | seed size, grain width | Zhu et al. [14] |
| 7 | 23951267 | -0.0477 | 3.74 | 0.68 | 6 | Os07g0603300 | *qGL7* | grain shape, grain width | Bai et al. [15] |
| 8 | 26019239 | -0.0414 | 3.58 | 0.59 | 1 | Os08g0537800 | *WTG1* | grain size, grain width | Huang et al. [16] |
| 8 | 27222965 | -0.0378 | 4.00 | 0.29 | 4 | Os08g0562500 | *SLG* | grain width | Feng et al. [17] |
| **9** | **21338733** | **-0.0411** | **4.05** | **0.76** | **2** | **Os09g0540800** | ***OsFD1*** | **transcription factors** | **Wang et al [5]** |
| Candidate genes associated with grain size | | | | | | | | | |
| 2 | 20311039 | -0.0175 | 4.03 | 0.15 | 4 | Os02g0554000 | *SDG725* | grain size | Sui et al. [18] |
| 2 | 24640538 | 0.0269 | 3.45 | 0.29 | 1 | Os02g0614100 | *OsNST1* | grain size | Zhang et al. [19] |
| 2 | 33721702 | -0.0385 | 4.76 | 0.37 | 1 | Os02g0787300 | *OsMKK4* | grain size | Duan et al. [20] |
| 3 | 5242456 | 0.0778~0.0871 | 3.60~7.04 | 0.50~2.45 | 2, 4, 5, 6 | Os03g0215400 | *OsMADS1* | grain shape, grain size | Liu et al. [21] |
| 3 | 6865287 | 0.0251 | 5.98 | 0.29 | 4 | Os03g0236900 | *OsAPC6* | grain size | Kumar et al. [22] |
| 3 | 7230027 | -0.0679~-0.0442 | 4.21~4.69 | 0.50~0.89 | 1, 3 | Os03g0254400 | *OspPLAIIIα* | grain size, seed size | Liu et al. [23] |
| 3 | 16733441 | -0.0603~-0.0281 | 5.014~12.00 | 0.37~1.20 | 1, 2, 3, 4 | Os03g0407400 | *GS3* | grain size, grain width | Fan et al. [24] |
| 3 | 35141614 | -0.0110 | 3.30 | 0.06 | 2 | Os03g0837300 | *OsNaPRT1* | grain size | Yan et al. [25] |
| 8 | 1660498 | 0.0400~0.0526 | 3.05~3.63 | 0.94~1.63 | 1, 2 | Os08g0137100 | *OsFIE2* | grain size | Na et al. [26] |
| 11 | 19643504 | -0.0340~-0.0280 | 4.11~7.22 | 0.28~0.46 | 2, 4 | Os11g0540500 | *IKU2* | grain width (Arabidopsis) | Luo et al. [27] |

Note: 1, 2, 3, 4, 5, and 6 represent mrMLM, FASTmrMLM, FASTmrEMMA, pLARmEB, pKWmEB, and ISIS EM-BLASSO, respectively. The genes with bold type were also detected by Wang et al. [5] (Nature 2018;557:43–49). Twenty-two genes were found at http://www.ricedata.cn/, and one gene was from Luo et al. [25] (Proc Nati Acad Sci U S A 2005; 102:17531–36) in *Arabidopsis*.

### References


5. Wang W, Mauleon R, Hu Z, Chebotarov D, Tai S, Wu Z, et al. Genomic variation in 3,010 diverse accessions of Asian cultivated rice. Nature 2018;557:43–49.
6. Schmitz AJ, Begcy K, Sarath G, Walia H. Rice ovate family protein 2 (*OFP2*) alters hormonal homeostasis and vasculature development. Plant Sci 2015;241:177–88.
7. Chen J, Gao H, Zheng XM, Jin M, Weng JF, Ma J et al. An evolutionarily conserved gene, *FUWA*, plays a role in determining panicle architecture, grain shape and grain weight in rice. Plant J 2015;83:427–38.
8. Liu L, Tong H, Xiao Y, Che R, Xu F, Hu B et al. Activation of *Big Grain1* significantly improves grain size by regulating auxin transport in rice. Proc Natl Acad Sci U S A 2015;112:11102–07.
9. She KC, Kusano H, Koizumi K, Yamakawa H, Hakata M, Imamura T, et al. A novel factor *FLOURY ENDOSPERM2* is involved in regulation of rice grain size and starch quality. Plant Cell 2010;22:3280–94.
10. Guo T, Chen K, Dong NQ, et al. *GRAIN SIZE* and *NUMBER1* negatively regulates the OsMKKK10- OsMKK4-OsMPK6 Cascade to coordinate the trade-off between grain number per panicle and grain size in rice. Plant Cell 2018;30:871–88.
11. Xu C, Yu L, Li Y, Xu X, Xu C, Li X, et al. Differential expression of *GS5* regulates grain size in rice. J Exp Bot 2015;66:2611-23.
12. Liu J, Chen J, Zheng X, Wu F, Lin Q, Heng Y et al. *GW5* acts in the brassinosteroid signalling pathway to regulate grain width and weight in rice. Nat Plants 2017;3:17043.
13. Matsushima R, Maekawa M, Kusano M, Tomita K, Kondo H, Nishimura H, et al. Amyloplast membrane protein *SUBSTANDARD STARCH GRAIN6* controls starch grain size in rice endosperm. Plant Physiol 2016;170:1445–59.
14. Zhu X, Liang W, Cui X, Chen M, Yin C, Luo Z, et al. Brassinosteroids promote development of rice pollen grains and seeds by triggering expression of Carbon Starved Anther, a MYB domain protein. Plant J 2015;82:570–81.
15. Bai X, Luo L, Yan W, Kovi MR, Zhan W, Xing Y. Genetic dissection of rice grain shape using a recombinant inbred line population derived from two contrasting parents and fine mapping a pleiotropic quantitative trait locus *qGL7*. BMC Genet 2010;11:16–27.

16. Huang K, Wang D, Duan P, Zhang B, Xu R, Li N, et al. *WIDE AND THICK GRAIN 1*, which encodes an otubain-like protease with deubiquitination activity, influences grain size and shape in rice. Plant J 2017;91: 849–60.
17. Feng Z, Wu C, Wang C, Roh J, Zhang L, Chen J, et al. *SLG* controls grain size and leaf angle by modulating brassinosteroid homeostasis in rice. J Exp Bot 2016;67:4241–53.
18. Sui P, Jin J, Ye S, Mu C, Gao J, Feng H, et al. H3K36 methylation is critical for brassinosteroid-regulated plant growth and development in rice. Plant J 2012;70:340–7.
19. Zhang B, Liu X, Qian Q, Liu L, Dong G, Xiong G, et al. Golgi nucleotide sugar transporter modulates cell wall biosynthesis and plant growth in rice. Proc Natl Acad Sci U S A 2011;108:5110–15.
20. Duan P, Rao Y, Zeng D, Yang Y, Xu R, Zhang B et al. *SMALL GRAIN 1*, which encodes a mitogen-activated protein kinase kinase 4, influences grain size in rice. Plant J 2014;77:547–57.
21. Liu Q, Han R, Wu K, Zhang J, Ye Y, Wang S, et al. G-protein betagamma subunits determine grain size through interaction with MADS-domain transcription factors in rice. Nat Commun 2018;9:852–64.
22. Kumar M, Basha PO, Puri A, Rajpurohit D, Randhawa GS, Sharma TR, et al. A candidate gene *OsAPC6* of anaphase-promoting complex of rice identified through T-DNA insertion. Funct Integr Genomics 2010;10: 349–58.
23. Liu G, Zhang K, Ai J, Deng X, Hong Y, Wang X, et al. Patatin-related phospholipase A, pPLAIIIα, modulates the longitudinal growth of vegetative tissues and seeds in rice. J Exp Bot 2015;66:6945–55.
24. Fan C, Xing Y, Mao H, Lu T, Han B, Xu C, et al. *GS3*, a major QTL for grain length and weight and minor QTL for grain width and thickness in rice, encodes a putative transmembrane protein. Theor Appl Genet 2006; 112:1164–71.
25. Yan S, Zou G, Li S, Wang H, Liu H, Zhai G, et al. Seed size is determined by the combinations of the genes controlling different seed characteristics in rice. Theor Appl Genet 2011; 123:1173-81.
26. Na JK, Seo MH, Yoon IS, Lee TH, Lee KO, Kim DY, et al. Involvement of rice Polycomb protein OsFIE2 in plant growth and seed size. Plant Biotechnology Reports 2012; 6:339–46.
27. Luo M, Dennis ES, Berger F, Peacock WJ, Chaudhury A. *MINISEED3* (*MINI3*), a WRKY family gene, and *HAIKU2* (*IKU2*), a Leucine-Rich repeat (LRR) KINASE gene, are regulators of seed size in *Arabidopsis*. Proc Nati Acad Sci U S A 2005; 102:17531–36.

**Table S4. Fifteen new candidate genes for rice grain size and development detected by six multi-locus GWAS methods**

| Chr | Position (bp) | Multi-locus genome-wide association studies | | | | KEGG analysis | | | | | Gene expressional analysis | |
|---|---|---|---|---|---|---|---|---|---|---|---|---|
| | | QTN effect | LOD | $r^2$ (%) | Method | ID | P-Value | Functional Annotation | Reference | Candidate gene | Nipponbare | Average expressional level |
| Two highly expressed genes, which are predicted to be associated with rice grain width | | | | | | | | | | | | |
| **2** | **762634** | **0.02** | **3.48** | **0.12** | **2** | **hsa04141** | **0.0153** | **Protein processing in endoplasmic reticulum** | **Takahashi et al.[28]** | **Os02g0115900** | **696.23, 187.01, 421.91, 748.59** | **12.26, 11.93, 10.60** |
| **5** | **4859223** | **-0.05~-0.03** | **4.42~8.18** | **0.47~0.78** | **2, 4** | **P00060** | **0.019** | **Ubiquitin proteasome pathway** | **Song et al.[29]** | **Os05g0182500** | **105.84, 61.56, 28.82, 45.0** | **10.51, 8.53, 8.63** |
| Thirteen new candidate genes, which are predicted to be associated with rice grain width | | | | | | | | | | | | |
| 1 | 41275767 | -0.03 | 7.59 | 0.56 | 2 | hsa00100 | 0.0005 | Steroid biosynthesis | Vriet et al.[30] | Os01g0940000 | 0, 0.30, 0, 0 | 3.99, 4.78, 3.26 |
| 4 | 4577211 | -0.03 | 5.71 | 0.47 | 2 | hsa00980 | 4.69E-06 | Metabolism of xenobiotics by cytochrome P450 | Chakrabarti et al.[31] | Os04g0167800 | 0, 0, 0, 0 | 3.80, 2.78, 3.01 |
| 4 | 5115500 | 0.06 | 5.16 | 0.81 | 4 | hsa00980 | 4.69E-06 | Metabolism of xenobiotics by cytochrome P450 | Chakrabarti et al.[31] | Os04g0174100 | 0, 0, 0, 0 | |
| 4 | 22310001 | -0.04~-0.04 | 3.28~3.64 | 0.72~2.28 | 5, 6 | hsa00600 | 0.0217 | Sphingolipid metabolism | Ishikawa et al.[32] | Os04g0447400 | 0.49, 0.58, 0, 0.15 | 3.90, 2.70, 3.09 |
| 5 | 4833479 | -0.07 | 4.25 | 0.58 | 3 | hsa00562 | 0.0479 | Inositol phosphate metabolism | Suzuki et al.[33] | Os05g0180600 | 6.05, 9.70, 5.28, 4.97 | 5.96, 4.23, 4.27 |
| 5 | 5291557 | -0.07 | 5.09 | 0.29 | 5 | hsa00520 | 0.0231 | Amino sugar and nucleotide sugar metabolism. | Zhang et al.[34] | Os05g0187100 | 11.42, 3.29, 0.27, 0.66 | 8.65, 6.72, 6.89 |
| 5 | 5656974 | -0.09 | 4.5 | 0.47 | 1 | hsa04120 | 0.0479 | Ubiquitin mediated proteolysis | Song et al.[29] | Os05g0193900 | 0.79, 1.16, 0, 0 | |
| 8 | 7568998 | 0.05 | 6.31 | 0.50 | 2 | P00060 | 0.019 | Ubiquitin proteasome pathway | Song et al. [29] | Os08g0224700 | 41.33, 29.70, 16.74, 22.93 | 9.39, 7.93, 7.98 |
| 9 | 4595714 | -0.07 | 4.29 | 0.57 | 3 | hsa00100 | 0.0005 | Steroid biosynthesis | Vriet et al.[30] | Os09g0262000 | 0, 0, 0, 0 | 3.24, 2.79, 2.77 |
| 9 | 5823793 | 0.05 | 4.18 | 0.30 | 4 | hsa00562 | 0.0479 | Inositol phosphate metabolism | Suzuki et al.[33] | Os09g0278300 | 0.86, 2.23, 0.52, 1.27 | |
| 9 | 7015224 | -0.06~-0.04 | 5.39~7.65 | 0.60~1.49 | 1, 2 | hsa04120 | 0.0479 | Ubiquitin mediated proteolysis | Song et al.[29] | Os09g0294300 | 2.81, 0.34, 0.94, 0.76 | 6.39, 3.62, 2.98 |
| 9 | 20251821 | 0.04 | 5.91 | 0.35 | 2 | hsa00562 | 0.0479 | Inositol phosphate metabolism | Suzuki et al.[33] | Os09g0518700 | 20.25, 51.81, 4.73, 9.16 | 9.37, 8.76, 9.28 |
| 10 | 9280914 | -0.08~-0.05 | 4.7 | 0.45~0.52 | 3, 4 | hsa00600 | 0.02167 | Sphingolipid metabolism | Ishikawa et al.[32] | Os10g0330600 | 0, 0, 0, 0 | 3.29, 2.71, 2.68 |

The six methods mrMLM, FASTmrMLM, FASTmrEMMA, pLARmEB, pKWmEB, and ISIS EM-BLASSO were marked by numbers 1 to 6, respectively. The numbers in the Nipponbare column were gene expression levels at four tissues and stages (seed-5 days after pollination (DAP); embryo-25 DAP; endosperm-25 DAP; seed-10 DAP), in which their average expression levels were 28.40, 26.73, 43.20 and 33.30 (FPKM), respectively (http://rice.plantbiology.msu.edu/index.shtml). The numbers in the column of average expressional level were average expressional levels of each gene at three stages (7, 14 and 21 DAP) between Minghui 63 and Zhenshan 97, and the average expression levels for all the genes were 5.63 (7 DAP), 4.76 (14 DAP) and 4.92 (21 DAP) (https://www.ncbi.nlm.nih.gov/geo/query/acc.cgi?acc=GSE19024). The two candidate genes with bold type are the most likely candidate genes.

#### References


28 Takahashi H, Saito Y, Kitagawa T, Morita S, Masumura T, Tanaka K, et al. A novel vesicle derived directly from endoplasmic reticulum is involved in the transport of vacuolar storage proteins in rice endosperm. Plant Cell Physiol 2005; 46:245–9.

29 Song XJ, Huang W, Shi M, Zhu MZ, Lin HX. A QTL for rice grain width and weight encodes a previously unknown RING-type E3 ubiquitin ligase. Nat Genet 2007; 39:623-30.

30 Vriet C, Russinova E, Reuzeau C. Boosting crop yields with plant steroids. Plant Cell 2012; 24:842–57.

31 Chakrabarti M, Zhang N, Sauvage C, Muños S, Blanca J, Cañizares J, et al. A cytochrome P450 regulates a domestication trait in cultivated tomato. Proc Natl Acad Sci U S A 2013;110:17125–30.

32 Ishikawa T, Ito Y, Kawaiyamada M. Molecular characterization and targeted quantitative profiling of the sphingolipidome in rice. Plant J 2016; 88:681–93.

33 Suzuki M, Tanaka K, Kuwano M, Yoshida KT. Expression pattern of inositol phosphate-related enzymes in rice (*Oryza sativa* L.): Implications for the phytic acid biosynthetic pathway. Gene 2007; 405:55–64.

34 Zhang B, Liu X, Qian Q, Liu L, Dong G, Xiong G, et al. Golgi nucleotide sugar transporter modulates cell wall biosynthesis and plant growth in rice. Proc Natl Acad Sci U S A 2011;108:5110–15.

**Table S5**. All the QTNs for oil concentration in maize detected by mrMLM, FASTmrMLM, FASTmrEMMA, pLARmEB, pKWmEB and ISIS EBLASSO

| Chr | Position (bp) | QTN effect | LOD score | $-\log_{10}(P)$ | $r^2$ (%) | Method | Chr | Position (bp) | QTN effect | LOD score | $-\log_{10}(P)$ | $r^2$ (%) | Method |
|---|---|---|---|---|---|---|---|---|---|---|---|---|---|
| 1 | 52762430 | -0.326 | 6.08 | 6.9185 | 0.593 | mrMLM | 10 | 113390827 | 0.055 | 4.14 | 4.9022 | 0.0987 | FASTmrMLM |
| 1 | 198416421 | 0.1638 | 7.73 | 8.6171 | 1.026 | mrMLM | 1 | 61716135 | 0.1268 | 3.21 | 3.9156 | 0.1484 | FASTmrEMMA |
| 1 | 296768029 | -0.197 | 4.10 | 4.8600 | 0.4604 | mrMLM | 2 | 24388940 | -0.2221 | 4.54 | 5.3196 | 0.4615 | FASTmrEMMA |
| 1 | 13536744 | -0.1207 | 4.56 | 5.3426 | 0.4347 | mrMLM | 2 | 53680157 | 0.4481 | 7.49 | 8.3641 | 0.8778 | FASTmrEMMA |
| 1 | 35740220 | 0.093 | 3.99 | 4.7383 | 0.3329 | mrMLM | 2 | 150020854 | 0.2362 | 3.16 | 3.8606 | 0.347 | FASTmrEMMA |
| 1 | 245479447 | -0.0731 | 3.90 | 4.6413 | 0.181 | mrMLM | 2 | 189219763 | 0.1956 | 3.78 | 4.5169 | 0.3031 | FASTmrEMMA |
| 1 | 292354425 | 0.1129 | 5.38 | 6.1910 | 0.3631 | mrMLM | 3 | 217876667 | 0.3502 | 8.07 | 8.9675 | 0.796 | FASTmrEMMA |
| 1 | 104610891 | -0.1088 | 6.41 | 7.2514 | 0.4119 | mrMLM | 4 | 5012443 | -0.2349 | 5.91 | 6.7352 | 0.5382 | FASTmrEMMA |
| 1 | 14847409 | -0.1221 | 5.18 | 5.9781 | 0.5568 | mrMLM | 4 | 6601755 | -0.4893 | 6.04 | 6.8765 | 0.8407 | FASTmrEMMA |
| 2 | 53680401 | -0.1785 | 5.96 | 6.7865 | 0.5254 | mrMLM | 4 | 47531927 | 0.2101 | 5.18 | 5.9800 | 0.393 | FASTmrEMMA |
| 2 | 24390872 | -0.1288 | 7.28 | 8.1531 | 0.6334 | mrMLM | 4 | 231411384 | -0.4372 | 6.24 | 7.0816 | 0.9882 | FASTmrEMMA |
| 2 | 183922016 | -0.0778 | 3.17 | 3.8711 | 0.2057 | mrMLM | 5 | 4769561 | -0.2764 | 8.17 | 9.0620 | 0.6749 | FASTmrEMMA |
| 2 | 150704340 | 0.073 | 3.61 | 4.3425 | 0.207 | mrMLM | 5 | 15186727 | -0.6589 | 11.60 | 12.5647 | 1.6394 | FASTmrEMMA |
| 2 | 193735059 | 0.1495 | 4.31 | 5.0750 | 0.599 | mrMLM | 5 | 25619033 | 0.2995 | 3.22 | 3.9292 | 0.3995 | FASTmrEMMA |
| 2 | 211393932 | 0.1247 | 9.75 | 10.6872 | 0.6045 | mrMLM | 5 | 40921864 | 0.3075 | 5.51 | 6.3254 | 0.3488 | FASTmrEMMA |
| 3 | 220509353 | -0.2918 | 17.22 | 18.2725 | 1.8379 | mrMLM | 5 | 150074309 | 0.2264 | 3.46 | 4.1852 | 0.4655 | FASTmrEMMA |
| 3 | 222753455 | 0.1035 | 4.59 | 5.3713 | 0.3783 | mrMLM | 5 | 179939948 | -0.2669 | 6.62 | 7.4685 | 0.5573 | FASTmrEMMA |
| 3 | 181564906 | -0.0915 | 3.79 | 4.5341 | 0.2992 | mrMLM | 6 | 97113705 | 0.355 | 8.22 | 9.1142 | 0.8974 | FASTmrEMMA |
| 3 | 29473154 | -0.1277 | 4.55 | 5.3279 | 0.4028 | mrMLM | 6 | 104865718 | -0.8046 | 18.64 | 19.7117 | 3.4443 | FASTmrEMMA |
| 3 | 222482863 | -0.1351 | 5.96 | 6.7957 | 0.5602 | mrMLM | 6 | 163018858 | -0.2897 | 4.59 | 5.3642 | 0.6121 | FASTmrEMMA |

| | | | | | | | | | | | | | |
|---|---|---|---|---|---|---|---|---|---|---|---|---|---|
| 4 | 216687458 | 0.1526 | 8.89 | 9.8035 | 0.6063 | mrMLM | 7 | 9198079 | -0.3048 | 8.09 | 8.9820 | 0.6776 | FASTmrEMMA |
| 5 | 188656939 | -0.1897 | 4.40 | 5.1658 | 0.6172 | mrMLM | 8 | 22170626 | -0.586 | 7.22 | 8.0886 | 0.9214 | FASTmrEMMA |
| 5 | 150074309 | 0.1039 | 4.60 | 5.3763 | 0.42 | mrMLM | 8 | 157465327 | -0.3783 | 13.62 | 14.6237 | 1.3356 | FASTmrEMMA |
| 5 | 51435319 | 0.1353 | 8.97 | 9.8811 | 0.6889 | mrMLM | 8 | 169138120 | -0.2304 | 4.50 | 5.2707 | 0.378 | FASTmrEMMA |
| 5 | 1362747 | -0.1493 | 9.35 | 10.2759 | 0.6602 | mrMLM | 8 | 171597499 | 0.1769 | 4.32 | 5.0838 | 0.295 | FASTmrEMMA |
| 5 | 193656423 | 0.1007 | 5.00 | 5.7928 | 0.3686 | mrMLM | 8 | 173031488 | 0.3357 | 7.58 | 8.4586 | 1.1015 | FASTmrEMMA |
| 6 | 104858202 | -0.2489 | 8.25 | 9.1473 | 1.1242 | mrMLM | 9 | 15614702 | -0.2808 | 7.49 | 8.3653 | 0.7429 | FASTmrEMMA |
| 7 | 173251838 | -0.8954 | 37.84 | 39.0638 | 5.132 | mrMLM | 9 | 153562715 | 0.3369 | 10.41 | 11.3599 | 1.0341 | FASTmrEMMA |
| 7 | 8846936 | 0.0863 | 6.54 | 7.3859 | 0.2828 | mrMLM | 10 | 24578257 | -0.4537 | 8.11 | 9.0032 | 0.9837 | FASTmrEMMA |
| 7 | 6146507 | -0.1329 | 6.03 | 6.8640 | 0.4554 | mrMLM | 10 | 26483664 | -0.5464 | 11.95 | 12.9252 | 1.7625 | FASTmrEMMA |
| 7 | 155392927 | 0.1477 | 9.08 | 9.9989 | 0.6005 | mrMLM | 10 | 102370515 | -0.2628 | 5.67 | 6.4864 | 0.5725 | FASTmrEMMA |
| 7 | 133531802 | -0.108 | 4.19 | 4.9516 | 0.2551 | mrMLM | 10 | 117214050 | -0.2643 | 5.41 | 6.2252 | 0.582 | FASTmrEMMA |
| 7 | 92391957 | -0.1302 | 6.56 | 7.4081 | 0.4095 | mrMLM | 1 | 7966534 | -0.3313 | 7.48 | 8.3550 | 0.0533 | pLARmEB |
| 8 | 125298375 | -0.4993 | 20.45 | 21.5345 | 1.494 | mrMLM | 1 | 8657361 | 0.1048 | 4.85 | 5.6438 | 0.036 | pLARmEB |
| 8 | 75605439 | -0.151 | 5.66 | 6.4783 | 0.3037 | mrMLM | 1 | 16349283 | -0.2684 | 6.26 | 7.1012 | 0.0791 | pLARmEB |
| 8 | 29146171 | -0.1522 | 7.14 | 8.0074 | 0.6325 | mrMLM | 1 | 17668962 | -0.2078 | 3.71 | 4.4454 | 0.0371 | pLARmEB |
| 8 | 111625183 | -0.1496 | 7.04 | 7.9051 | 0.6626 | mrMLM | 1 | 55071145 | -0.1667 | 3.98 | 4.7293 | 0.0379 | pLARmEB |
| 8 | 138882711 | -0.1112 | 8.15 | 9.0495 | 0.468 | mrMLM | 2 | 24390872 | -0.1069 | 5.44 | 6.2539 | 0.0446 | pLARmEB |
| 9 | 153836357 | -0.1498 | 11.89 | 12.8598 | 0.8055 | mrMLM | 2 | 149341223 | -0.4782 | 8.58 | 9.4854 | 0.0914 | pLARmEB |
| 9 | 153562715 | 0.1084 | 4.20 | 4.9619 | 0.4488 | mrMLM | 2 | 149517374 | -0.1826 | 4.42 | 5.1932 | 0.0515 | pLARmEB |
| 10 | 102319748 | -0.1384 | 8.57 | 9.4812 | 0.5323 | mrMLM | 2 | 212526845 | -0.87 | 12.53 | 13.5159 | 0.3677 | pLARmEB |
| 10 | 88614957 | -0.1147 | 5.11 | 5.9147 | 0.2557 | mrMLM | 3 | 171029218 | 0.1341 | 3.35 | 4.0672 | 0.0252 | pLARmEB |

| | | | | | | | | | | | | | |
|---|---|---|---|---|---|---|---|---|---|---|---|---|---|
| 1 | 162267119 | 0.0751 | 5.06 | 5.8547 | 0.1646 | FASTmrMLM | 3 | 202363278 | -0.5809 | 6.08 | 6.9167 | 0.2068 | pLARmEB |
| 1 | 171060219 | -0.1986 | 6.01 | 6.8460 | 0.316 | FASTmrMLM | 3 | 222596913 | 0.0668 | 3.34 | 4.0579 | 0.0111 | pLARmEB |
| 1 | 198416421 | 0.0982 | 8.33 | 9.2319 | 0.3687 | FASTmrMLM | 4 | 185387136 | -0.1498 | 3.73 | 4.4691 | 0.0453 | pLARmEB |
| 1 | 231186353 | 0.0603 | 4.33 | 5.0947 | 0.1376 | FASTmrMLM | 5 | 4769561 | -0.1229 | 6.73 | 7.5841 | 0.0546 | pLARmEB |
| 1 | 248149904 | 0.2619 | 7.78 | 8.6693 | 0.5496 | FASTmrMLM | 5 | 25615959 | 0.1039 | 3.50 | 4.2260 | 0.0208 | pLARmEB |
| 1 | 293269574 | -0.0795 | 4.37 | 5.1400 | 0.1526 | FASTmrMLM | 6 | 163012904 | -0.1388 | 8.99 | 9.9031 | 0.0657 | pLARmEB |
| 2 | 2551778 | -0.2423 | 6.43 | 7.2737 | 0.3518 | FASTmrMLM | 7 | 17889236 | 0.1112 | 4.13 | 4.8836 | 0.0493 | pLARmEB |
| 2 | 19283564 | -0.0675 | 4.46 | 5.2335 | 0.1532 | FASTmrMLM | 7 | 109329336 | -0.2769 | 6.11 | 6.9516 | 0.0782 | pLARmEB |
| 2 | 149517635 | -0.3293 | 14.17 | 15.1817 | 0.9115 | FASTmrMLM | 7 | 141513616 | 0.1239 | 4.89 | 5.6773 | 0.0388 | pLARmEB |
| 2 | 172555792 | -0.0807 | 5.93 | 6.7645 | 0.2504 | FASTmrMLM | 8 | 38489776 | -0.2909 | 3.45 | 4.1757 | 0.0829 | pLARmEB |
| 2 | 183922016 | -0.1204 | 11.09 | 12.0471 | 0.4927 | FASTmrMLM | 8 | 100960678 | -0.5827 | 12.32 | 13.3057 | 0.2782 | pLARmEB |
| 2 | 193735059 | 0.1194 | 5.77 | 6.5930 | 0.3819 | FASTmrMLM | 8 | 142177238 | -0.4392 | 8.99 | 9.9023 | 0.0603 | pLARmEB |
| 2 | 212636843 | 0.0783 | 6.60 | 7.4532 | 0.2347 | FASTmrMLM | 8 | 157463010 | 0.1253 | 6.46 | 7.3045 | 0.0401 | pLARmEB |
| 2 | 212744513 | -0.0918 | 5.70 | 6.5183 | 0.2988 | FASTmrMLM | 8 | 169138120 | -0.1091 | 3.81 | 4.5512 | 0.0347 | pLARmEB |
| 3 | 150169183 | -0.0874 | 8.99 | 9.9018 | 0.248 | FASTmrMLM | 9 | 10320276 | -0.475 | 10.36 | 11.3056 | 0.0902 | pLARmEB |
| 3 | 217633424 | 0.0802 | 4.59 | 5.3655 | 0.2513 | FASTmrMLM | 9 | 15614693 | -0.0602 | 3.68 | 4.4146 | 0.014 | pLARmEB |
| 3 | 221689683 | 0.0881 | 8.10 | 8.9990 | 0.2173 | FASTmrMLM | 9 | 17648206 | -0.1434 | 4.77 | 5.5576 | 0.0409 | pLARmEB |
| 4 | 62868426 | -0.1606 | 5.01 | 5.8115 | 0.227 | FASTmrMLM | 9 | 26601762 | 0.1785 | 8.59 | 9.4979 | 0.0633 | pLARmEB |
| 5 | 4769561 | -0.0726 | 5.26 | 6.0706 | 0.1863 | FASTmrMLM | 9 | 93146938 | 0.1198 | 5.40 | 6.2102 | 0.0415 | pLARmEB |
| 5 | 30995288 | -0.0523 | 3.98 | 4.7340 | 0.104 | FASTmrMLM | 1 | 248149904 | 0.2631 | 3.89 | 4.6399 | 1.5504 | pKWmEB |
| 5 | 171274922 | 0.2473 | 4.69 | 5.4779 | 0.1078 | FASTmrMLM | 1 | 268158349 | -0.1328 | 4.01 | 4.7611 | 0.322 | pKWmEB |
| 5 | 196508497 | 0.0678 | 3.21 | 3.9190 | 0.0923 | FASTmrMLM | 1 | 292354425 | 0.1453 | 4.59 | 5.3731 | 0.9693 | pKWmEB |

| | | | | | | | | | | | | | |
|---|---|---|---|---|---|---|---|---|---|---|---|---|---|
| 6 | 104865718 | -0.2528 | 18.25 | 19.3195 | 1.3604 | FASTmrMLM | 1 | 162267119 | 0.152 | 5.32 | 6.1251 | 0.717 | pKWmEB |
| 7 | 5138788 | -0.1107 | 6.10 | 6.9316 | 0.4547 | FASTmrMLM | 2 | 149517374 | -0.1973 | 3.47 | 4.1975 | 0.9209 | pKWmEB |
| 7 | 17889236 | 0.0967 | 3.91 | 4.6575 | 0.365 | FASTmrMLM | 2 | 2805975 | 0.116 | 3.23 | 3.9420 | 0.955 | pKWmEB |
| 7 | 141513616 | 0.0675 | 3.23 | 3.9423 | 0.1125 | FASTmrMLM | 4 | 5306611 | 0.2592 | 7.14 | 8.0042 | 0.887 | pKWmEB |
| 7 | 173251838 | -0.4825 | 13.80 | 14.8026 | 1.49 | FASTmrMLM | 5 | 4769561 | -0.1628 | 7.90 | 8.7899 | 0.9587 | pKWmEB |
| 8 | 21818669 | -0.1363 | 4.10 | 4.8576 | 0.2132 | FASTmrMLM | 5 | 75890500 | -0.1166 | 4.50 | 5.2793 | 0.6919 | pKWmEB |
| 8 | 26347033 | -0.2665 | 10.95 | 11.9060 | 0.5973 | FASTmrMLM | 6 | 104865718 | -0.1752 | 3.74 | 4.4825 | 1.7929 | pKWmEB |
| 8 | 38412621 | -0.2182 | 8.90 | 9.8175 | 0.6345 | FASTmrMLM | 8 | 141649337 | 0.2186 | 9.09 | 10.0127 | 2.388 | pKWmEB |
| 8 | 118681100 | 0.1413 | 6.00 | 6.8291 | 0.2292 | FASTmrMLM | 8 | 26346491 | -0.4393 | 5.05 | 5.8529 | 2.938 | pKWmEB |
| 8 | 125317148 | 0.0892 | 3.85 | 4.5919 | 0.0823 | FASTmrMLM | 8 | 21818669 | -0.1941 | 3.08 | 3.7769 | 1.5137 | pKWmEB |
| 8 | 148482834 | 0.2478 | 10.03 | 10.9702 | 0.3678 | FASTmrMLM | 8 | 160374255 | -0.2746 | 7.28 | 8.1484 | 0.8709 | pKWmEB |
| 8 | 158369778 | 0.1012 | 9.49 | 10.4222 | 0.3971 | FASTmrMLM | 10 | 23787517 | -0.8471 | 12.79 | 13.7840 | 8.9586 | pKWmEB |
| 8 | 160374255 | -0.3099 | 18.86 | 19.9356 | 0.9202 | FASTmrMLM | 10 | 16487724 | -0.2464 | 3.41 | 4.1351 | 1.2385 | pKWmEB |
| 9 | 3477249 | 0.0691 | 4.70 | 5.4858 | 0.1436 | FASTmrMLM | 10 | 141946713 | -0.2304 | 3.71 | 4.4495 | 0.9887 | pKWmEB |
| 9 | 15614702 | -1.00E-04 | 3.02 | 3.7152 | 4.12E-07 | FASTmrMLM | 6 | 102199028 | -0.4623 | 4.82 | 5.6067 | 1.455 | ISIS EM-BLASSO |
| 9 | 133900817 | -0.0518 | 3.35 | 4.0666 | 0.0909 | FASTmrMLM | 6 | 104865718 | -0.3231 | 7.51 | 8.3865 | 2.2225 | ISIS EM-BLASSO |
| 9 | 153836357 | -0.0364 | 3.68 | 4.4130 | 0.0475 | FASTmrMLM | 7 | 109329336 | -0.3232 | 3.64 | 4.3757 | 1.0412 | ISIS EM-BLASSO |
| 10 | 23787517 | -0.6508 | 21.92 | 23.0190 | 2.7114 | FASTmrMLM | 8 | 26346491 | -0.859 | 14.16 | 15.1713 | 5.3208 | ISIS EM-BLASSO |
| 10 | 45433753 | 0.1734 | 9.21 | 10.1282 | 0.4541 | FASTmrMLM | 8 | 100960678 | -0.5971 | 4.86 | 5.6475 | 2.8563 | ISIS EM-BLASSO |
| 10 | 102366558 | -0.0553 | 3.93 | 4.6768 | 0.0898 | FASTmrMLM | 10 | 23787517 | -0.91 | 8.79 | 9.7053 | 5.3004 | ISIS EM-BLASSO |

**Table S6**. Comparison of seed oil related genes in maize identified by the software mrMLM in this study with those in Li et al. (2013)

| Chr | QTN position (bp) | GWAS | | | | Comparative genomics analysis | | | |
|---|---|---|---|---|---|---|---|---|---|
| | | Effect | LOD or P-value | $r^2$ (%) | Method † | Locus name | Gene and its functional annotation | Distance (kb) ‡ | Reference |
| QTNs detected simultaneously by the software mrMLM and Li et al. (2013) Nat Genet 45(1):43-50 | | | | | | | | | |
| 1 | 16349283 | -0.27 | 6.26 | 0.08 | 4 | *GRMZM2G080524* | Epoxide hydrolase, *EH* | 21 | [1,2] |
| 1 | 248149904 | 0.26 | 7.78 | 0.55 | 2 | *GRMZM2G110298* | Acyl carrier protein, *ACP* | 0 | [3-5] |
| 2 | 53680401 | -0.18 | 5.96 | 0.53 | 3 | *GRMZM2G134432* | Phosphatidylinosito 3 kinase, *PI3Ks.a* | 678 | [3,6] |
| 2 | 149341223~149517635 | -0.48~-0.18 | 4.42~14.17 | 0.05~0.91 | 2,4 | *GRMZM2G079236* | Long-chain Acyl-CoA synthetase, *LACS* | 0 | [3,7,8] |
| 4 | 6601755 | -0.49 | 6.04 | 0.84 | 3 | *GRMZM2G099666* | Acyl-coenzyme A oxidase, *ACX2* | 193 | |
| 6 | 104858202~104865718 | -0.80~-0.32 | 3.74~18.25 | 1.12~3.44 | 1~3,5,6 | *GRMZM2G169089* | Diglyceride acyltransferase, *DGAT1-2* | 0 | [3,9~11] |
| 7 | 109329336 | -0.32~-0.28 | 3.64~6.11 | 0.08~1.04 | 4,6 | *GRMZM2G092550* | Phosphatidylinositol 3 kinase, *PI3Ks.b* | 4 | [3,6] |
| 8 | 38412621~38489776 | -0.29~-0.22 | 3.45~8.90 | 0.08~0.63 | 2,4 | *GRMZM2G003022* | COPII-coated vesicles, *COPII* | 31 | [3,12] |
| 10 | 16487724 | -0.25 | 3.41 | 1.23 | 5 | *GRMZM2G169240* | Fatty acid desaturase-1, *FAD2* | 2582 | [3,9,13~16] |
| 10 | 117214050 | -0.2643 | 5.41 | 0.58 | 3 | *GRMZM5G828253* | Oxidoreductase activity, cytochrome P450, *CYPOR* | 313 | [17,18] |
| QTNs detected only by the software mrMLM, candidate genes were identified by key words like: fatty acid, oil and triacylglycerol biosynthesis in the MaizeGDB (https://www.maizegdb.org/). | | | | | | | | | |
| 1 | 13536744 | -0.12 | 4.56 | 0.43 | 1 | *GRMZM2G031790* | 3-ketoacyl-CoA synthase 2, *KCS2* | 589 | [19] |
| 1 | 296768029 | -0.20 | 4.10 | 0.46 | 1 | *GRMZM2G369815* | Seed fatty acid reducer 4, *SFAR4* | 609 | [20] |
| 2 | 211393932 | 0.12 | 9.75 | 0.60 | 1 | *GRMZM2G124335* | Fatty acid biosynthesis 1, *FAB1* | 2 | [21] |
| 2 | 24388940 | -0.22 | 4.54 | 0.46 | 3 | *GRMZM2G078373* | Sphingolipid delta4-desaturase, *DES-1-LIKE* | 840 | [22] |
| 4 | 216687458 | 0.15 | 8.89 | 0.61 | 1 | *GRMZM2G091715* | Acyl carrier protein 4, *ACP4* | 19 | [23] |
| 5 | 1362747 | -0.15 | 9.35 | 0.66 | 1 | *GRMZM2G019866* | Acyl carrier protein 1, *ACP1* | 548 | [24] |
| 6 | 97113705 | 0.36 | 8.22 | 0.90 | 3 | *GRMZM2G322892* | Triacylglycerol lipase 2, *LIP2* | 268 | [25] |
| 6 | 102199028 | -0.46 | 4.82 | 1.46 | 6 | *GRMZM2G079308* | Fatty acyl-ACP thioesterases B, *FATB* | 950 | [26] |
| 7 | 8846936 | 0.09 | 6.54 | 0.28 | 1 | *GRMZM2G174766* | Fatty acid desaturase 2, *FAD2* | 928 | [27] |
| 7 | 141513616 | 0.07~0.12 | 3.23~4.89 | 0.04~0.11 | 2,4 | *GRMZM2G020740* | 3-ketoacyl-CoA synthase 4, *KCS4* | 502 | [28] |
| 7 | 92391957 | -0.13 | 6.56 | 0.41 | 1 | *GRMZM2G129453* | Delta(8)-fatty-acid desaturase 2, *SLD2* | 549 | [29] |
| 9 | 10320276 | -0.48 | 10.36 | 0.09 | 4 | *GRMZM2G012863* | 3-ketoacyl-acyl carrier protein synthase I, *KASI* | 897 | [30] |
| 9 | 17648206 | -0.14 | 4.77 | 0.04 | 4 | *GRMZM5G864319* | Peroxisomal acyl-coenzyme A oxidase 1, *ACX1* | 327 | |
| QTNs detected only by Li et al. (2013) Nat Genet 45(1):43-50 | | | | | | | | | |
| 3 | 166664152 | | 4.6e-07 | | | *GRMZM2G176542* | Triglyceride lipases, *TAGL* | 0 | [3,31~33] |
| 3 | 167431166 | | 1.1e-06 | | | *GRMZM2G118423* | Oxidoreductase activity, Cytochrome P450, *CYPOR* | 45 | [17,18] |
| 3 | 178136002 | | 8.5e-07 | | | *GRMZM2G083195* | Glycerol-phosphate acyltransferase, *GPAT* | 46 | [3,34~36] |
| 4 | 32810884 | | 9.4e-07 | | | *GRMZM5G847159* | Oxidoreductase activity, cytochrome P450, *CYPOR* | 1 | [17,18] |

†: 1, 2, 3, 4, 5, and 6 represent mrMLM, FASTmrMLM, FASTmrEMMA, pLARmEB, pKWmEB, and ISIS EM-BLASSO, respectively. ‡: distance (kb) between QTNs and gene.

### References

1 Nawrath C. The Biopolymers Cutin and Suberin. Arabidopsis Book 2002; 1: e0021. DOI: 10.1199/tab.0021

2 Li-Beisson Y, Shorrosh B, Beisson F, Andersson MX, Arondel V, Bates PD, et al. Acyl-Lipid Metabolism. Arabidopsis Book 2010; 8: e0133. DOI: 10.1199/tab.0133

3 Safford R, Windust JH, Lucas C, De Silva J, James CM, Hellyer A, et al. Plastid–localised seed acyl–carrier protein of *Brassica napus* is encoded by a distinct, nuclear multigene family. Eur J Biochem 1988; 174: 287–95.

4 Evans DE, Taylor PE, Singh MB, Knox RB. The interrelationship between the accumulation of lipids, protein and the level of acyl carrier protein during the development of *Brassica napus* L. pollen. Planta 1992;186: 343–54.

5 Kurz EU, Lees-Miller SP. DNA damage-induced activation of ATM and ATM-dependent signaling pathways. DNA Repair (Amst) 2004;3:889-900.

6 Shockey JM, Fulda MS, Browse JA. *Arabidopsis* contains nine long–chain acyl–coenzyme a synthetase genes that participate in fatty acid and glycerolipid metabolism. Plant Physiol 2002;129:1710-22 .

7 Fulda M, Shockey J, Werber M, Wolter FP, Heinz E. Two long-chain acyl-CoA synthetases from *Arabidopsis thaliana* involved in peroxisomal fatty acid beta-oxidation. Plant J 2002;32: 93-103.

8 Beló, A, Zheng P, Luck S, Shen B, Meyer DJ, Li B, et al. Whole genome scan detects an allelic variant of *fad2* associated with increased oleic acid levels in maize. Mol Genet Genomics 2008;279:1–10.

9 Oakes J, Brackenridge D, Colletti R, Daley M, Hawkins DJ, Xiong H, et al. Expression of fungal *diacylglycerol acyltransferase2* genes to increase kernel oil in maize. Plant Physiol 2011;155:1146–57.

10 Xu J, Francis T, Mietkiewska E, Giblin EM, Barton DL, Zhang Y, et al. Cloning and characterization of an acyl–CoA–dependent *diacylglycerol acyltransferase 1 (DGAT1)* gene from Tropaeolum majus, and a study of the functional motifs of the DGAT protein using site–directed mutagenesis to modify enzyme activity and oil content. Plant Biotechnol J 2008;6:799–818.

11 Nickel W, Brugger B, Wieland FT. Protein and lipid sorting between the endoplasmic reticulum and the Golgi complex. Semin Cell Dev Biol 1998;9:493–501.

12 Mikkilineni V, Rocheford TR. Sequence variation and genomic organization of fatty acid desaturase–2 (fad2) and fatty acid desaturase–6 (fad6) cDNAs in maize. Theor Appl Genet 2003;106:1326–32.

13 Wassom JJ, Mikkelineni V, Bohn MO, Rocheford TR. QTL for fatty acid composition of maize kernel oil in Illinois high oil × B73 backcross–derived lines. Crop Sci 2007;48:69–78.

14 Okuley J, Lightner J, Feldmann K, Yadav N, Lark E, Browse J. *Arabidopsis FAD2* gene encodes the enzyme that is essential for polyunsaturated lipid synthesis. Plant Cell 1994;6:147–158.

15 Dyer JM, Mullen RT. Immunocytological localization of two plant fatty acid desaturases in the endoplasmic reticulum. FEBS Lett 2001;494:44–47.

16 Benveniste I, Tijet N, Adas F, Philipps G, Salaün JP, Durst F. *CYP86A1* from *Arabidopsis thaliana* encodes a cytochrome P450-dependent fatty acid omega–hydroxylase. Biochem Bioph Res Commun 1998;243:688–93.

17 Song WC, Funk CD, Brash AR. Molecular cloning of an allene oxide synthase: a cytochrome P450 specialized for the metabolism of fatty acid hydroperoxides. Proc Natl Acad Sci U S A 1993;90:8519–23.

18 Lee SB, Jung SJ, Go YS, Kim HU, Kim JK, Cho HJ, et al. Two *Arabidopsis* 3-ketoacyl CoA synthase genes, *KCS20* and *KCS2/DAISY*, are functionally redundant in cuticular wax and root suberin biosynthesis, but differentially controlled by osmotic stress. Plant J 2009; 60:462–75.

19 Huang LM, Lai CP, Chen LO, Chan MT, Shaw JF. *Arabidopsis SFAR4* is a novel GDSL-type esterase involved in fatty acid degradation and glucose tolerance. Bot Stud 2015;56:33. DOI: 10.1186/s40529-015- 0114-6.

20 Hirano T, Sato MH. Diverse physiological functions of FAB1 and phosphatidylinositol 3,5-bisphosphate in plants. Front Plant Sci 2019;10:274.

21 Michaelson LV, Zäuner S, Markham JE, Haslam RP, Desikan R, Mugford S, et al. Functional characterization of a higher plant sphingolipid Delta4-desaturase: defining the role of sphingosine and sphingosine-1-phosphate in Arabidopsis. Plant Physiol 2009;149:487–98.

22 Huang J, Xue C, Wang H, Wang L, Schmidt W, Shen R, et al. Genes of acyl carrier protein family show different expression profiles and overexpression of acyl carrier protein 5 modulates fatty acid composition and enhances salt stress tolerance in *Arabidopsis*. Front Plant Sci 2017;8:987.

23 Tong X, Oh EK, Lee BH, Lee JK. Production of long-chain free fatty acids from metabolically engineered *Rhodobacter sphaeroides* heterologously producing periplasmic phospholipase A2 in dodecane-overlaid two-phase culture. Microb Cell Fact 2019;18:20.

24 EI-Kouhen K, Blangy S, Ortiz E, Gardies AM, Ferté N, Arondel V. Identification and characterization of a triacylglycerol lipase *in Arabidopsis* homologous to mammalian acid lipases. FEBS Lett 2005;579:6067–73.

25 Bonaventure G, Salas JJ, Pollard MR, Ohlrogge JB. Disruption of the *FATB* gene in *Arabidopsis* demonstrates an essential role of saturated fatty acids in plant growth. Plant Cell 2003;15:1020–

33.

26 Nguyen VC, Nakamura Y, Kanehara K. Membrane lipid polyunsaturation mediated by *fatty acid desaturase 2* (*FAD2*) is involved in endoplasmic reticulum stress tolerance in *Arabidopsis thaliana*. Plant J 2019;99:478–93.

27 Paul S, Gable K, Beaudoin F, Cahoon E, Jaworski J, Napier JA, et al. Members of the *Arabidopsis* FAE1-like 3-ketoacyl-CoA synthase gene family substitute for the Elop proteins of *Saccharomyces cerevisiae*. J Biol Chem 2006;281:9018–29.

28 Chen M, Markham JE, Cahoon EB. Sphingolipid Δ8 unsaturation is important for glucosylceramide biosynthesis and low-temperature performance *in Arabidopsis*. Plant J 2012;69:769–81.

29 Ding W, Lin L, Zhang B, Xiang X, Wu J, Pan Z, et al. *OsKASI,* a β-ketoacyl-[acyl carrier protein] synthase I, is involved in root development in rice (*Oryza sativa* L.). Planta 2015;242:203–13.

30 Hellyer SA, Chandler IC, Bosley JA. Can the fatty acid selectivity of plant lipases be predicted from the composition of the seed triglyceride? Biochim Biophys Acta 1999;1440:215–24.

31 Rosnitschek I, Theimer RR. Properties of a membrane–bound triglyceride lipase of rapeseed (*Brassica napus* L.) cotyledons. Planta 1980;148:193–8.

32 Paloccia C, Soro S, Cernia E, Fiorillo F, Belsito CMA, Monacelli B. Lipolytic isoenzymes from *Euphorbia latex*. Plant Sci 2003;165:577–82.

33 Murata N, Tasaka Y. Glycerol-3-phosphate acyltransferase in plants. Biochim Biophys Acta 1997;1348:10–6.

34 Yang WL, Pollard M, Li-Beisson Y, Beisson F, Feig M, Ohlrogge J. A distinct type of glycerol–3–phosphate acyltransferase with sn–2 preference and phosphatase activity producing 2–monoacylglycerol. Proc Natl Acad Sci U S A 2010;107:12040–45.

35 Tamada T, Feese MD, Ferri SR, Kato Y, Yajima R, Toguri T, et al. Substrate recognition and selectivity of plant glycerol-3-phosphate acyltransferases (GPATs) from *Cucurbita moscata* and *Spinacea oleracea*. Acta Crystallogr D Biol Crystallogr 2004;60:13–21.

36 Fernández ME, Prando A, Rogberg-Muñoz A, Peral-García P, Baldo A, Giovambattista G, et al. Association of a region of bovine chromosome 1 (BTA1) with age at puberty in Angus bulls. Reproduction, Fertility and Development 2016; 28: 1618–21.

**Table S7**. All the QTNs for kidney weight in Simmental beef cattle detected by mrMLM, FASTmrMLM, FASTmrEMMA, pLARmEB, pKWmEB and ISIS EBLASSO

| Chr | Position (bp) | QTN effect | LOD score | $-\log_{10}(P)$ | $r^2$ (%) | Method | Chr | Position (bp) | QTN effect | LOD score | $-\log_{10}(P)$ | $r^2$ (%) | Method |
|---|---|---|---|---|---|---|---|---|---|---|---|---|---|
| 10 | 87848389 | -0.0539 | 7.03 | 7.9 | 6.5537 | mrMLM | 23 | 21702537 | -0.0034 | 14.1 | 15.11 | 0.0022 | FASTmrEMMA |
| 11 | 33921550 | 0.0305 | 3.76 | 4.5 | 2.3121 | mrMLM | 23 | 24692423 | -0.0051 | 4.02 | 4.77 | 0.0085 | FASTmrEMMA |
| 19 | 26569324 | 0.0367 | 3.12 | 3.82 | 2.9988 | mrMLM | 24 | 2010941 | -0.0025 | 3.24 | 3.95 | 0.0011 | FASTmrEMMA |
| 26 | 36522514 | 0.0395 | 3.27 | 3.99 | 3.8983 | mrMLM | 24 | 9740339 | 8.00E-04 | 3.19 | 3.9 | 2.00E-04 | FASTmrEMMA |
| 1 | 106358966 | 0.019 | 4.76 | 5.55 | 0.8435 | FASTmrMLM | 24 | 25697647 | -7.00E-04 | 3.12 | 3.82 | 1.69E-05 | FASTmrEMMA |
| 1 | 110649323 | -0.0268 | 5.79 | 6.61 | 1.6839 | FASTmrMLM | 24 | 25768771 | -7.00E-04 | 3.14 | 3.84 | 1.52E-05 | FASTmrEMMA |
| 2 | 2254298 | 0.0187 | 5.96 | 6.8 | 0.8176 | FASTmrMLM | 24 | 25951960 | -3.00E-04 | 4.16 | 4.92 | 1.62E-06 | FASTmrEMMA |
| 2 | 18246330 | 0.0107 | 3.12 | 3.83 | 0.2469 | FASTmrMLM | 24 | 58552772 | 9.00E-04 | 13.31 | 14.31 | 1.80E-05 | FASTmrEMMA |
| 3 | 115147732 | 0.0204 | 8.2 | 9.1 | 0.436 | FASTmrMLM | 25 | 7598966 | 0.0078 | 4.75 | 5.54 | 0.0211 | FASTmrEMMA |
| 4 | 2835171 | -0.0131 | 7.15 | 8.02 | 0.3461 | FASTmrMLM | 26 | 21564772 | 0.0099 | 18.28 | 19.35 | 0.0293 | FASTmrEMMA |
| 4 | 43362940 | 0.0221 | 5.99 | 6.82 | 0.9088 | FASTmrMLM | 26 | 27826727 | -0.0024 | 4.71 | 5.5 | 5.00E-04 | FASTmrEMMA |
| 4 | 61093205 | 0.0139 | 3.8 | 4.54 | 0.2639 | FASTmrMLM | 27 | 3852629 | 0.0011 | 6.67 | 7.52 | 4.00E-04 | FASTmrEMMA |
| 4 | 111413395 | 0.0155 | 4.71 | 5.49 | 0.5003 | FASTmrMLM | 27 | 3853699 | 0.0014 | 8.75 | 9.66 | 7.00E-04 | FASTmrEMMA |
| 4 | 120615269 | -0.0201 | 3.98 | 4.73 | 0.3849 | FASTmrMLM | 27 | 14236927 | -8.00E-04 | 8.83 | 9.74 | 1.86E-05 | FASTmrEMMA |
| 5 | 13463428 | 0.0029 | 6.14 | 6.98 | 0.0163 | FASTmrMLM | 27 | 16625147 | -0.0019 | 5.31 | 6.12 | 6.00E-04 | FASTmrEMMA |
| 5 | 19436069 | -0.0123 | 3.56 | 4.29 | 0.3455 | FASTmrMLM | 27 | 16832909 | -0.0012 | 3.36 | 4.08 | 1.00E-04 | FASTmrEMMA |
| 5 | 120338298 | -0.0094 | 3.09 | 3.79 | 0.1952 | FASTmrMLM | 27 | 16834042 | -0.0015 | 6.47 | 7.32 | 3.00E-04 | FASTmrEMMA |
| 6 | 21123441 | -0.032 | 6.6 | 7.45 | 1.159 | FASTmrMLM | 27 | 18546318 | -0.005 | 5.26 | 6.07 | 0.0092 | FASTmrEMMA |
| 6 | 25471760 | 0.0105 | 5.08 | 5.88 | 0.1698 | FASTmrMLM | 27 | 22739277 | -0.0024 | 3.02 | 3.71 | 0.002 | FASTmrEMMA |
| 6 | 42110373 | 0.0067 | 3.65 | 4.39 | 0.0349 | FASTmrMLM | 27 | 22757505 | -0.0035 | 4.23 | 5 | 0.0041 | FASTmrEMMA |
| 7 | 20296966 | -0.0225 | 4.77 | 5.55 | 0.8617 | FASTmrMLM | 27 | 38287694 | 0.003 | 12.16 | 13.14 | 0.0021 | FASTmrEMMA |
| 7 | 72721038 | -0.022 | 3.98 | 4.73 | 1.1011 | FASTmrMLM | 27 | 42890396 | -0.0024 | 4.43 | 5.2 | 8.00E-04 | FASTmrEMMA |
| 8 | 27592207 | -0.0165 | 4.83 | 5.62 | 0.553 | FASTmrMLM | 27 | 43733075 | 0.0019 | 7.44 | 8.32 | 0.001 | FASTmrEMMA |
| 8 | 107120944 | 0.0258 | 3.82 | 4.56 | 1.0643 | FASTmrMLM | 28 | 3532094 | -0.0029 | 5.1 | 5.9 | 0.001 | FASTmrEMMA |

| | | | | | | | | | | | | | |
|---|---|---|---|---|---|---|---|---|---|---|---|---|---|
| 10 | 5526827 | 0.0195 | 4.35 | 5.12 | 0.6004 | FASTmrMLM | 29 | 17566806 | 0.0062 | 5.67 | 6.49 | 0.0129 | FASTmrEMMA |
| 10 | 21317870 | -0.0336 | 5.84 | 6.66 | 1.5648 | FASTmrMLM | 29 | 49511085 | 0.0151 | 4.33 | 5.1 | 0.0772 | FASTmrEMMA |
| 10 | 87848389 | -0.0373 | 10 | 10.94 | 2.9108 | FASTmrMLM | 1 | 9427932 | 0.0088 | 5.72 | 6.54 | 0.1315 | pLARmEB |
| 10 | 90978450 | -0.0184 | 8.59 | 9.5 | 0.7477 | FASTmrMLM | 1 | 34164923 | -0.0085 | 3.84 | 4.59 | 0.1793 | pLARmEB |
| 11 | 50973309 | -0.0016 | 3.87 | 4.62 | 0.0052 | FASTmrMLM | 1 | 65752195 | 0.001 | 4.98 | 5.77 | 0.0014 | pLARmEB |
| 11 | 66048270 | 0.003 | 3.38 | 4.1 | 0.0189 | FASTmrMLM | 1 | 120987043 | 0.0032 | 9.00 | 9.91 | 0.0206 | pLARmEB |
| 11 | 91627517 | 0.0186 | 3.25 | 3.96 | 0.2778 | FASTmrMLM | 1 | 133085461 | 0.0036 | 4.53 | 5.30 | 0.0339 | pLARmEB |
| 12 | 30043755 | 4.00E-04 | 4.13 | 4.89 | 4.00E-04 | FASTmrMLM | 1 | 155322206 | -0.0034 | 8.02 | 8.92 | 0.0312 | pLARmEB |
| 12 | 59135006 | -0.0303 | 5.01 | 5.8 | 1.8107 | FASTmrMLM | 2 | 11903885 | 0.0051 | 13.03 | 14.03 | 0.0703 | pLARmEB |
| 12 | 84524504 | 0.0253 | 10.2 | 11.14 | 1.4713 | FASTmrMLM | 2 | 20169565 | 0.0011 | 6.37 | 7.21 | 7.00E-04 | pLARmEB |
| 13 | 42320076 | -0.0243 | 6.4 | 7.25 | 1.2289 | FASTmrMLM | 2 | 52417495 | -7.00E-04 | 15.21 | 16.24 | 2.00E-04 | pLARmEB |
| 13 | 60570356 | 0.0257 | 4.47 | 5.24 | 1.4059 | FASTmrMLM | 3 | 25794439 | -2.04E-05 | 4.34 | 5.11 | 1.34E-07 | pLARmEB |
| 14 | 10829335 | 0.0137 | 3.02 | 3.71 | 0.2039 | FASTmrMLM | 3 | 35135984 | 0.0113 | 6.37 | 7.22 | 0.322 | pLARmEB |
| 14 | 21917782 | 0.0079 | 3.29 | 4.01 | 0.0856 | FASTmrMLM | 3 | 37282673 | 0.0028 | 9.11 | 10.03 | 0.0188 | pLARmEB |
| 15 | 8576846 | 0.0233 | 4.6 | 5.38 | 1.2459 | FASTmrMLM | 3 | 51935982 | 6.00E-04 | 6.07 | 6.90 | 5.00E-04 | pLARmEB |
| 15 | 68472538 | -5.00E-04 | 3.85 | 4.6 | 7.00E-04 | FASTmrMLM | 3 | 55957304 | 0.0021 | 4.91 | 5.71 | 0.0029 | pLARmEB |
| 16 | 14813526 | -0.0271 | 4.09 | 4.85 | 1.7082 | FASTmrMLM | 3 | 57416268 | -0.0024 | 6.19 | 7.03 | 0.007 | pLARmEB |
| 17 | 32625001 | 0.0329 | 5.17 | 5.98 | 2.2679 | FASTmrMLM | 3 | 93556351 | -0.0037 | 7.99 | 8.88 | 0.0356 | pLARmEB |
| 17 | 51676255 | 0.0187 | 5.79 | 6.62 | 0.6449 | FASTmrMLM | 3 | 120403408 | 0.005 | 8.97 | 9.89 | 0.0592 | pLARmEB |
| 18 | 38317858 | 0.0269 | 5.64 | 6.46 | 1.6866 | FASTmrMLM | 4 | 2845004 | -0.0052 | 8.42 | 9.33 | 0.0707 | pLARmEB |
| 18 | 54604727 | -0.0027 | 3.3 | 4.02 | 0.0151 | FASTmrMLM | 4 | 5006344 | 6.00E-04 | 7.06 | 7.93 | 3.00E-04 | pLARmEB |
| 19 | 60859466 | -0.0131 | 5.24 | 6.04 | 0.4042 | FASTmrMLM | 4 | 26662279 | -0.0011 | 13.27 | 14.26 | 0.0015 | pLARmEB |
| 20 | 44864240 | 0.0288 | 9.71 | 10.64 | 1.9296 | FASTmrMLM | 4 | 30681746 | 0.004 | 10.39 | 11.34 | 0.0257 | pLARmEB |
| 22 | 21951237 | -0.0139 | 5.09 | 5.89 | 0.3223 | FASTmrMLM | 4 | 47490095 | -9.00E-04 | 3.27 | 3.98 | 9.00E-04 | pLARmEB |
| 23 | 14902969 | -0.0241 | 4.95 | 5.74 | 1.3311 | FASTmrMLM | 4 | 50267564 | 0.0043 | 15.45 | 16.48 | 0.017 | pLARmEB |
| 23 | 21336942 | 0.0564 | 3.23 | 3.93 | 2.2647 | FASTmrMLM | 4 | 55527318 | -2.00E-04 | 39.35 | 40.57 | 1.25E-05 | pLARmEB |

| | | | | | | | | | | | | | |
|---|---|---|---|---|---|---|---|---|---|---|---|---|---|
| 24 | 45436667 | -0.0076 | 4.61 | 5.39 | 0.1349 | FASTmrMLM | 4 | 89075410 | 0.0037 | 21.17 | 22.27 | 0.0282 | pLARmEB |
| 26 | 21564772 | 0.0231 | 5.2 | 6 | 1.2334 | FASTmrMLM | 4 | 111413395 | 0.0087 | 7.20 | 8.07 | 0.1873 | pLARmEB |
| 26 | 26369697 | -0.0201 | 8.78 | 9.7 | 0.5686 | FASTmrMLM | 4 | 118912501 | -7.00E-04 | 13.98 | 14.99 | 1.00E-04 | pLARmEB |
| 27 | 792244 | 0.0099 | 3.54 | 4.26 | 0.1449 | FASTmrMLM | 5 | 15061346 | 0.0066 | 12.50 | 13.49 | 0.0665 | pLARmEB |
| 27 | 8476228 | 0.0101 | 3.36 | 4.08 | 0.2049 | FASTmrMLM | 5 | 15116972 | 0.005 | 14.83 | 15.86 | 0.037 | pLARmEB |
| 27 | 39546339 | 0.0132 | 4.65 | 5.43 | 0.3908 | FASTmrMLM | 5 | 23903481 | 0.0043 | 7.19 | 8.06 | 0.0454 | pLARmEB |
| 28 | 34780664 | 0.0224 | 5.56 | 6.38 | 1.1451 | FASTmrMLM | 5 | 38190841 | -0.0023 | 18.36 | 19.43 | 0.0146 | pLARmEB |
| 28 | 36504079 | 9.04E-05 | 3.52 | 4.25 | 1.37E-05 | FASTmrMLM | 5 | 47042381 | 4.00E-04 | 4.53 | 5.31 | 3.00E-04 | pLARmEB |
| 29 | 43539616 | 0.0193 | 5.25 | 6.05 | 0.8177 | FASTmrMLM | 5 | 48121153 | 0.0063 | 5.37 | 6.18 | 0.1051 | pLARmEB |
| 1 | 12953453 | -0.0023 | 7.6 | 8.48 | 0.0012 | FASTmrEMMA | 5 | 67286371 | -0.0011 | 10.12 | 11.06 | 4.00E-04 | pLARmEB |
| 1 | 27730401 | 8.00E-04 | 3.29 | 4.01 | 1.47E-05 | FASTmrEMMA | 5 | 68166725 | 0.0031 | 7.18 | 8.05 | 0.0013 | pLARmEB |
| 1 | 110520102 | 0.0013 | 3.33 | 4.05 | 4.00E-04 | FASTmrEMMA | 5 | 104413139 | 2.00E-04 | 8.07 | 8.97 | 1.00E-04 | pLARmEB |
| 1 | 133085461 | 0.006 | 6.2 | 7.04 | 0.0078 | FASTmrEMMA | 5 | 118127379 | 0.0013 | 6.26 | 7.10 | 0.0035 | pLARmEB |
| 1 | 133086487 | 0.0053 | 5.22 | 6.02 | 0.0062 | FASTmrEMMA | 5 | 118397277 | 4.00E-04 | 10.62 | 11.57 | 3.00E-04 | pLARmEB |
| 1 | 138305770 | -0.0014 | 7.19 | 8.06 | 1.00E-04 | FASTmrEMMA | 6 | 21123441 | -0.0064 | 9.49 | 10.42 | 0.0543 | pLARmEB |
| 1 | 141823000 | 5.00E-04 | 9.31 | 10.23 | 4.07E-06 | FASTmrEMMA | 6 | 39387542 | 0.0026 | 12.00 | 12.98 | 0.0057 | pLARmEB |
| 1 | 141826766 | 5.00E-04 | 9.58 | 10.51 | 5.70E-06 | FASTmrEMMA | 6 | 39410541 | 0.005 | 25.96 | 27.10 | 0.0426 | pLARmEB |
| 1 | 148538451 | -5.00E-04 | 6.1 | 6.93 | 5.71E-06 | FASTmrEMMA | 6 | 44967444 | 0.0012 | 14.16 | 15.17 | 0.0039 | pLARmEB |
| 1 | 150311512 | -0.0032 | 3.44 | 4.16 | 0.0028 | FASTmrEMMA | 6 | 72068201 | -0.0015 | 4.25 | 5.02 | 0.0049 | pLARmEB |
| 1 | 150317595 | -0.0029 | 4.11 | 4.87 | 0.0023 | FASTmrEMMA | 7 | 28834853 | -9.00E-04 | 5.89 | 6.72 | 9.00E-04 | pLARmEB |
| 1 | 155312956 | -4.00E-04 | 10.42 | 11.37 | 2.30E-06 | FASTmrEMMA | 7 | 53107671 | 0.0032 | 7.77 | 8.65 | 0.028 | pLARmEB |
| 2 | 18246330 | 0.01 | 8.3 | 9.19 | 0.0333 | FASTmrEMMA | 7 | 71033098 | -0.003 | 8.24 | 9.14 | 0.0185 | pLARmEB |
| 2 | 46723266 | -0.001 | 5.77 | 6.6 | 2.00E-04 | FASTmrEMMA | 7 | 83599948 | -0.001 | 17.26 | 18.31 | 3.00E-04 | pLARmEB |
| 2 | 54235770 | 0.0415 | 11.71 | 12.68 | 0.407 | FASTmrEMMA | 7 | 89744240 | -0.0018 | 3.02 | 3.72 | 0.0032 | pLARmEB |
| 2 | 81187013 | -3.00E-04 | 3 | 3.7 | 9.00E-07 | FASTmrEMMA | 7 | 98954416 | 8.00E-04 | 19.50 | 20.58 | 8.00E-04 | pLARmEB |
| 2 | 85037178 | 0.004 | 6.2 | 7.04 | 0.0046 | FASTmrEMMA | 8 | 27592207 | -0.0106 | 3.80 | 4.54 | 0.2705 | pLARmEB |

| | | | | | | | | | | | | | |
|---|---|---|---|---|---|---|---|---|---|---|---|---|---|
| 2 | 136271704 | 0.0028 | 5.3 | 6.11 | 0.0011 | FASTmrEMMA | 8 | 38000683 | -0.0014 | 25.25 | 26.39 | 0.0022 | pLARmEB |
| 2 | 136535374 | 0.0022 | 4.81 | 5.6 | 7.00E-04 | FASTmrEMMA | 9 | 41431307 | 0.0022 | 7.01 | 7.87 | 0.0061 | pLARmEB |
| 3 | 2245496 | -0.001 | 3.14 | 3.85 | 1.94E-05 | FASTmrEMMA | 9 | 82856321 | -0.0018 | 16.51 | 17.55 | 0.0079 | pLARmEB |
| 3 | 22281672 | -0.0043 | 3.98 | 4.73 | 0.0056 | FASTmrEMMA | 10 | 10622667 | 0.0039 | 15.35 | 16.38 | 0.0214 | pLARmEB |
| 3 | 32247139 | -8.00E-04 | 10.52 | 11.47 | 6.49E-05 | FASTmrEMMA | 10 | 21317870 | -0.0093 | 4.74 | 5.52 | 0.1408 | pLARmEB |
| 3 | 32260522 | -0.0117 | 5.07 | 5.86 | 0.0472 | FASTmrEMMA | 10 | 87848389 | -0.0165 | 11.05 | 12.01 | 0.6724 | pLARmEB |
| 3 | 35135984 | 0.0042 | 4.27 | 5.04 | 0.0038 | FASTmrEMMA | 10 | 94890549 | 0.0016 | 6.54 | 7.39 | 9.00E-04 | pLARmEB |
| 3 | 35570341 | 0.004 | 10.39 | 11.34 | 0.0034 | FASTmrEMMA | 11 | 105397592 | 0.0047 | 8.18 | 9.07 | 0.0274 | pLARmEB |
| 3 | 35635939 | -0.001 | 13.5 | 14.5 | 2.10E-05 | FASTmrEMMA | 12 | 8001704 | 0.0011 | 22.41 | 23.52 | 0.0032 | pLARmEB |
| 3 | 60755786 | -0.0015 | 6.18 | 7.02 | 2.00E-04 | FASTmrEMMA | 12 | 45303004 | -0.0051 | 6.63 | 7.48 | 0.0482 | pLARmEB |
| 3 | 80430626 | -0.0047 | 7.2 | 8.07 | 0.0067 | FASTmrEMMA | 12 | 52455037 | -7.00E-04 | 8.34 | 9.24 | 2.00E-04 | pLARmEB |
| 3 | 90262349 | 0.0013 | 4.6 | 5.38 | 2.00E-04 | FASTmrEMMA | 12 | 56168000 | -8.00E-04 | 9.08 | 10.00 | 2.00E-04 | pLARmEB |
| 3 | 93556351 | -0.0016 | 5.33 | 6.13 | 9.00E-04 | FASTmrEMMA | 12 | 59135006 | -0.0075 | 4.83 | 5.62 | 0.1318 | pLARmEB |
| 3 | 101850519 | 5.00E-04 | 3.34 | 4.06 | 5.02E-06 | FASTmrEMMA | 12 | 81211856 | 0.0011 | 4.56 | 5.34 | 0.0014 | pLARmEB |
| 3 | 115147732 | 0.0067 | 8.16 | 9.06 | 0.0143 | FASTmrEMMA | 13 | 10102104 | 0.0068 | 16.85 | 17.89 | 0.1196 | pLARmEB |
| 4 | 5589375 | -4.00E-04 | 3.24 | 3.95 | 2.11E-06 | FASTmrEMMA | 13 | 39533788 | 0.0022 | 5.06 | 5.86 | 0.0136 | pLARmEB |
| 4 | 5767201 | 0.0036 | 3.8 | 4.54 | 0.0021 | FASTmrEMMA | 13 | 43482413 | -7.00E-04 | 6.34 | 7.18 | 1.00E-04 | pLARmEB |
| 4 | 11587232 | 0.0038 | 6.17 | 7.01 | 0.002 | FASTmrEMMA | 13 | 68196929 | 0.0026 | 3.78 | 4.52 | 0.0185 | pLARmEB |
| 4 | 23580709 | -0.0089 | 6.08 | 6.91 | 0.0296 | FASTmrEMMA | 14 | 7154835 | 0.0019 | 4.41 | 5.18 | 0.005 | pLARmEB |
| 4 | 31233839 | -0.0013 | 3.43 | 4.15 | 4.00E-04 | FASTmrEMMA | 14 | 59442036 | 0.0013 | 12.47 | 13.45 | 7.00E-04 | pLARmEB |
| 4 | 31790534 | -0.0062 | 10.89 | 11.84 | 0.0101 | FASTmrEMMA | 14 | 80350162 | 0.0073 | 8.46 | 9.37 | 0.1345 | pLARmEB |
| 4 | 72633007 | 5.00E-04 | 3.58 | 4.31 | 3.38E-06 | FASTmrEMMA | 15 | 8835116 | -0.0016 | 4.66 | 5.44 | 0.0016 | pLARmEB |
| 4 | 89058238 | 0.0055 | 9.03 | 9.94 | 0.0101 | FASTmrEMMA | 15 | 32268220 | -0.001 | 5.31 | 6.12 | 7.00E-04 | pLARmEB |
| 4 | 89075410 | 0.0061 | 7.99 | 8.89 | 0.0132 | FASTmrEMMA | 15 | 40521052 | -0.0015 | 10.09 | 11.03 | 0.0038 | pLARmEB |
| 4 | 104775809 | -0.0012 | 5.97 | 6.81 | 8.05E-05 | FASTmrEMMA | 15 | 48963568 | 9.00E-04 | 7.34 | 8.22 | 8.00E-04 | pLARmEB |
| 4 | 106194302 | -0.0019 | 5.19 | 5.99 | 3.00E-04 | FASTmrEMMA | 16 | 1595222 | -0.0025 | 6.04 | 6.87 | 0.0062 | pLARmEB |

| | | | | | | | | | | | | | |
|---|---|---|---|---|---|---|---|---|---|---|---|---|---|
| 4 | 111413395 | 0.0056 | 4.09 | 4.84 | 0.0102 | FASTmrEMMA | 16 | 4452514 | -0.0043 | 9.01 | 9.93 | 0.0298 | pLARmEB |
| 4 | 118912501 | -6.00E-04 | 4.08 | 4.84 | 7.46E-06 | FASTmrEMMA | 16 | 14813526 | -0.0102 | 16.91 | 17.96 | 0.2855 | pLARmEB |
| 5 | 1559287 | 6.00E-04 | 9.66 | 10.59 | 9.33E-05 | FASTmrEMMA | 16 | 17770213 | -9.00E-04 | 10.24 | 11.18 | 4.00E-04 | pLARmEB |
| 5 | 13075314 | -0.0012 | 3.59 | 4.32 | 5.00E-04 | FASTmrEMMA | 16 | 23268022 | 0.0012 | 4.16 | 4.92 | 2.00E-04 | pLARmEB |
| 5 | 38001237 | -0.0028 | 5.01 | 5.81 | 0.0027 | FASTmrEMMA | 16 | 59383104 | 2.00E-04 | 4.24 | 5.00 | 2.68E-05 | pLARmEB |
| 5 | 88048251 | 7.00E-04 | 7.5 | 8.38 | 2.00E-04 | FASTmrEMMA | 16 | 60496060 | -7.00E-04 | 3.23 | 3.94 | 4.00E-04 | pLARmEB |
| 5 | 88054351 | 3.00E-04 | 6.47 | 7.32 | 2.48E-05 | FASTmrEMMA | 16 | 60942210 | 0.0068 | 10.17 | 11.11 | 0.1089 | pLARmEB |
| 5 | 112293257 | 2.00E-04 | 4.13 | 4.89 | 6.92E-06 | FASTmrEMMA | 16 | 69766481 | -0.0023 | 3.08 | 3.78 | 0.014 | pLARmEB |
| 5 | 118399856 | 6.00E-04 | 8.75 | 9.66 | 2.02E-05 | FASTmrEMMA | 16 | 72291455 | 0.0023 | 3.41 | 4.13 | 0.009 | pLARmEB |
| 6 | 21105453 | -0.0054 | 8.1 | 8.99 | 0.0083 | FASTmrEMMA | 17 | 32625001 | 0.0058 | 25.36 | 26.49 | 0.0825 | pLARmEB |
| 6 | 39410541 | 0.0031 | 14.92 | 15.94 | 0.0011 | FASTmrEMMA | 17 | 32661304 | 0.0022 | 9.99 | 10.93 | 0.0129 | pLARmEB |
| 6 | 71870305 | -0.0011 | 5.35 | 6.16 | 4.35E-05 | FASTmrEMMA | 17 | 69892739 | 8.00E-04 | 7.67 | 8.55 | 2.00E-04 | pLARmEB |
| 6 | 72076018 | -0.0027 | 4.24 | 5 | 0.0012 | FASTmrEMMA | 18 | 13016845 | 4.00E-04 | 3.82 | 4.56 | 1.00E-04 | pLARmEB |
| 6 | 72841953 | 0.0025 | 10.19 | 11.13 | 8.00E-04 | FASTmrEMMA | 18 | 25754843 | -0.0107 | 3.62 | 4.35 | 0.2923 | pLARmEB |
| 6 | 100620519 | -8.00E-04 | 9.32 | 10.25 | 1.54E-05 | FASTmrEMMA | 18 | 38317858 | 0.0089 | 35.11 | 36.32 | 0.2164 | pLARmEB |
| 6 | 100626375 | -9.00E-04 | 5.56 | 6.38 | 1.75E-05 | FASTmrEMMA | 18 | 39961400 | 0.0051 | 4.23 | 5.00 | 0.0534 | pLARmEB |
| 6 | 113157379 | -0.0036 | 3.1 | 3.81 | 0.0034 | FASTmrEMMA | 19 | 33155236 | 7.00E-04 | 16.55 | 17.59 | 1.00E-04 | pLARmEB |
| 7 | 30916696 | -9.00E-04 | 8.8 | 9.72 | 7.45E-05 | FASTmrEMMA | 19 | 52265051 | 0.003 | 5.65 | 6.47 | 0.0217 | pLARmEB |
| 7 | 36236133 | -0.001 | 3.38 | 4.1 | 3.00E-04 | FASTmrEMMA | 19 | 57708681 | -8.48E-05 | 4.58 | 5.36 | 1.84E-06 | pLARmEB |
| 7 | 36268647 | -0.0014 | 3.77 | 4.51 | 6.00E-04 | FASTmrEMMA | 20 | 10328240 | -5.00E-04 | 10.75 | 11.70 | 5.30E-05 | pLARmEB |
| 7 | 53124827 | 0.0021 | 12.75 | 13.73 | 0.0011 | FASTmrEMMA | 20 | 24932795 | -0.0015 | 4.64 | 5.42 | 0.0047 | pLARmEB |
| 7 | 65539778 | 3.00E-04 | 6.12 | 6.96 | 1.12E-06 | FASTmrEMMA | 20 | 37391207 | -0.0013 | 3.94 | 4.68 | 0.003 | pLARmEB |
| 7 | 70532694 | -8.00E-04 | 3.85 | 4.59 | 1.16E-05 | FASTmrEMMA | 20 | 71511933 | -0.006 | 3.52 | 4.25 | 0.0542 | pLARmEB |
| 7 | 70779399 | -5.00E-04 | 3.41 | 4.13 | 8.15E-06 | FASTmrEMMA | 21 | 5370856 | -0.0061 | 8.32 | 9.22 | 0.0876 | pLARmEB |
| 7 | 70816154 | -5.00E-04 | 4.81 | 5.6 | 7.13E-06 | FASTmrEMMA | 21 | 27386386 | -0.0013 | 3.98 | 4.73 | 5.00E-04 | pLARmEB |
| 8 | 8522529 | 0.0011 | 3.4 | 4.11 | 2.00E-04 | FASTmrEMMA | 21 | 31277031 | 0.0028 | 15.41 | 16.44 | 0.0205 | pLARmEB |

| | | | | | | | | | | | | | |
|---|---|---|---|---|---|---|---|---|---|---|---|---|---|
| 8 | 38200914 | 0.0027 | 3.8 | 4.55 | 0.0011 | FASTmrEMMA | 21 | 45767354 | -0.0038 | 4.63 | 5.41 | 0.0242 | pLARmEB |
| 8 | 38210062 | 0.0024 | 5.94 | 6.78 | 8.00E-04 | FASTmrEMMA | 21 | 69841951 | 5.00E-04 | 9.18 | 10.11 | 6.20E-05 | pLARmEB |
| 8 | 40422559 | -0.0015 | 5 | 5.79 | 7.00E-04 | FASTmrEMMA | 22 | 43700547 | 0.0013 | 14.64 | 15.66 | 0.0046 | pLARmEB |
| 8 | 40425857 | 4.00E-04 | 5.47 | 6.28 | 4.33E-05 | FASTmrEMMA | 22 | 56907006 | 0.0059 | 20.97 | 22.07 | 0.0464 | pLARmEB |
| 8 | 54150455 | -5.00E-04 | 4.02 | 4.77 | 6.41E-06 | FASTmrEMMA | 22 | 57693714 | 0.001 | 6.61 | 7.46 | 0.001 | pLARmEB |
| 8 | 98559973 | 8.00E-04 | 8.34 | 9.24 | 2.13E-05 | FASTmrEMMA | 22 | 59128692 | 0.0047 | 4.29 | 5.06 | 0.0299 | pLARmEB |
| 9 | 26042975 | -0.0014 | 3.32 | 4.04 | 3.00E-04 | FASTmrEMMA | 24 | 25951960 | -4.00E-04 | 8.21 | 9.11 | 3.54E-05 | pLARmEB |
| 9 | 32079319 | -6.00E-04 | 8.67 | 9.58 | 1.17E-05 | FASTmrEMMA | 24 | 27838377 | -0.0073 | 10.64 | 11.59 | 0.111 | pLARmEB |
| 9 | 85391712 | 0.0034 | 3.42 | 4.14 | 0.0037 | FASTmrEMMA | 24 | 34515639 | 0.0045 | 3.24 | 3.95 | 0.0068 | pLARmEB |
| 9 | 98048267 | 0.0048 | 7.33 | 8.2 | 0.006 | FASTmrEMMA | 26 | 27651115 | -0.001 | 3.07 | 3.77 | 0.0026 | pLARmEB |
| 10 | 55918765 | 0.0013 | 3.16 | 3.87 | 6.00E-04 | FASTmrEMMA | 27 | 16835142 | -0.0031 | 9.50 | 10.43 | 0.0124 | pLARmEB |
| 10 | 56798705 | -6.00E-04 | 6.84 | 7.7 | 7.02E-06 | FASTmrEMMA | 27 | 22604390 | -0.005 | 5.97 | 6.80 | 0.067 | pLARmEB |
| 10 | 57724034 | 0.035 | 4.21 | 4.98 | 0.2061 | FASTmrEMMA | 27 | 37937057 | -0.0033 | 4.11 | 4.87 | 0.0243 | pLARmEB |
| 10 | 94890549 | 8.00E-04 | 4.81 | 5.6 | 1.30E-05 | FASTmrEMMA | 27 | 42241046 | 0.0021 | 11.29 | 12.25 | 0.0113 | pLARmEB |
| 10 | 102677040 | 0.016 | 3.93 | 4.68 | 0.0629 | FASTmrEMMA | 29 | 49713592 | -0.0058 | 3.65 | 4.38 | 0.0923 | pLARmEB |
| 11 | 48806854 | -0.0074 | 6.94 | 7.8 | 0.0185 | FASTmrEMMA | 2 | 52417495 | -0.0013 | 3.31 | 4.02 | 4.849 | pKWmEB |
| 11 | 48808618 | -0.0081 | 6.17 | 7.01 | 0.0218 | FASTmrEMMA | 3 | 6582830 | -0.0284 | 3.25 | 3.96 | 2.3517 | pKWmEB |
| 11 | 87229489 | -6.00E-04 | 4.35 | 5.12 | 3.43E-05 | FASTmrEMMA | 5 | 87945266 | 0.0266 | 5.06 | 5.86 | 1.3354 | pKWmEB |
| 11 | 103285126 | -0.0031 | 4.36 | 5.13 | 0.0034 | FASTmrEMMA | 7 | 98954416 | 0.0261 | 3.44 | 4.16 | 2.2349 | pKWmEB |
| 11 | 103289035 | -0.0058 | 3.43 | 4.16 | 0.0115 | FASTmrEMMA | 12 | 2694960 | -0.0225 | 3.23 | 3.94 | 3.7222 | pKWmEB |
| 11 | 105397592 | 0.0224 | 3.32 | 4.04 | 0.1571 | FASTmrEMMA | 15 | 68722924 | -0.0269 | 3.09 | 3.79 | 4.6118 | pKWmEB |
| 11 | 106085417 | -0.0017 | 4.11 | 4.86 | 2.00E-04 | FASTmrEMMA | 16 | 14557973 | 0.025 | 3.32 | 4.03 | 1.7466 | pKWmEB |
| 12 | 2693770 | -0.0034 | 6.54 | 7.39 | 0.0033 | FASTmrEMMA | 23 | 21336942 | 0.0174 | 3.25 | 3.96 | 5.0578 | pKWmEB |
| 12 | 2694960 | -0.005 | 4.22 | 4.98 | 0.006 | FASTmrEMMA | 1 | 9427932 | 0.0232 | 3.43 | 4.16 | 0.7465 | ISIS EM-BLASSO |
| 12 | 2706809 | -0.0044 | 3.71 | 4.45 | 0.0058 | FASTmrEMMA | 1 | 98943363 | -0.0291 | 4.29 | 5.06 | 1.359 | ISIS EM-BLASSO |
| 12 | 45303004 | -0.0019 | 3.99 | 4.74 | 5.00E-04 | FASTmrEMMA | 2 | 46721161 | -0.0342 | 6.5 | 7.35 | 2.3939 | ISIS EM-BLASSO |

| | | | | | | | | | | | | | |
|---|---|---|---|---|---|---|---|---|---|---|---|---|---|
| 12 | 52534782 | -2.00E-04 | 3.96 | 4.71 | 6.90E-07 | FASTmrEMMA | 3 | 32260522 | -0.0236 | 5.66 | 6.48 | 0.8113 | ISIS EM-BLASSO |
| 12 | 57008805 | 0.0032 | 3.28 | 4 | 0.0016 | FASTmrEMMA | 3 | 44060152 | 0.0255 | 4.85 | 5.64 | 1.4004 | ISIS EM-BLASSO |
| 12 | 78378824 | -0.0044 | 6.21 | 7.05 | 0.0065 | FASTmrEMMA | 3 | 79036558 | 0.0207 | 3.51 | 4.24 | 0.9599 | ISIS EM-BLASSO |
| 12 | 84524504 | 0.006 | 5.99 | 6.83 | 0.0085 | FASTmrEMMA | 3 | 120936971 | 0.0221 | 4.48 | 5.26 | 1.0596 | ISIS EM-BLASSO |
| 12 | 84530638 | 0.0046 | 9.96 | 10.89 | 0.0048 | FASTmrEMMA | 4 | 23622809 | -0.0231 | 4.79 | 5.57 | 1.0662 | ISIS EM-BLASSO |
| 13 | 2914469 | 0.0015 | 7.57 | 8.45 | 4.00E-04 | FASTmrEMMA | 4 | 82581345 | 0.0203 | 3.35 | 4.06 | 0.8658 | ISIS EM-BLASSO |
| 13 | 39533788 | 0.005 | 4.16 | 4.92 | 0.0068 | FASTmrEMMA | 4 | 101451691 | 0.0246 | 3.03 | 3.73 | 1.241 | ISIS EM-BLASSO |
| 13 | 59620161 | -0.0017 | 3.52 | 4.25 | 0.001 | FASTmrEMMA | 4 | 103251884 | 0.0252 | 5.87 | 6.7 | 1.3963 | ISIS EM-BLASSO |
| 13 | 60014103 | -7.00E-04 | 6.08 | 6.92 | 8.30E-06 | FASTmrEMMA | 5 | 19436069 | -0.0246 | 6.66 | 7.52 | 1.3361 | ISIS EM-BLASSO |
| 13 | 60574380 | 0.0066 | 6.89 | 7.75 | 0.0108 | FASTmrEMMA | 5 | 87945266 | 0.0279 | 8.29 | 9.19 | 0.7976 | ISIS EM-BLASSO |
| 13 | 72588925 | -0.0015 | 3.74 | 4.48 | 2.00E-04 | FASTmrEMMA | 5 | 120339577 | -0.0345 | 7.86 | 8.75 | 2.618 | ISIS EM-BLASSO |
| 14 | 49785058 | 7.00E-04 | 5.52 | 6.34 | 1.34E-05 | FASTmrEMMA | 7 | 98954416 | 0.0304 | 4.18 | 4.94 | 0.9864 | ISIS EM-BLASSO |
| 15 | 17658204 | -0.0026 | 3.28 | 3.99 | 0.0013 | FASTmrEMMA | 8 | 27592207 | -0.0243 | 6.9 | 7.76 | 1.15 | ISIS EM-BLASSO |
| 15 | 68722924 | -0.0024 | 3.43 | 4.15 | 7.00E-04 | FASTmrEMMA | 8 | 30093176 | -0.0179 | 3.44 | 4.16 | 0.3918 | ISIS EM-BLASSO |
| 16 | 1595222 | -0.0013 | 4.51 | 5.28 | 1.00E-04 | FASTmrEMMA | 10 | 57724034 | 0.028 | 4.67 | 5.45 | 0.1219 | ISIS EM-BLASSO |
| 16 | 11506312 | -0.002 | 5.27 | 6.08 | 0.001 | FASTmrEMMA | 10 | 60006026 | 0.0155 | 4.01 | 4.76 | 0.5108 | ISIS EM-BLASSO |
| 16 | 14813526 | -0.0089 | 4.81 | 5.6 | 0.022 | FASTmrEMMA | 10 | 87848389 | -0.0307 | 4.46 | 5.23 | 1.8986 | ISIS EM-BLASSO |
| 16 | 26192996 | 0.0055 | 6.78 | 7.64 | 0.0105 | FASTmrEMMA | 11 | 13638961 | 0.0134 | 3.07 | 3.77 | 0.3922 | ISIS EM-BLASSO |
| 16 | 55784564 | -0.005 | 3.54 | 4.26 | 0.0075 | FASTmrEMMA | 11 | 105557304 | 0.0261 | 3.65 | 4.38 | 1.1643 | ISIS EM-BLASSO |
| 16 | 58558880 | -0.0043 | 3.58 | 4.31 | 0.006 | FASTmrEMMA | 12 | 2694960 | -0.028 | 5.67 | 6.5 | 1.6648 | ISIS EM-BLASSO |
| 16 | 58565978 | -0.0043 | 3.68 | 4.42 | 0.0064 | FASTmrEMMA | 12 | 57008805 | 0.0294 | 4.53 | 5.3 | 1.4951 | ISIS EM-BLASSO |
| 16 | 61791136 | -0.0055 | 10.15 | 11.1 | 0.0077 | FASTmrEMMA | 13 | 10102104 | 0.0173 | 3.06 | 3.76 | 0.6406 | ISIS EM-BLASSO |
| 16 | 73840001 | 0.0024 | 11.64 | 12.61 | 0.0017 | FASTmrEMMA | 13 | 60570356 | 0.0217 | 3.38 | 4.1 | 0.9572 | ISIS EM-BLASSO |
| 16 | 73859947 | 3.00E-04 | 14.72 | 15.74 | 2.08E-05 | FASTmrEMMA | 14 | 61307574 | 0.0234 | 3.49 | 4.22 | 1.1752 | ISIS EM-BLASSO |
| 17 | 52566415 | -0.0024 | 7.11 | 7.98 | 0.001 | FASTmrEMMA | 15 | 42998912 | -0.0208 | 4.24 | 5.01 | 0.9563 | ISIS EM-BLASSO |
| 17 | 53620065 | 0.0018 | 6.04 | 6.87 | 0.001 | FASTmrEMMA | 15 | 61936585 | 0.0162 | 3.24 | 3.95 | 0.5773 | ISIS EM-BLASSO |

| | | | | | | | | | | | | | |
|---|---|---|---|---|---|---|---|---|---|---|---|---|---|
| 17 | 53755199 | 0.0014 | 4.16 | 4.92 | 4.00E-04 | FASTmrEMMA | 16 | 14813526 | -0.023 | 4.17 | 4.93 | 1.1822 | ISIS EM-BLASSO |
| 17 | 60241679 | -0.0054 | 7.38 | 8.26 | 0.0094 | FASTmrEMMA | 16 | 26192996 | 0.0182 | 3.91 | 4.65 | 0.5214 | ISIS EM-BLASSO |
| 18 | 38315068 | 0.0043 | 25.58 | 26.71 | 0.0057 | FASTmrEMMA | 17 | 19876360 | 0.0277 | 4.38 | 5.15 | 1.4099 | ISIS EM-BLASSO |
| 18 | 38317858 | 0.0075 | 32.76 | 33.95 | 0.0174 | FASTmrEMMA | 17 | 43400994 | 0.0211 | 3.78 | 4.52 | 0.8969 | ISIS EM-BLASSO |
| 18 | 39908826 | 0.0018 | 8.6 | 9.5 | 5.00E-04 | FASTmrEMMA | 17 | 51676255 | 0.0138 | 3.19 | 3.9 | 0.3373 | ISIS EM-BLASSO |
| 18 | 39961400 | 0.0014 | 6.2 | 7.04 | 3.00E-04 | FASTmrEMMA | 17 | 53620065 | 0.023 | 3.94 | 4.69 | 1.143 | ISIS EM-BLASSO |
| 18 | 51399522 | 0.0013 | 4.66 | 5.44 | 6.64E-05 | FASTmrEMMA | 18 | 41347362 | -0.0044 | 4.08 | 4.84 | 0.0325 | ISIS EM-BLASSO |
| 19 | 57385189 | -0.0019 | 8.68 | 9.58 | 3.00E-04 | FASTmrEMMA | 20 | 44864240 | 0.0062 | 4.67 | 5.45 | 0.0868 | ISIS EM-BLASSO |
| 20 | 24932795 | -0.0035 | 5.25 | 6.06 | 0.0021 | FASTmrEMMA | 21 | 30978657 | -0.0257 | 6.59 | 7.44 | 0.7558 | ISIS EM-BLASSO |
| 20 | 37391207 | -0.0017 | 8.67 | 9.57 | 4.00E-04 | FASTmrEMMA | 22 | 8023152 | 0.015 | 4.09 | 4.85 | 0.4859 | ISIS EM-BLASSO |
| 20 | 52784792 | -0.0021 | 3.32 | 4.04 | 6.00E-04 | FASTmrEMMA | 22 | 21951237 | -0.0145 | 3.01 | 3.7 | 0.3374 | ISIS EM-BLASSO |
| 21 | 30983811 | -0.0058 | 3.44 | 4.16 | 0.0114 | FASTmrEMMA | 24 | 27838377 | -0.0365 | 5.49 | 6.3 | 2.2718 | ISIS EM-BLASSO |
| 21 | 45748303 | -0.0024 | 3.14 | 3.84 | 0.001 | FASTmrEMMA | 25 | 16334342 | -0.026 | 5.57 | 6.39 | 1.3828 | ISIS EM-BLASSO |
| 21 | 45767354 | -0.002 | 3.6 | 4.33 | 5.00E-04 | FASTmrEMMA | 27 | 8476228 | 0.0157 | 4.71 | 5.49 | 0.4761 | ISIS EM-BLASSO |
| 22 | 19073643 | -9.00E-04 | 6.21 | 7.05 | 3.52E-05 | FASTmrEMMA | 27 | 22604390 | -0.0187 | 3.06 | 3.76 | 0.7701 | ISIS EM-BLASSO |
| 22 | 21953136 | 8.00E-04 | 4.37 | 5.14 | 2.00E-04 | FASTmrEMMA | 27 | 38286763 | -0.0327 | 8.73 | 9.64 | 0.8199 | ISIS EM-BLASSO |
| 22 | 43829445 | 0.0056 | 4.05 | 4.8 | 0.0087 | FASTmrEMMA | 27 | 43737541 | 0.0216 | 3.61 | 4.34 | 0.5826 | ISIS EM-BLASSO |
| 22 | 56906352 | 9.00E-04 | 3.46 | 4.18 | 7.32E-05 | FASTmrEMMA | 28 | 45890977 | 0.0226 | 4.85 | 5.64 | 0.9327 | ISIS EM-BLASSO |
| 22 | 57693714 | 0.0013 | 6.65 | 7.5 | 1.00E-04 | FASTmrEMMA | 29 | 49511085 | 0.0224 | 5.53 | 6.34 | 1.0817 | ISIS EM-BLASSO |
| 23 | 13428133 | 0.0076 | 14.6 | 15.62 | 0.0191 | FASTmrEMMA | | | | | | | |

**Table S8**. Previously reported genes for kidney weight in Simmental beef cattle around the QTNs identified by our multi-locus GWAS methods

| Chr | Position (bp) | Multi-locus GWAS | | | | Comparative genomics analysis | | |
|---|---|---|---|---|---|---|---|---|
| | | QTN effect | LOD score | $r^2$ (%) | Method | Candidate gene and its functional annotation | Distance (kb) ‡ | Reference |
| 1 | 98943363 | -0.0291 | 4.29 | 1.3590 | 6 | MDS1 and EVI1 complex locus; *MECOM* | 147 | [1] |
| 6 | 39387542 | 0.0026 | 12.0 | 0.0057 | 4 | non-SMC condensin I complex subunit G; *NCAPG* | 575 | [2] |
| 6 | 39410541 | 0.0031~0.0050 | 14.92~25.96 | 0.0011~0.0426 | 3,4 | ligand dependent nuclear receptor corepressor like; *LCORL* | 418 | [2] |

†: 1, 2, 3, 4, 5, and 6 represent mrMLM, FASTmrMLM, FASTmrEMMA, pLARmEB, pKWmEB, and ISIS EM-BLASSO, respectively. ‡: distance (kb) between QTNs and gene.

#### References


1 Fernández ME, Prando A, Rogberg-Muñoz A, Peral-García P, Baldo A, Giovambattista G, et al. Association of a region of bovine chromosome 1 (BTA1) with age at puberty in Angus bulls. Reproduction, Fertility and Development 2016; 28: 1618–21.

2 Zhang W, Li J, Guo Y, Zhang L, Xu L, Gao X, et al. Multi-strategy genome-wide association studies identify the DCAF16-NCAPG region as a susceptibility locus for average daily gain in cattle. Sci Rep 2016;6:38073

### Supplementary material D. Monte Carlo simulation experiments

All the simulation datasets used in this study were downloaded from the Dryad Digital Repository (http://dx.doi.org/10.5061/dryad.sk652) [1] and re-analyzed to validate the software package. Here we simply described them. Sample size was 199, the number of markers was 10000, six QTNs were simulated on marker positions, and the number of replicates was 1000. For each simulated QTN, we counted the samples in which the LOD scores exceeded 3.0 for our multi-locus methods, and the P-value was less than 0.05/$m$ for others. In the first, second and third simulation experiments, the phenotypic values were simulated by the models $\mathbf{y}=\boldsymbol{\mu}+\sum_{k=1}^{6}\mathbf{x}_k b_k+\boldsymbol{\varepsilon}$, $\mathbf{y}=\boldsymbol{\mu}+\sum_{k=1}^{6}\mathbf{x}_k b_k+\mathbf{u}+\boldsymbol{\varepsilon}$ and $\mathbf{y}=\boldsymbol{\mu}+\sum_{k=1}^{6}\mathbf{x}_k b_k+\sum_{l=1}^{3}(\mathrm{A}_l\#\mathrm{B}_l)b_{ll}+\boldsymbol{\varepsilon}$, respectively, where polygenic effect $\mathbf{u}\sim\mathrm{MVN}_n(0,2\times\mathbf{K})$, residual effect $\boldsymbol{\varepsilon}\sim\mathrm{MVN}_n(0,10\times\mathbf{I})$, $b_k$ is QTN effect ($k=1,\cdots,6$), $b_{ll}$ is epistatic effect ($l=1,\cdots,3$), $\mathrm{A}_l\#\mathrm{B}_l$ is the incidence coefficient of epistatic effect, and $\mathbf{K}$ is the kinship matrix between a pair of individuals.

#### References


1 Zhang J, Feng JY, Ni YL, Wen YJ, Niu Y, Tamba CL, et al. pLARmEB: integration of least angle regression with empirical Bayes for multilocus genome-wide association studies. Heredity 2017;**118:**517–24.

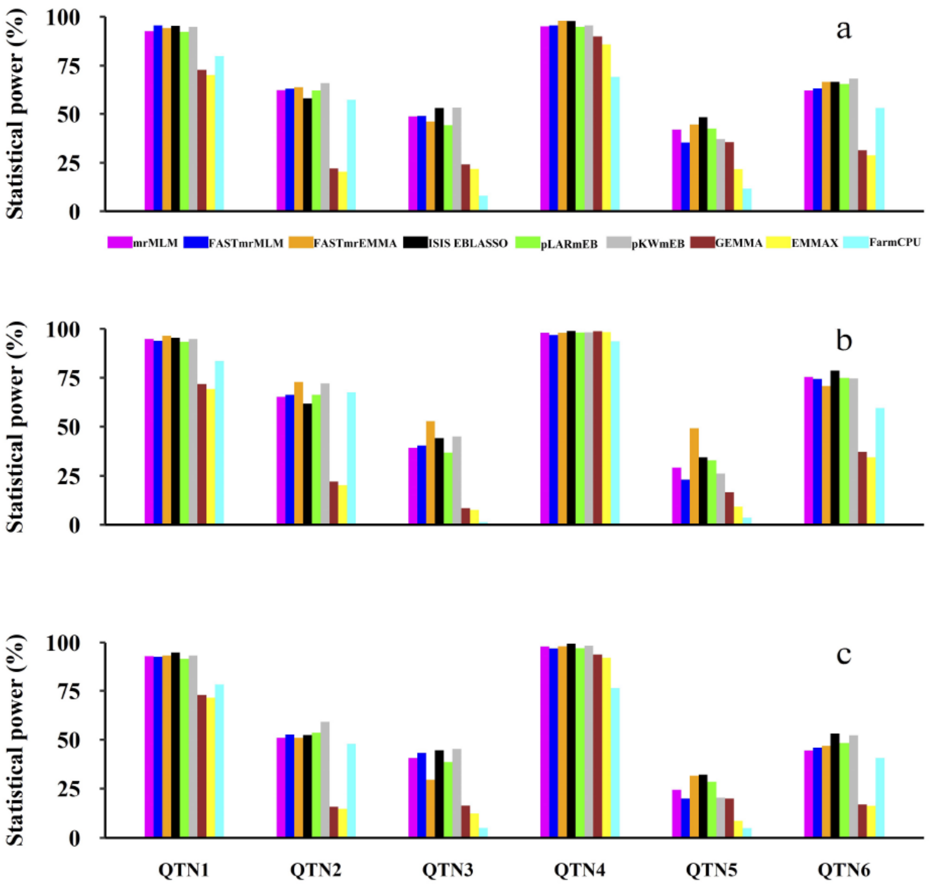


**Figure S3 Comparison of statistical powers in QTN detection between the new and existing methods in three simulation experiments**

**a.** The first simulation experiment. **b.** The second simulation experiment. **c.** The third simulation experiment. The new methods include mrMLM, FASTmrMLM, FASTmrEMMA, ISIS EBLASSO, pLARmEB and pKWmEB, while the existing methods include GEMMA, EMMAX and FarmCPU.

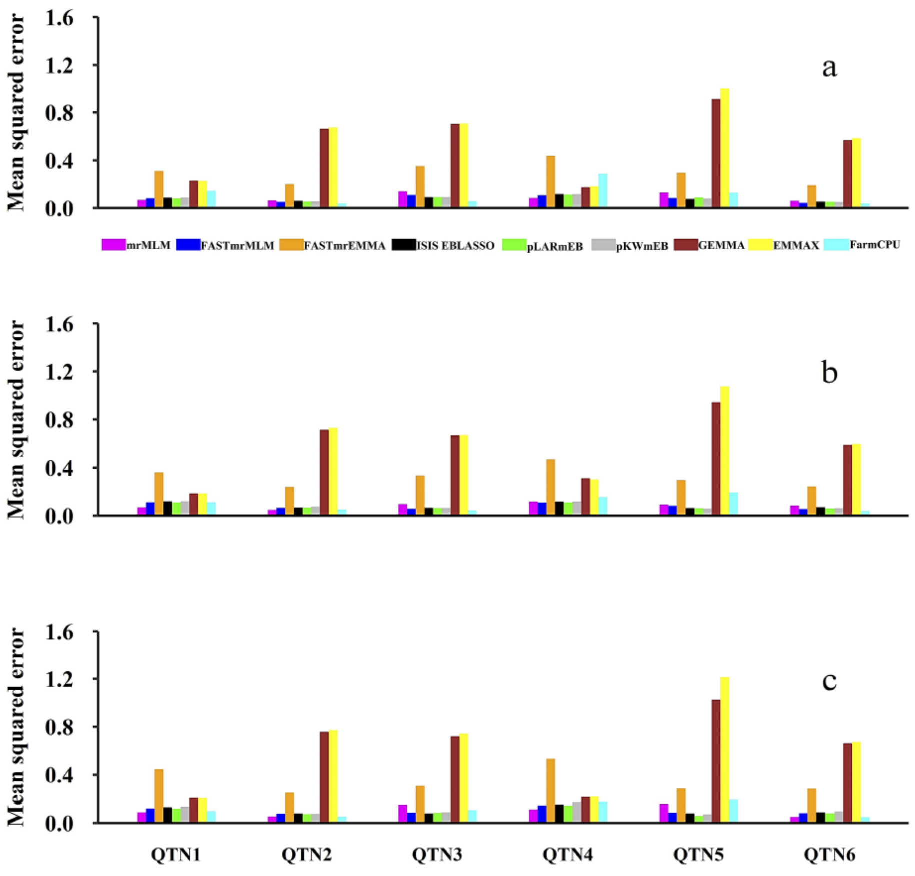


**Figure S4 Comparison of mean squared errors in QTN effect estimation between the new and existing methods in three simulation experiments**

**a.** The first simulation experiment. **b.** The second simulation experiment. **c.** The third simulation experiment. The new and existing methods are the same as those in Figure S3.

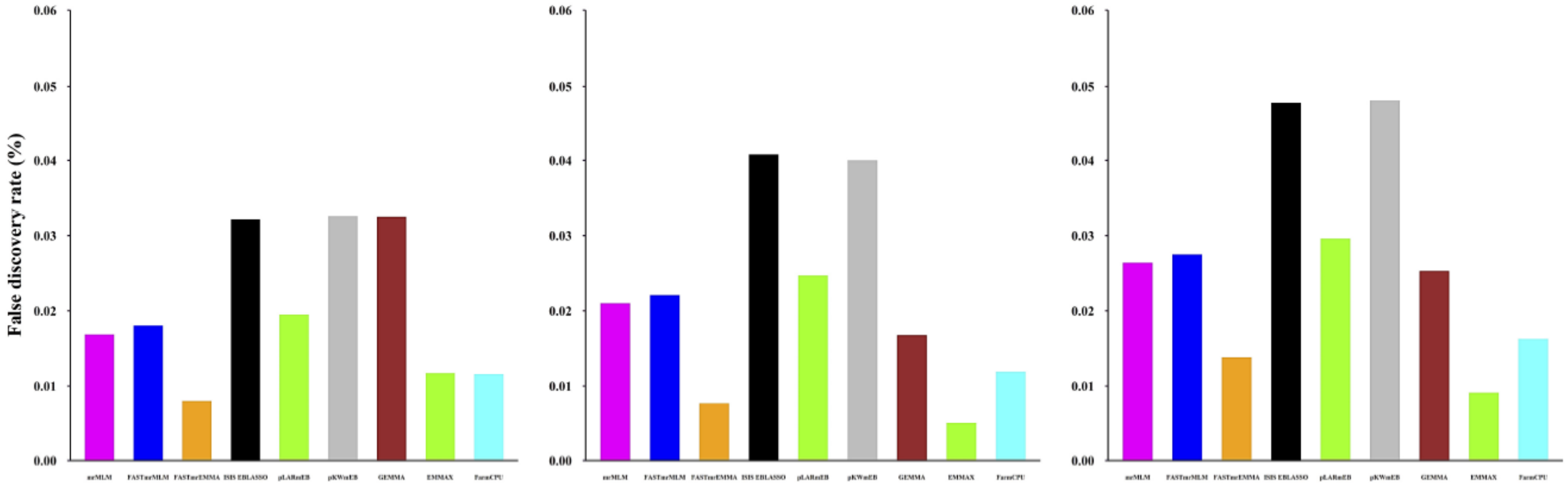


**Figure S5 Comparison of false positive rates (%) in QTN detection between the new and existing methods in three simulation experiments**

**a.** The first simulation experiment. **b.** The second simulation experiment. **c.** The third simulation experiment. The new and existing methods are the same as those in Figure S3.

**Table S9. Comparison of power (%), mean squared error (MSE), and false positive rate (FPR, %) for nine GWAS methods in the first Monte Carlo simulation experiment where six QTNs were simulated**

| Method* | $QTL_1$ | | $QTL_2$ | | $QTL_3$ | | $QTL_4$ | | $QTL_5$ | | $QTL_6$ | | FPR (%) |
|---|---|---|---|---|---|---|---|---|---|---|---|---|---|
| | Power | MSE | Power | MSE | Power | MSE | Power | MSE | Power | MSE | Power | MSE | |
| mrMLM | 92.6 | 0.0703 | 61.9 | 0.0663 | 48.7 | 0.1413 | 95.0 | 0.0867 | 42.0 | 0.1316 | 61.7 | 0.0638 | 0.0167 |
| FASTmrMLM | 95.5 | 0.0839 | 63.1 | 0.0533 | 48.9 | 0.1115 | 95.6 | 0.1092 | 35.4 | 0.0856 | 63.3 | 0.0465 | 0.0179 |
| FASTmrEMMA | 94.2 | 0.3111 | 63.9 | 0.2011 | 46.2 | 0.3516 | 97.9 | 0.4394 | 44.6 | 0.2952 | 66.7 | 0.1920 | 0.0080 |
| ISIS EBLASSO | 95.3 | 0.0897 | 58 | 0.0622 | 53.0 | 0.0930 | 97.8 | 0.1190 | 48.3 | 0.0784 | 66.6 | 0.0556 | 0.0322 |
| pLARmEB | 92.3 | 0.0821 | 61.8 | 0.0561 | 44.2 | 0.0930 | 94.8 | 0.1128 | 42.6 | 0.0927 | 65.6 | 0.0531 | 0.0195 |
| pKWmEB | 94.8 | 0.0898 | 66.1 | 0.0580 | 53.3 | 0.0943 | 95.6 | 0.1177 | 37.1 | 0.0801 | 68.3 | 0.0519 | 0.0326 |
| GEMMA | 72.8 | 0.2309 | 22.1 | 0.6634 | 24.2 | 0.7049 | 89.9 | 0.1766 | 35.6 | 0.9128 | 31.4 | 0.5704 | 0.0325 |
| EMMAX | 70.2 | 0.2284 | 20.4 | 0.6754 | 21.9 | 0.7091 | 85.8 | 0.1834 | 21.7 | 1.0052 | 28.9 | 0.5856 | 0.0117 |
| FarmCPU | 79.9 | 0.1457 | 57.2 | 0.0370 | 8.20 | 0.0590 | 69.2 | 0.2882 | 11.8 | 0.1297 | 53.1 | 0.0341 | 0.0116 |

*: All the results were re-calculated using our mrMLM v3.0, including mrMLM, FASTmrMLM, FASTmrEMMA, ISIS EBLASSO, pLARmEB and pKWmEB, which were published in Sci Rep 2016,6:19444, Doctoral Thesis of Tamba (2017) at Nanjing Agricultural University, Briefings in Bioinformatics 2017, bbw145, doi: 10.1093/bib/bbw145, PLoS Computational Biology 13(1): e1005357, Heredity 2017, 118(6): 517-524, and Heredity 2018, 120: 208-218, respectively. Note that the results of pLARmEB in Zhang et al. (2017) aren't consistent with those in the published paper, because there is one mistake in selecting potentially associated markers in the Monte Carlo simulation experiments of Zhang et al. (2017). The same is true for the later Tables.

**Table S10. Comparison of power (%), mean squared error (MSE) and false positive rate (FPR, %) for nine GWAS methods in the second Monte Carlo simulation experiment where six QTNs along with polygenic background were simulated**

| Method* | $QTL_1$ | | $QTL_2$ | | $QTL_3$ | | $QTL_4$ | | $QTL_5$ | | $QTL_6$ | | FPR (%) |
|---|---|---|---|---|---|---|---|---|---|---|---|---|---|
| | Power | MSE | Power | MSE | Power | MSE | Power | MSE | Power | MSE | Power | MSE | |
| mrMLM | 94.8 | 0.0694 | 65.3 | 0.0481 | 39.4 | 0.0959 | 98.0 | 0.1176 | 29.3 | 0.0929 | 75.4 | 0.0850 | 0.0210 |
| FASTmrMLM | 93.9 | 0.1118 | 66.2 | 0.0664 | 40.6 | 0.0586 | 96.9 | 0.1107 | 23.1 | 0.0837 | 74.4 | 0.0566 | 0.0221 |
| FASTmrEMMA | 96.5 | 0.3620 | 72.8 | 0.2394 | 52.7 | 0.3342 | 98.0 | 0.4690 | 49.1 | 0.2961 | 70.8 | 0.2435 | 0.0077 |
| ISIS EBLASSO | 95.5 | 0.1187 | 61.8 | 0.0665 | 44.0 | 0.0662 | 99.0 | 0.1184 | 34.5 | 0.0641 | 78.6 | 0.0704 | 0.0408 |
| pLARmEB | 93.4 | 0.1098 | 66.2 | 0.0682 | 37.0 | 0.0638 | 98.1 | 0.1095 | 33.0 | 0.0615 | 74.9 | 0.0598 | 0.0247 |
| pKWmEB | 94.8 | 0.1202 | 72.1 | 0.0757 | 44.9 | 0.0634 | 98.2 | 0.1175 | 26.3 | 0.0568 | 74.6 | 0.0614 | 0.0400 |
| GEMMA | 71.7 | 0.1852 | 22.1 | 0.7121 | 8.50 | 0.6665 | 98.8 | 0.3111 | 16.6 | 0.9415 | 37.3 | 0.5915 | 0.0166 |
| EMMAX | 69.2 | 0.1857 | 20.2 | 0.7301 | 7.60 | 0.6704 | 98.4 | 0.3031 | 9.30 | 1.0748 | 34.6 | 0.5989 | 0.0051 |
| FarmCPU | 83.5 | 0.1120 | 67.5 | 0.0521 | 1.30 | 0.0426 | 93.7 | 0.1537 | 3.60 | 0.1950 | 59.5 | 0.0381 | 0.0119 |

**Table S11. Comparison of power (%), mean squared error (MSE) and false positive rate (FPR, %) for nine GWAS methods in the third Monte Carlo simulation experiment where six QTNs along with three epistatic QTNs were simulated**

| Method* | $QTL_1$ | | $QTL_2$ | | $QTL_3$ | | $QTL_4$ | | $QTL_5$ | | $QTL_6$ | | FPR (%) |
|---|---|---|---|---|---|---|---|---|---|---|---|---|---|
| | Power | MSE | Power | MSE | Power | MSE | Power | MSE | Power | MSE | Power | MSE | |
| mrMLM | 93.0 | 0.0885 | 50.9 | 0.0548 | 40.6 | 0.1519 | 97.9 | 0.1118 | 24.4 | 0.1611 | 44.5 | 0.0507 | 0.0264 |
| FASTmrMLM | 92.6 | 0.1216 | 52.7 | 0.0770 | 43.2 | 0.0858 | 97.0 | 0.1443 | 19.7 | 0.0852 | 45.9 | 0.0820 | 0.0275 |
| FASTmrEMMA | 93.3 | 0.4450 | 50.9 | 0.2464 | 29.5 | 0.3030 | 98.0 | 0.5331 | 31.6 | 0.2827 | 46.9 | 0.2799 | 0.0138 |
| ISIS EBLASSO | 94.9 | 0.1323 | 52.4 | 0.0804 | 44.6 | 0.0794 | 99.4 | 0.1553 | 32.0 | 0.0788 | 53.1 | 0.0892 | 0.0477 |
| pLARmEB | 91.6 | 0.1196 | 53.6 | 0.0747 | 38.6 | 0.0835 | 97.1 | 0.1433 | 28.5 | 0.0598 | 48.3 | 0.0795 | 0.0296 |
| pKWmEB | 93.3 | 0.1359 | 59.1 | 0.0757 | 45.3 | 0.0893 | 98.4 | 0.1654 | 20.1 | 0.0722 | 52.3 | 0.0966 | 0.0480 |
| GEMMA | 72.9 | 0.2044 | 15.4 | 0.7576 | 16.0 | 0.7218 | 93.8 | 0.2131 | 19.7 | 1.0254 | 16.7 | 0.6615 | 0.0253 |
| EMMAX | 71.6 | 0.2031 | 14.4 | 0.7716 | 12.6 | 0.7415 | 92.1 | 0.2160 | 8.70 | 1.2131 | 15.9 | 0.6732 | 0.0091 |
| FarmCPU | 78.3 | 0.0981 | 47.8 | 0.0534 | 5.10 | 0.1065 | 76.5 | 0.1709 | 5.00 | 0.1899 | 40.6 | 0.0487 | 0.0161 |

### Supplementary material E. User manual for software mrMLM v4.0

**Disclaimer**: While extensive testing has been performed by Yuan-Ming Zhang's Lab at Crop Information Center of College of Plant Science and Technology, Huazhong Agricultural University, the results are, in general, reliable, correct or appropriate. However, results are not guaranteed for any specific datasets. We strongly recommend that users validate the mrMLM results with other software packages, i.e., GEMMA, EMMAX, GAPIT v2 & PLINK.

**Download website**:
https://cran.r-project.org/web/packages/mrMLM/index.html or
https://bigd.big.ac.cn/biocode/tools/BT007077

**Citation**:

| Method or software | References |
|---|---|
| **mrMLM** | Wang et al. ***Scientific Reports*** 2016, **6**:19444 |
| **ISIS EM-BLASSO** | Tamba et al. ***PLoS Computational Biology*** 2017, 13(1): e1005357. |
| **pLARmEB** | Zhang et al. ***Heredity*** 2017, 118: 517–524 |
| **FASTmrEMMA** | Wen et al. ***Briefings in Bioinformatics*** 2018, 19(4): 700–712. DOI: 10.1093/bib/bbw145 |
| **pKWmEB** | Ren et al. ***Heredity*** 2018, 120(3): 418–428 |
| **FASTmrMLM** | Tamba & Zhang, ***bioRxiv*** preprint first posted online 2018, doi: https://doi.org/10.1101/341784 |
| | Zhang et al. ***Genomics, Proteomics & Bioinformatics***, Resubmission |
| **Software mrMLM** | Zhang et al. ***Genomics, Proteomics & Bioinformatics***, Resubmission |

Note: These references are listed in section of References.

This work was supported by the National Natural Science Foundation of China (31571268, 31871242 and U1602261), Huazhong Agricultural University Scientific & Technological Self-innovation Foundation (Program No. 2014RC020), and State Key Laboratory of Cotton Biology Open Fund (CB2017B01 & CB2019B01).

### INTRODUCTION

#### 1.1 Why mrMLM?

**mrMLM** (**m**ulti-locus **r**andom-SNP-effect **M**ixed **L**inear **M**odel) program is an R package for multi-locus genome-wide association studies (GWAS). At present this program (v4.0) includes six methods: 1) mrMLM, 2) FASTmrMLM (Fast multi-locus random-SNP-effect EMMA), 3) ISIS EM-BLASSO (Iterative Sure Independence Screening EM-Bayesian LASSO), 4) pLARmEB (polygenic-background-control-based least angle regression plus empirical Bayes), 5) pKWmEB (polygenic-background-control-based Kruskal-Wallis test plus empirical Bayes); and 6) fast mrMLM (FASTmrMLM).

In the mrMLM, FASTmrMLM, FASTmrEMMA and pKWmEB methods, the package qqman is used to draw the Manhattan and QQ plots. In the pLARmEB and ISIS EM-BLASSO methods, the package ggplot2 is used to draw the LOD score plot.

mrMLM 4.0 works well on Windows, Linux (desktop) and MacOS.

#### 1.2 Getting started

The software package mrMLM runs only in the R software environment and can be freely downloaded from https://cran.r-project.org/web/packages/mrMLM.GUI/index.html, or requested from the maintainer, Dr Yuan-Ming Zhang at College of Plant Science and Technology, Huazhong Agri Univ.

##### 1.2.1 One-Click installation

Within R environment, the mrMLM software can be installed online using the below command:

```
install.packages("mrMLM")
```

##### 1.2.2 Step-by-step installation

###### 1.2.2.1 Install the add-on packages

**Offline installation** Users may download the below 49 packages from CRAN (https://cran.r-project.org/), github (https://github.com/) and google search.

assertthat, calibrate, cli, codetools, coin, colorspace, crayon, data.table, dichromat, digest, doParallel, foreach, ggplot2, glue, gtable, iterators, labeling, lars, lazyeval, lpsolve, magrittr, MASS, modeltools, multcomp, munsell, mvtnorm, ncvreg, openxlsx, pillar, plyr, qqman, R6, RColorBrewer, Rcpp, reshape2, rlang, sampling, sandwich, scales, sourcetools, stringi, stringr, sbl, TH.data, tibble, utf8, viridisLite, zip, zoo.

Under the R environment, then, users find “Packages”—“Install package(s) from local files…”, select all the above 49 packages, and install them offline.

###### 1.2.2.2 Install mrMLM

Open R GUI, select "Packages"—"Install package(s) from local files…" and then find the mrMLM package which you have downloaded on your desktop.

**User Manual file** Users can decompress the mrMLM package and find the User Manual file (name: **Instruction.pdf**) in the folder of “…/mrMLM/inst/doc”.

##### 1.2.3 Run mrMLM

Once the software mrMLM is installed, users may run it using two commands:

```
library("mrMLM")
mrMLM(***)
```
(***: please see § 2.1.2 Example)

If users re-use the software mrMLM, users also use the above two commands.

### 2. Function

#### 2.1 mrMLM()

##### 2.1.1 Parameter settings

| Parameter | Meaning | File format | Note |
|---|---|---|---|
| **fileGen** | File path & name in your computer, i.e., fileGen="D:/Users/Genotype_num.csv" | *.csv; *.txt (Genotypic values. **Row**: markers; **Column**: individuals) | Tables 1~3 |
| **filePhe** | File path & name in your computer, i.e., filePhe="D:/Users/Phenotype.csv" | *.csv; *.txt (Phenotypic values. **Row**: individual; **Column**: traits) | Table 4 |
| **fileKin** | File path & name in your computer, i.e., fileKin="D:/Users/Kinship.csv" or fileKin=NULL | *.csv; *.txt (Kinship matrix. **Row** & **Column**: individuals) | Table 5 |
| **filePS** | File path & name in your computer,i.e., filePS="D:/Users/PopStr.csv" or filePS=NULL | *.csv; *.txt [Population structure. **Row**: individual; **Column**: sub-populations 1, 2, ---, *k* (No. of sub-populations)] | Table 6~8 |
| **PopStrType** | Three types of population structures: *Q* (*Q* matrix), PCA (Principal components), EvolPopStr (Evolutionary population structure) | | |
| **fileCov** | File path & name in your computer,i.e., fileCov="D:/Users/Covariate.csv" or fileCov=NULL | *.csv; *.txt (Covariate. **Row**: individual; **Column**: covariate 1, 2, ---, *k* (No. of covariate)) | Table 9 |
| **Genformat** | Format for genotypic codes: Num (number), Cha (character) & Hmp (Hapmap), i.e., Genformat="Num" | | |
| **method** | Six multi-locus GWAS methods. Users may select one to six methods. For example, method=c("mrMLM","FASTmrMLM","FASTmrEMMA","pLARmEB","pKWmEB","ISIS EM-BLASSO") | | |
| **Likelihood** | This parameter is only for FASTmrEMMA, including restricted maximum likelihood (REML) and maximum likelihood (ML). Likelihood="REML" or Likelihood="ML" | | |
| **trait** | Traits analyzed from number 1 to number 2. For example, **trait=1:3** indicates that users analyze the first to third traits. | | |
| **SearchRadius** | This parameter is only for mrMLM and FASTmrMLM, indicating Search Radius in search of potentially associated QTN. **SearchRadius=20** indicates that only one potentially associated QTN was selected within 20 kb. | | |
| **CriLOD** | Critical LOD score for significant QTN. **CriLOD=3** indicates that the critical LOD score for significant QTN is set at 3.0. | | |
| **SelectVariable** | This parameter is only for pLARmEB. **SelectVariable=50** indicates that 50 potentially associated variables are selected from each chromosome. Users may change this number in real data analysis in order to obtain the best final results. | | |
| **Bootstrap** | This parameter is only for pLARmEB, including FALSE & TRUE. **Bootstrap=FALSE** indicates the analysis of only real dataset; **Bootstrap=TRUE** indicates the analysis of both real dataset and four resampling datasets. | | |
| **DrawPlot** | This parameter is for all the six methods, including FALSE and TRUE. **DrawPlot=FALSE** indicates no figure output; **DrawPlot=TRUE** indicates the output of the Manhattan, QQ and LOD score against genome position figures. | | |
| **Plotformat** | This parameter is for all the figure files, including *.jpeg, *.png, *.tiff and *.pdf. **Plotformat="jpeg"** indicates the *.jpeg format of plot file. | | |
| **Resolution** | This parameter is for all the figure files, including Low and High. **Resolution="Low"** indicates low figure resolution. | | |
| **dir** | Save path in your computer, i.e, "D:/Users" | | |

#### 2.1.2 Example

#### The full codes

```
mrMLM(fileGen="D:/Users/Genotype_num.csv",filePhe="D:/Users/Phenotype.csv",fileKin=NULL,filePS=NULL
,PopStrType=NULL,fileCov=NULL,Genformat="Num",method=c("mrMLM","FASTmrMLM","FASTmrEMMA
","pLARmEB","pKWmEB","ISIS     EM-BLASSO"),Likelihood="REML",trait=1:3,SearchRadius=20,CriLOD=3,
SelectVariable=50,Bootstrap=FALSE,DrawPlot=FALSE, Plotformat="jpeg",Resolution="Low",dir="D:/Users")
```

#### The reduced codes

mrMLM(fileGen="D:/Users/Genotype_num.csv", filePhe="D:/Users/Phenotype.csv", Genformat="Num", method=c("mrMLM","FASTmrMLM","FASTmrEMMA","pLARmEB","pKWmEB","ISIS EM-BLASSO"), trait=1:3, CriLOD=3, dir="D:/Users")

It should be noted that users must set "fileGen", "filePhe", "Genformat", "method", "trait", "CriLOD" and "dir", and the other eight parameters can be default in function, including PopStrType="Q"; Likelihood="REML"only for FASTmrEMMA; SearchRadius=20 only for mrMLM and FASTmrMLM; SelectVariable=50 & Bootstrap=FALSE only for pLARmEB; DrawPlot=TRUE; Plotformat= "jpeg"; Resolution= "Low".

##### 2.1.3 Dataset format

**Numeric format for dataset "fileGen"** (Table 1) The first column, named "**rs#**", stands for marker ID, i.e., "PZB00859.1". The second column, named "**chrom**", stands for chromosome, i.e., numeric variable "1". The third column, named "**pos**", stands for the position (bp) of SNP on the chromosome. The fourth column, named "**genotype for code 1**", indicates reference base for code variable $x = 1$. Among the remaining columns, each column lists all the genotypes for one individual, and the first row shows the individual names. For each marker, homozygous genotypes are expressed by 1 and -1, respectively, and the heterozygous and missing genotypes are indicated by zero. If the base for the first individual is missing, the base firstly observed in this row is what we list. Note that the genotype with code **1** will be also appeared in the **Result** files.

**Table 1. The numeric format of the genotypic dataset**

| rs# | chrom | pos | genotype for code 1 | 33-16 | Nov-38 | A4226 | A4722 |
|---|---|---|---|---|---|---|---|
| PZB00859.1 | 1 | 157104 | C | 1 | 1 | 1 | 1 |
| PZA01271.1 | 1 | 1947984 | C | 1 | -1 | 1 | -1 |
| PZA03613.2 | 1 | 2914066 | G | 1 | 1 | 1 | 1 |
| PZA03613.1 | 1 | 2914171 | T | 1 | 1 | 1 | 1 |
| PZA03614.2 | 1 | 2915078 | G | 1 | 1 | 1 | 1 |
| PZA03614.1 | 1 | 2915242 | T | 1 | 1 | 1 | 1 |
| PZA02117.1 | 1 | 223466480 | A | 1 | 1 | 1 | -1 |
| PZA00403.5 | 1 | 223466873 | T | 1 | 1 | 1 | 0 |
| ⋮ | ⋮ | ⋮ | ⋮ | ⋮ | ⋮ | ⋮ | ⋮ |

**Character format for dataset "fileGen"** (Table 2) The first three columns are same as those in Table 1. The differences are that the marker values are character, such as **A, T, C, G** and **N**, and the other notations are heterozygous genotypes. The "**N**" indicates the missing of genotypes. The first rows from the fourth to last columns are individual name.

**Table 2. The character format of the genotypic dataset**

| rs# | chrom | pos | 33-16 | Nov-38 | A4226 | A4722 |
|---|---|---|---|---|---|---|
| PZB00859.1 | 1 | 157104 | C | C | C | C |
| PZA01271.1 | 1 | 1947984 | C | G | C | G |
| PZA03613.2 | 1 | 2914066 | G | G | G | G |
| PZA03613.1 | 1 | 2914171 | T | T | T | T |
| ⋮ | ⋮ | ⋮ | ⋮ | ⋮ | ⋮ | ⋮ |

**Hapmap format for dataset "fileGen"** (Table 3) Please see the TASSEL software in details. Here we introduce simply. The first eleven columns describe the specific information of markers and individuals, and these column names must be **"rs#"**, **"alleles"**, **"chrom"**, **"pos"**, **"strand"**, **"assembly#"**, **"center"**, **"protLSID"**, **"assayLSID"**, **"panelLSID"** and **"QCcode"**. In the **"rs#"** (1st)**, "chrom"** (3rd) **and "pos"** (4th) columns, the information has been described as the above. The values of marker genotypes should be character, such as **AA, TT, CC, GG, NN, AC** and **AG**, where the "**NN**" indicates the missing or unknown of genotypes. In the 2nd and 5th to 11th columns, **"NA"** indicates **no information** available. All the individual genotypic information will be showed from the 12th to last columns. In each column, individual name is listed in the first row, i.e., "33-16", and the others are the genotypes (character).

**Table 3. The hapmap format of the genotypic dataset**

| rs# | alleles | chrom | pos | strand | assembly# | center | protLSID | assayLSID | panelLSID | QCcode | 33-16 | ··· |
|---|---|---|---|---|---|---|---|---|---|---|---|---|
| PZB00859.1 | A/C | 1 | 157104 | + | AGPv1 | Panzea | NA | NA | maize282 | NA | CC | ··· |
| PZA01271.1 | C/G | 1 | 1947984 | + | AGPv1 | Panzea | NA | NA | maize282 | NA | CC | ··· |
| PZA03613.2 | G/T | 1 | 2914066 | + | AGPv1 | Panzea | NA | NA | maize282 | NA | GG | ··· |
| PZA03613.1 | A/T | 1 | 2914171 | + | AGPv1 | Panzea | NA | NA | maize282 | NA | TT | ··· |
| PZA03614.2 | A/G | 1 | 2915078 | + | AGPv1 | Panzea | NA | NA | maize282 | NA | GG | ··· |
| PZA03614.1 | A/T | 1 | 2915242 | + | AGPv1 | Panzea | NA | NA | maize282 | NA | TT | ··· |
| PZA02117.1 | A/G | 1 | 223466480 | + | AGPv1 | Panzea | NA | NA | maize282 | NA | AA | ··· |
| ⋮ | ⋮ | ⋮ | ⋮ | ⋮ | ⋮ | ⋮ | ⋮ | ⋮ | ⋮ | ⋮ | ⋮ | ··· |

Before implementing GWAS, the above character genotypes should be transferred into numeric information, here the homozygous genotype of each marker for the first individual is transferred into 1, another homozygous genotype for this marker is transferred into -1, and heterozygous and missing genotypes are transferred into zero.

If the base for the first individual is missing, the base firstly observed in this row is what we list.

**Format for the dataset "filePhe"** (Table 4) The **Phenotypic** file should be a file with ***.csv** or ***.txt** format. The first column lists individual ID, i.e., "B46", and "<Phenotype>" should be showed in the first row. Among the other columns, each column lists all the observations for the trait, its trait name is showed in the first row, i.e., "trait1", and phenotypic values are in the corresponding rows of their individuals.

**Table 4. The format of Phenotypic dataset**

| <Phenotype> | trait1 | trait2 | trait3 |
|---|---|---|---|
| B46 | 42 | 43.02 | 44.32 |
| B52 | 72.5 | 71.88 | 72.8 |
| B57 | 41 | 41.7 | 41.42 |
| B64 | 74.5 | 74.43 | 74.5 |
| ⋮ | ⋮ | ⋮ | ⋮ |

**The format for dataset "fileKin"** (Table 5) The Kinship file should be a file with ***.csv** or ***.txt** format. In the first column in Table 5, "263" is sample size ($n$), and "33-16", "Nov-38" and "A4226" are individual ID. Note that "$n$" is the number of common individuals between the phenotypic and genotypic datasets. All the kinship coefficients are listed as an $n \times n$ matrix.

fileKin=NULL indicates that the Kinship matrix is calculated by the software mrMLM. Here only the above $n$ individuals are used to calculate the Kinship matrix. fileKin="D:/Users/Kinship.csv" means that the K matrix with name Kinship.csv is uploaded from the folder "D:/Users". If the number and order of individuals in Kinship.csv are not consistent with those of the above $n$ individuals, our software may match the K matrix in order that the number and order of the transferred K matrix are consistent with those in the above $n$ individuals.

**Table 5. The format of the Kinship dataset**

| 263 | | | | | |
|---|---|---|---|---|---|
| 33-16 | 1.00809 | 0.45954 | 0.50677 | 0.42503 | 0.45591 |
| Nov-38 | 0.45954 | 1.03352 | 0.43048 | 0.47044 | 0.39597 |
| A4226 | 0.50677 | 0.43048 | 1.01717 | 0.45409 | 0.43775 |
| A4722 | 0.42503 | 0.47044 | 0.45409 | 0.89002 | 0.34874 |

| | | | | | |
|---|---|---|---|---|---|
| A188 | 0.45591 | 0.39597 | 0.43775 | 0.34874 | 1.0099 |
| A214N | 0.34693 | 0.33421 | 0.39779 | 0.29244 | 0.33058 |
| A239 | 0.43593 | 0.46499 | 0.40323 | 0.36691 | 0.39597 |
| A272 | 0.34874 | 0.40505 | 0.31423 | 0.3887 | 0.44138 |
| A441-5 | 0.47952 | 0.44138 | 0.47226 | 0.47952 | 0.49224 |
| A554 | 0.39779 | 0.45954 | 0.5431 | 0.48679 | 0.4214 |
| ⋮ | ⋮ | ⋮ | ⋮ | ⋮ | ⋮ |

***Q* matrix format for dataset "filePS"** (Table 6) The *Q* matrix dataset in **Table 6** consists of a ($n$+2) × ($k$+1) matrix, where $n$ is the number of the above common individuals and $k$ is the number of sub-populations. In the first column, "<**PopStr**>" and "<**ID**>" should present in the first and second rows, respectively; "33-16", "Nov-38" and "A4226" are individual ID. In the 2nd to ($k$+1)-th columns, "$Q_1$" to "$Q_k$" indicate sub-populations. In the third row, "0.014", "0.972" and "0.014" are the posterior probabilities of the "33-16" individual in the 1st, 2nd and 3rd subpopulations, respectively. When the *Q* matrix is uploaded to the software, the software will automatically delete the column whose sum is the smallest.

**Table 6. The format of the filePS dataset**

| <PopStr> | | | |
|---|---|---|---|
| <ID> | Q1 | Q2 | Q3 |
| 33-16 | 0.014 | 0.972 | 0.014 |
| Nov-38 | 0.003 | 0.993 | 0.004 |
| A4226 | 0.071 | 0.917 | 0.012 |
| A4722 | 0.035 | 0.854 | 0.111 |
| A188 | 0.013 | 0.982 | 0.005 |
| A214N | 0.762 | 0.017 | 0.221 |
| A239 | 0.035 | 0.963 | 0.002 |
| A272 | 0.019 | 0.122 | 0.859 |
| ⋮ | ⋮ | ⋮ | ⋮ |

**Principal components format for dataset "filePS"** (Table 7) The principal component dataset in Table 7 consists of a ($n$+2) × ($k$+1) matrix, where $n$ is the number of the common individuals and $k$ is the number of principal components. In the first column, "<**PCA**>" and "<**ID**>" should present in the first and second rows, respectively; "33-16", "Nov-38" and "A4226" are individual ID. In the 2nd to ($k$+1)-th columns, "$PC_1$" to "$PC_k$" indicate the first to $k$-th principal components. In the second column,

"0.306", ⋯, "0.216" are the scores of the first principal component for the 1st to 9-th individuals, respectively.

**Table 7. The dataset format of principal components**

| <PCA> | | | |
|---|---|---|---|
| <ID> | PC1 | PC2 | PC3 |
| 33-16 | 0.306 | 0.029 | 0.226 |
| Nov-38 | -0.708 | -2.071 | 1.413 |
| A4226 | -2.330 | 0.116 | -0.824 |
| A4722 | 1.059 | 0.470 | -1.315 |
| A188 | -2.376 | 1.087 | -0.135 |
| A214N | -2.346 | 0.516 | 0.666 |
| A239 | -0.099 | -0.318 | -0.473 |
| A272 | -0.053 | 0.093 | -0.275 |
| A441-5 | 0.216 | -0.535 | -0.159 |
| ⋮ | ⋮ | ⋮ | ⋮ |

**Evolutionary population structure format for dataset "filePS"** (Table 8) The evolutionary population structure dataset in Table 8 consists of a ($n$+2) × 2 matrix, where $n$ is the number of the common individuals. In the first column, "<**EvolPopStr**>" and "<**ID**>" should present in the first and second rows, respectively; "33-16", "Nov-38" and "A4226" are individual ID. In the second column, "EvolType" indicates the evolutionary type, i.e., the evolutionary types for individuals "33-16" and "A4722" are "A" and "B", respectively.

**Table 8. The dataset format of evolutionary population structure**

| <EvolPopStr> | |
|---|---|
| <ID> | EvolType |
| 33-16 | A |
| A4722 | B |
| A188 | A |
| A239 | B |
| ⋮ | ⋮ |

filePS=NULL indicates no inclusion of population structure in the genetic model. filePS="D:/Users/PopStr.csv" means that population structure dataset with name PopStr.csv is uploaded from the folder "D:/Users". If the number and order of individuals in PopStr.csv aren't consistent with those of the above common individuals, our software may match the population structure matrix in order that the number and order of new matrix are consistent with those in the above common individuals.

**The format for dataset "fileCov"** (Table 9) The "**Covariate**" dataset consists of the ($n$+2) × ($k$+1) matrix, where $n$ is the number of the common individuals and $k$ is the number of covariates. In the first column, "<**Covariate**>" and "<**ID**>" should present in the first and second rows, respectively. If covariate is categorical, it should be named as Cate_covariate*. If covariate is continuous, it should be named as Con_covariate* (Table 9).

fileCov=NULL indicates no inclusion of covariates in the genetic model. fileCov="D:/Users/covariate.csv" means that the covariates with name covariate.csv are uploaded from the folder "D:/Users". If the number and order of individuals in the uploaded file are not consistent with those in the above common individuals, our software need to change the number and order of individuals in order to match the above datasets.

**Table 9. The format of the fileCov dataset**

| <Covariate> | | | | |
|---|---|---|---|---|
| <ID> | Cate_covariate1 | Cate_covariate2 | Con_covariate1 | Con_covariate2 |
| 33-16 | A | C | 349.5 | 374 |
| Nov-38 | B | C | 205 | 452 |
| A4226 | A | D | 300 | 374 |
| A4722 | A | D | 190 | 452 |
| A188 | B | C | 213 | 374 |
| ⋮ | ⋮ | ⋮ | ⋮ | ⋮ |

##### 2.1.4 Result

At the work directory of your R, two Result files for the $i$th trait, "$i$_intermediate result.csv" and "$i$_Final result.csv", will appear.

In the **intermediate result** from the mrMLM method, the results include: Trait ID, Trait name, method, reference sequence number (rs#, marker name), chromosome, marker's position (bp) in the chromosome, SNP effect ( $\gamma_k$ , Effect), -log$_{10}$(P), and genotype for code 1.

In the **Final result** from the mrMLM method, the results include: Trait ID, Trait name, method, reference sequence number (rs#, marker names), chromosome, marker's position (bp) in the chromosome, QTN effect, LOD score, -$\log_{10}$(P), the proportion of phenotypic variance explained by significant QTN ($r^2$), minor allelic frequency, genotype for code 1, residual error variance, and total phenotypic variance.

#### 3. References


1. Zhang Yuan-Ming, Mao Yongcai, Xie Chongqing, Howie Smith , Luo Lang, Xu Shizhong*. Mapping quantitative trait loci using naturally occurring genetic variance among commercial inbred lines of maize (*Zea mays* L.). ***Genetics*** 2005, **169**: 2267–2275. DOI: 10.1534/genetics.104.033217
2. Wang Shi-Bo, Feng Jian-Ying, Ren Wen-Long, Huang Bo, Zhou Ling, Wen Yang-Jun, Zhang Jin, Jim M. Dunwell, Xu Shizhong*, Zhang Yuan-Ming*. Improving power and accuracy of genome-wide association studies via a multi-locus mixed linear model methodology. ***Scientific Reports*** 2016, **6**: 19444. DOI: 10.1038/srep19444
3. Tamba Cox Lwaka, Ni Yuan-Li, Zhang Yuan-Ming*. Iterative sure independence screening EM-Bayesian LASSO algorithm for multi-locus genome-wide association studies. ***PLoS Computational Biology*** 2017, **13**(1): e1005357, DOI: 10.1371/journal.pcbi.1005357
4. Zhang Jin[#], Feng Jian-Ying[#], Ni Yuan-Li, Wen Yang-Jun, Niu Yuan, Tamba Cox Lwaka, Yue Chao, Song Qi-Jian, Zhang Yuan-Ming*. pLARmEB: Integration of least angle regression with empirical Bayes for multi-locus genome-wide association studies. ***Heredity*** 2017, **118**: 517–524. DOI:10.1038/hdy.2017.8
5. Ren Wen-Long[#], Wen Yang-Jun[#], Jim M. Dunwell, Zhang Yuan-Ming*. pKWmEB: Integration of Kruskal-Wallis test with empirical Bayes under polygenic background control for multi-locus genome-wide association study. ***Heredity*** 2018, **120**: 208–218. https://doi.org/10.1038/s41437-017-0007-4
6. Wen Yang-Jun, Zhang Hanwen, Ni Yuan-Li, Huang Bo, Zhang Jin, Feng Jian-Ying, Wang Shi-Bo, Jim M. Dunwell, Zhang Yuan-Ming*, Wu Rongling*. Methodological implementation of mixed linear models in multi-locus genome-wide association studies. ***Briefings in Bioinformatics*** 2018, **19**(4): 700–712. https://doi.org/10.1093/bib/bbw145
7. Tamba Cox Lwaka, Zhang Yuan-Ming*. A fast mrMLM algorithm for multi-locus genome-wide association studies. *bioRxiv* 2018, doi: https://doi.org/10.1101/341784, online (June 7, 2018)
8. Zhang Ya-Wen, Tamba Cox Lwaka, Wen Yang-Jun, Li Pei, Ren Wen-Long, Ni Yuan-Li, Gao Jun, Zhang Yuan-Ming*. mrMLM v4.0: An R platform for multi-locus genome-wide association studies. ***Genomics, Proteomics & Bioinformatics***, resubmission

#### Supplementary material F. User manual for software mrMLM.GUI v4.0

**Disclaimer**: While extensive testing has been performed by Yuan-Ming Zhang's Lab at the Crop Information Center of College of Plant Science and Technology, Huazhong Agricultural University, the results are, in general, reliable, correct or appropriate. However, results are not guaranteed for any specific datasets. We strongly recommend that users validate the mrMLM.GUI results with other software packages, i.e., GEMMA, EMMAX, GAPIT v2 & PLINK.

**Download website**:
https://cran.r-project.org/web/packages/mrMLM.GUI/index.html or
https://bigd.big.ac.cn/biocode/tools/BT007077

| Method or software | References |
|---|---|
| **mrMLM** | Wang et al. ***Scientific Reports*** 2016, **6**:19444 |
| **ISIS EM-BLASSO** | Tamba et al. ***PLoS Computational Biology*** 2017, 13(1): e1005357. |
| **pLARmEB** | Zhang et al. ***Heredity*** 2017, 118: 517–524 |
| **FASTmrEMMA** | Wen et al. ***Briefings in Bioinformatics*** 2018, 19(4): 700–712. DOI: 10.1093/bib/bbw145 |
| **pKWmEB** | Ren et al. ***Heredity*** 2018, 120(3): 418–428 |
| **FASTmrMLM** | Tamba & Zhang, ***bioRxiv*** preprint first posted online 2018, doi: https://doi.org/10.1101/341784 |
| | Zhang et al. ***Genomics, Proteomics & Bioinformatics***, Resubmission |
| **Software mrMLM** | Zhang et al. ***Genomics, Proteomics & Bioinformatics***, Resubmission |

Note: These references are listed in section of References.

**m**ulti-locus
**r**andom-SNP-effect
**M**ixed
**L**inear
**M**odel

This work was supported by the National Natural Science Foundation of China (31571268, 31871242 and U1602261), Huazhong Agricultural University Scientific & Technological Self-innovation Foundation (Program No. 2014RC020), and State Key Laboratory of Cotton Biology Open Fund (CB2017B01 & CB2019B01).

### INTRODUCTION

#### 1.1 Why mrMLM.GUI?

**mrMLM.GUI** (**m**ulti-locus **r**andom-SNP-effect **M**ixed **L**inear **M**odel with **G**raphical **U**ser **I**nterface) program is an R package for multi-locus genome-wide association study (GWAS). At present this program (v4.0) includes six methods: 1) mrMLM, 2) FASTmrMLM (Fast multi-locus random-SNP-effect EMMA), 3) ISIS EM-BLASSO (Iterative Sure Independence Screening EM-Bayesian LASSO), 4) pLARmEB (polygenic-background-control-based least angle regression plus empirical Bayes), 5) pKWmEB (polygenic-background-control-based Kruskal-Wallis test plus empirical Bayes); and 6) FASTmrMLM (fast mrMLM).

In the mrMLM.GUI, the package qqman is used to draw the Manhattan and QQ plots in the FASTmrMLM, FASTmrEMMA and pKWmEB methods, and the package ggplot2 is used to draw the the LOD score plot in the pLARmEB and ISIS EM-BLASSO methods.

The software package mrMLM.GUI 4.0 works well on Windows, Linux (desktop) and MacOS.

#### 1.2 Getting started

The software package mrMLM runs only in the R software environment and can be freely downloaded from https://cran.r-project.org/web/packages/mrMLM.GUI/index.html, or requested from the maintainer, Dr Yuan-Ming Zhang at College of Plant Science and Technology, Huazhong Agri Univ.

##### 1.2.1 One-Click installation

Within R environment, the mrMLM.GUI software can be installed online using the below command:

```
install.packages("mrMLM.GUI")
```

##### 1.2.2 Step-by-step installation

###### 1.2.2.1 Install the add-on packages

**Offline installation** Users may download the below 62 packages from CRAN (https://cran.r-project.org/), github (https://github.com/) and google search.

assertthat, bigmemory, bigmemory.sri, calibrate, cli, codetools, coin, colorspace, crayon, data.table, dichromat, digest, doParallel, foreach, ggplot2, glue, gtable, htmltools, httpuv, iterators, jsonlite, labeling, lars, later, lazyeval, lpsolve, magrittr, MASS, mime, miniUI, modeltools, mrMLM, multcomp, munsell, mvtnorm, ncvreg, openxlsx, pillar, plyr, promises, qqman, R6, RColorBrewer, Rcpp, reshape2, rlang, sampling, sandwich, scales, shiny, shinyjs, sourcetools, stringi, stringr, sbl, TH.data, tibble, utf8, viridisLite, xtable, zip, zoo.

Under the R environment, then, users find “Packages”—“Install package(s) from local files…”, select all the above 62 packages, and install them offline.

###### 1.2.2.2 Install mrMLM.GUI

Open R GUI, select "Packages"—"Install package(s) from local files…" and then find the mrMLM.GUI package in which you have downloaded on your desktop.

**User Manual** Users can decompress the mrMLM.GUI package and find the User Manual file (name: **Instruction.pdf**) in the folder of “…/mrMLM.GUI/inst/doc”.

##### 1.2.3 Run mrMLM.GUI

Once the software mrMLM.GUI is installed, users may run the software using two commands:

```
library("mrMLM.GUI")
mrMLM.GUI()
```

If users re-use the software mrMLM.GUI, users also use the above two commands.

#### 3. Dataset input

##### 2.1 Genotypic dataset

The **Genotypic** file should be a ***.csv** or ***.txt** format file.

**Numeric format for Genotypic dataset** (Table 1) The first column, named "**rs#**", stands for marker ID, i.e., “PZB00859.1”. The second column, named "**chrom**", stands for chromosome, i.e., numeric variable “1”. The third column, named "**pos**", stands for the position (bp) of SNP on the chromosome. The fourth column, named "**genotype for code 1**", indicates reference base for code variable *x* = 1. Among the remaining columns, each column lists all the genotypes for one individual, and the first row shows the individual names. For each marker, homozygous genotypes are expressed by 1 and -1, respectively, and the heterozygous and missing genotypes are indicated by zero. If the base for the first individual is missing, the base firstly observed in this row is what we list. Note that the genotype with code **1** will be also listed in the **Result** files.

**Table 1. The numeric format of the genotypic dataset**

| rs# | chrom | pos | genotype for code 1 | 33-16 | Nov-38 | A4226 | A4722 |
|---|---|---|---|---|---|---|---|
| PZB00859.1 | 1 | 157104 | C | 1 | 1 | 1 | 1 |
| PZA01271.1 | 1 | 1947984 | C | 1 | -1 | 1 | -1 |
| PZA03613.2 | 1 | 2914066 | G | 1 | 1 | 1 | 1 |
| PZA03613.1 | 1 | 2914171 | T | 1 | 1 | 1 | 1 |
| PZA03614.2 | 1 | 2915078 | G | 1 | 1 | 1 | 1 |
| PZA03614.1 | 1 | 2915242 | T | 1 | 1 | 1 | 1 |
| PZA02117.1 | 1 | 223466480 | A | 1 | 1 | 1 | -1 |
| PZA00403.5 | 1 | 223466873 | T | 1 | 1 | 1 | 0 |
| ⋮ | ⋮ | ⋮ | ⋮ | ⋮ | ⋮ | ⋮ | ⋮ |

**Character format for Genotypic dataset** The first three columns in Table 2 are same as those in Table 1. The differences are that the marker values are characters, such as **A, T, C, G** and **N**, and the other notations are heterozygous genotypes. The "**N**" indicates the missing of genotypes. The first row from the fourth to last columns lists the names of individuals, i.e., "33-16" and "Nov-38".

**Table 2. The character format of the genotypic dataset**

| rs# | chrom | pos | 33-16 | Nov-38 | A4226 | A4722 |
|---|---|---|---|---|---|---|
| PZB00859.1 | 1 | 157104 | C | C | C | C |
| PZA01271.1 | 1 | 1947984 | C | G | C | G |
| PZA03613.2 | 1 | 2914066 | G | G | G | G |
| PZA03613.1 | 1 | 2914171 | T | T | T | T |
| ⋮ | ⋮ | ⋮ | ⋮ | ⋮ | ⋮ | ⋮ |

**Hapmap format for Genotypic dataset** Please see the TASSEL software in details. Here we describe simply. The first eleven columns describe the specific information of markers and individuals, and their column names must be **"rs#"**, **"alleles"**, **"chrom"**, **"pos"**, **"strand"**, **"assembly#"**, **"center"**, **"protLSID"**,

**"assayLSID"**, **"panelLSID"** and **"QCcode"**. In the **"rs#"** (1st)**, "chrom"** (3rd) **and "pos"** (4th) columns, their information is described as the above in **Table 3**. The values of marker genotypes should be character, such as **AA, TT, CC, GG, NN, AC** and **AG**, where the "**NN**" indicates the missing or unknown of genotypes. In the 2nd and 5th to 11th columns, **"NA"** indicates **no information** available. All the individual genotypic information will be showed from the 12th to last columns. In each column, individual name is listed in the first row, i.e., “33-16”, and the others are the genotypes (character).

**Table 3. The hapmap format of the genotypic dataset**

| rs# | alleles | chrom | pos | strand | assembly# | center | protLSID | assayLSID | panelLSID | QCcode | 33-16 | ··· |
|---|---|---|---|---|---|---|---|---|---|---|---|---|
| PZB00859.1 | A/C | 1 | 157104 | + | AGPv1 | Panzea | NA | NA | maize282 | NA | CC | ··· |
| PZA01271.1 | C/G | 1 | 1947984 | + | AGPv1 | Panzea | NA | NA | maize282 | NA | CC | ··· |
| PZA03613.2 | G/T | 1 | 2914066 | + | AGPv1 | Panzea | NA | NA | maize282 | NA | GG | ··· |
| PZA03613.1 | A/T | 1 | 2914171 | + | AGPv1 | Panzea | NA | NA | maize282 | NA | TT | ··· |
| PZA03614.2 | A/G | 1 | 2915078 | + | AGPv1 | Panzea | NA | NA | maize282 | NA | GG | ··· |
| PZA03614.1 | A/T | 1 | 2915242 | + | AGPv1 | Panzea | NA | NA | maize282 | NA | TT | ··· |
| PZA02117.1 | A/G | 1 | 223466480 | + | AGPv1 | Panzea | NA | NA | maize282 | NA | AA | ··· |
| PZA00403.5 | C/T | 1 | 223466873 | + | AGPv1 | Panzea | NA | NA | maize282 | NA | TT | ··· |
| ⋮ | ⋮ | ⋮ | ⋮ | ⋮ | ⋮ | ⋮ | ⋮ | ⋮ | ⋮ | ⋮ | ⋮ | ··· |

Before implementing GWAS, the above character genotypes should be transferred into numeric information. Here the homozygous genotype of each marker for the first individual is transferred into 1, another homozygous genotype for this marker is transferred into -1, and the heterozygous and missing genotypes are transferred into zero. If the base for the first individual is missing, the base firstly observed in this row is what we list.

#### 2.2 Phenotypic dataset

The **Phenotypic** file with the ***.csv** or ***.txt** format is showed in Table 4. The first column lists individual ID, i.e., “B46”, and “<Phenotype>” should be showed in the first row. Among the other columns, each column lists all the observations for one trait, and its trait name is showed in the first row, i.e., “trait1”.

**Table 4. The format of Phenotypic dataset**

| <Phenotype> | trait1 | trait2 | trait3 |
|---|---|---|---|

| | | | |
|---|---|---|---|
| B46 | 42 | 43.02 | 44.32 |
| B52 | 72.5 | 71.88 | 72.8 |
| B57 | 41 | 41.7 | 41.42 |
| B64 | 74.5 | 74.43 | 74.5 |
| ⋮ | ⋮ | ⋮ | ⋮ |

#### 2.3 Kinship dataset

The Kinship file with the ***.csv** or ***.txt** format is showed in Table 5. In the first column, “263” is sample size (*n*), and “33-16”, “Nov-38” and “A4226” are individual ID. Note that “*n*” is the number of common individuals between the phenotypic and genotypic datasets. All the kinship coefficients are listed as an $n \times n$ matrix.

**Table 5. The format of the Kinship dataset**

| | | | | | |
|---|---|---|---|---|---|
| 263 | | | | | |
| 33-16 | 1.00809 | 0.45954 | 0.50677 | 0.42503 | 0.45591 |
| Nov-38 | 0.45954 | 1.03352 | 0.43048 | 0.47044 | 0.39597 |
| A4226 | 0.50677 | 0.43048 | 1.01717 | 0.45409 | 0.43775 |
| A4722 | 0.42503 | 0.47044 | 0.45409 | 0.89002 | 0.34874 |
| A188 | 0.45591 | 0.39597 | 0.43775 | 0.34874 | 1.0099 |
| A214N | 0.34693 | 0.33421 | 0.39779 | 0.29244 | 0.33058 |
| A239 | 0.43593 | 0.46499 | 0.40323 | 0.36691 | 0.39597 |
| A272 | 0.34874 | 0.40505 | 0.31423 | 0.3887 | 0.44138 |
| A441-5 | 0.47952 | 0.44138 | 0.47226 | 0.47952 | 0.49224 |
| A554 | 0.39779 | 0.45954 | 0.5431 | 0.48679 | 0.4214 |
| ⋮ | ⋮ | ⋮ | ⋮ | ⋮ | ⋮ |

When users select “**Calculate kinship (K) matrix by this software**”, these coefficients between pairs of the above common individuals in the phenotypic and genotypic datasets can be calculated. When users select to input and upload “**Kinship (K)**” matrix file, the number and order of individuals in the uploaded file may be not consistent with those in the phenotypic and genotypic datasets. At this case, our software can let the number and order of individuals in the uploaded K matrix file be consistent with those in the phenotypic and genotypic datasets.

#### 2.4 Population Structure dataset

**Dataset format of $Q$ matrix** The $Q$ matrix dataset in Table 6 consists of a ($n$+2) × ($k$+1) matrix, where $n$ is the number of the common individuals and $k$ is the number of sub-populations. In the first column, "<**PopStr**>" and "<**ID**>" should present in the first and second rows, respectively; "33-16", "Nov-38" and "A4226" are individual ID. In the 2nd to ($k$+1)-th columns, "$Q_1$" to "$Q_k$" indicate sub-populations. In the third row, "0.014", "0.972" and "0.014" are the posterior probabilities of the "33-16" individual from the first, second and third subpopulations, respectively. When the $Q$ matrix is uploaded to the software, the software will automatically delete the column whose sum is the smallest.

**Table 6. The format of the Population Structure dataset**

| <PopStr> | | | |
|---|---|---|---|
| <ID> | Q1 | Q2 | Q3 |
| 33-16 | 0.014 | 0.972 | 0.014 |
| Nov-38 | 0.003 | 0.993 | 0.004 |
| A4226 | 0.071 | 0.917 | 0.012 |
| A4722 | 0.035 | 0.854 | 0.111 |
| A188 | 0.013 | 0.982 | 0.005 |
| A214N | 0.762 | 0.017 | 0.221 |
| A239 | 0.035 | 0.963 | 0.002 |
| A272 | 0.019 | 0.122 | 0.859 |
| A441-5 | 0.005 | 0.531 | 0.464 |
| ⋮ | ⋮ | ⋮ | ⋮ |

**Dataset format of principal components** The principal component dataset in Table 7 consists of a ($n$+2) × ($k$+1) matrix, where $n$ is the number of the common individuals and $k$ is the number of principal components. In the first column, "<**PCA**>" and "<**ID**>" should present in the first and second rows, respectively; "33-16", "Nov-38" and "A4226" are individual ID. In the 2nd to ($k$+1)-th columns, "$PC_1$" to "$PC_k$" indicate the first to $k$-th principal components. In the second column, "0.306", ⋯, "0.216" are the scores of the first principal component for the 1st to 9-th individuals, respectively.

**Table 7. The format of the Principal components dataset**

| <PCA> | | | |
|---|---|---|---|
| <ID> | PC1 | PC2 | PC3 |
| 33-16 | 0.306 | 0.029 | 0.226 |

| | | | |
|---|---|---|---|
| Nov-38 | -0.708 | -2.071 | 1.413 |
| A4226 | -2.330 | 0.116 | -0.824 |
| A4722 | 1.059 | 0.470 | -1.315 |
| A188 | -2.376 | 1.087 | -0.135 |
| A214N | -2.346 | 0.516 | 0.666 |
| A239 | -0.099 | -0.318 | -0.473 |
| A272 | -0.053 | 0.093 | -0.275 |
| A441-5 | 0.216 | -0.535 | -0.159 |
| ⋮ | ⋮ | ⋮ | ⋮ |

**Table 8. The format of the Evolutionary population structure dataset**

| <EvolPopStr> | |
|---|---|
| <ID> | EvolType |
| 33-16 | A |
| Nov-38 | A |
| A4226 | A |
| A4722 | B |
| A188 | A |
| A214N | A |
| A239 | B |
| ⋮ | ⋮ |

**Dataset format of evolutionary population structure** The evolutionary population structure dataset in Table 8 consists of a ($n$+2) × 2 matrix, where $n$ is the number of the common individuals. In the first column, "<**EvolPopStr**>" and "<**ID**>" should present in the first and second rows, respectively; "33-16", "Nov-38" and "A4226" are individual ID. In the second column, "EvolType" indicates the evolutionary type, i.e., the evolutionary types for individuals "33-16" and "A4722" are "A" and "B", respectively.

"**Not included in the model**" indicates no inclusion of population structure in the genetic model. On the contrary, it should be "**Included**". At this case, users should upload the population structure file. If the number and order of individuals in the uploaded file aren't consistent with those in the phenotypic and genotypic datasets, our software may change the population structure matrix in order that the number and order of individuals are consistent with those in the above common individuals.

#### 2.5 Covariate dataset

The "**Covariate**" dataset in Table 9 consists of the ($n$+2) × ($k$+1) matrix, where $n$ is the number of the common individuals and $k$ is the number of covariates. In the first column, "<**Covariate**>" and "<**ID**>" should present in the first and second rows, respectively. The 2nd to ($k$+1)-th columns are covariates. If covariate is categorical, it should be named as Cate_covariate*. If covariate is continuous, it should be named as Con_covariate*.

**Table 9. The format of the fileCov dataset**

| <Covariate> | | | | |
|---|---|---|---|---|
| <ID> | Cate_covariate1 | Cate_covariate2 | Con_covariate1 | Con_covariate2 |
| 33-16 | A | C | 349.5 | 374 |
| Nov-38 | B | C | 205 | 452 |
| A4226 | A | D | 300 | 374 |
| A4722 | A | D | 190 | 452 |
| A188 | B | C | 213 | 374 |
| ⋮ | ⋮ | ⋮ | ⋮ | ⋮ |

"**Not included in the model**" indicates no inclusion of covariates in the genetic model. On the contrary, it should be "**Included**". At this case, users should upload the covariate file. If the number and order of individuals in the uploaded file aren't consistent with those in the above common individuals, our software may change the number and order of individual in order to match the original datasets.

### 4. Operation process

#### 3.1 The Graphical User Interface of mrMLM.GUI

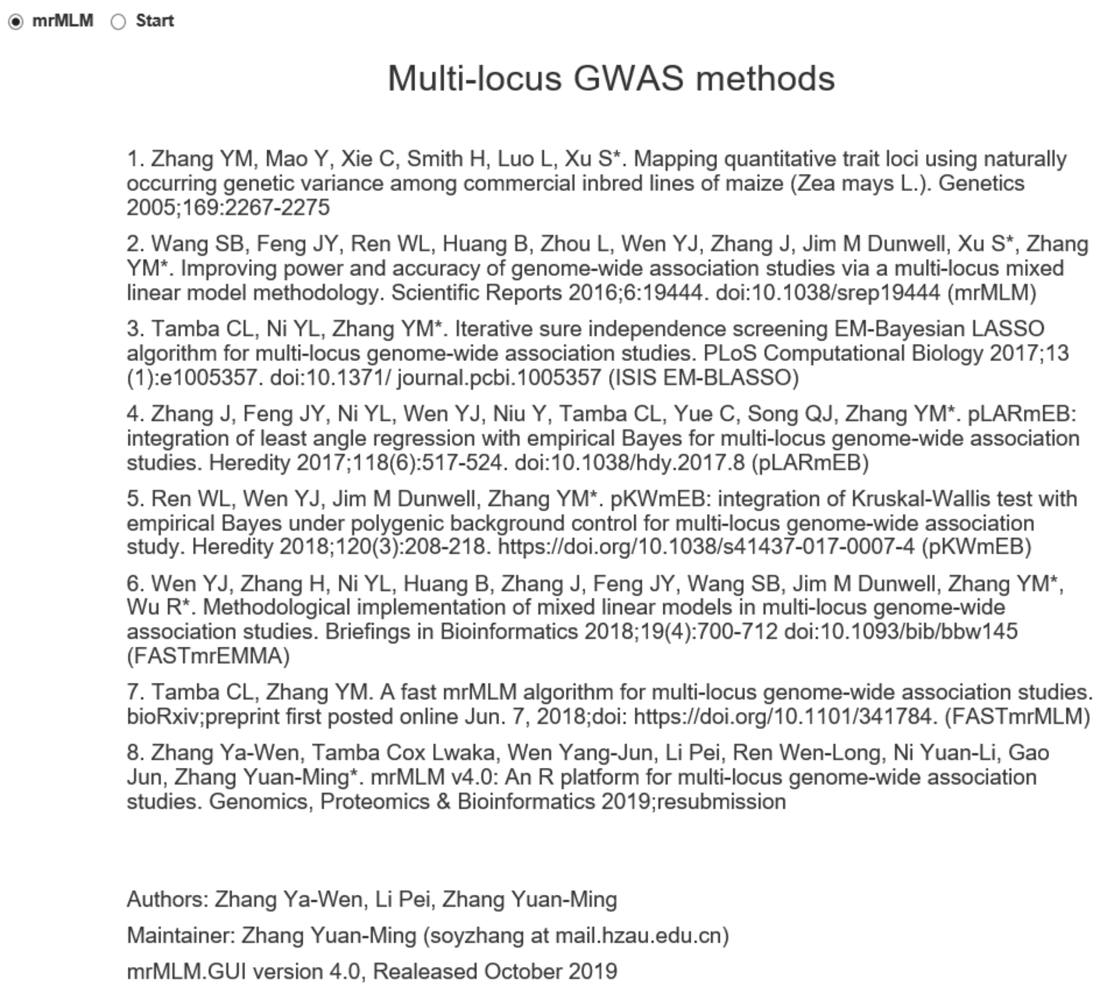


**Figure 1. The Graphical User Interface of mrMLM.GUI**

#### 3.2 Input dataset

Users must upload the genotypic and phenotypic files (Figs 2 & 3), while the Kinship, Population-Structure and Covariate files are optional. In Kinship module, users should upload the Kinship matrix if users select "**Input Kinship (K) matrix file**" (Fig 4). Users don't need to upload this file if users select "**Calculate Kinship (K) matrix by this software**", at this case, the K matrix can be calculate automatically. In Population Structure module, users should upload the Population Structure file if users select "**Included**" (Fig 5). There is no inclusion of population structure information in the genetic model if users select "**Not included in the model**". In Covariate module, users should upload the covariate file if users select "**Included**" (Fig 6). There is no inclusion of covariates in the genetic model if users select "**Not included in the model**".

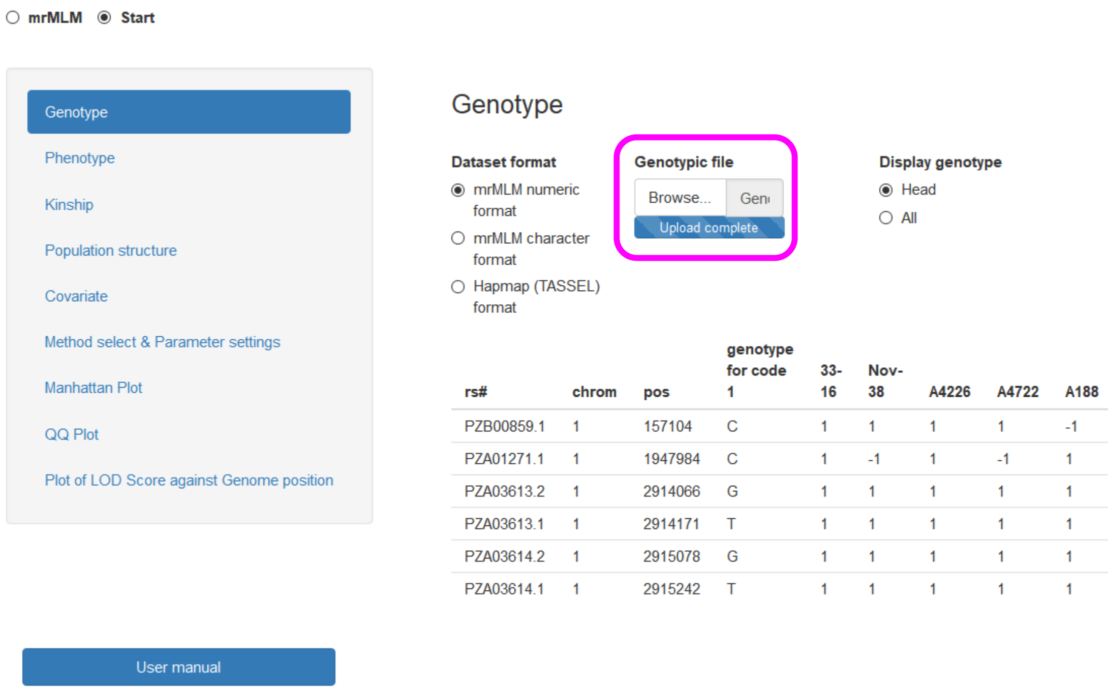


**Figure 2. Input genotypic dataset**

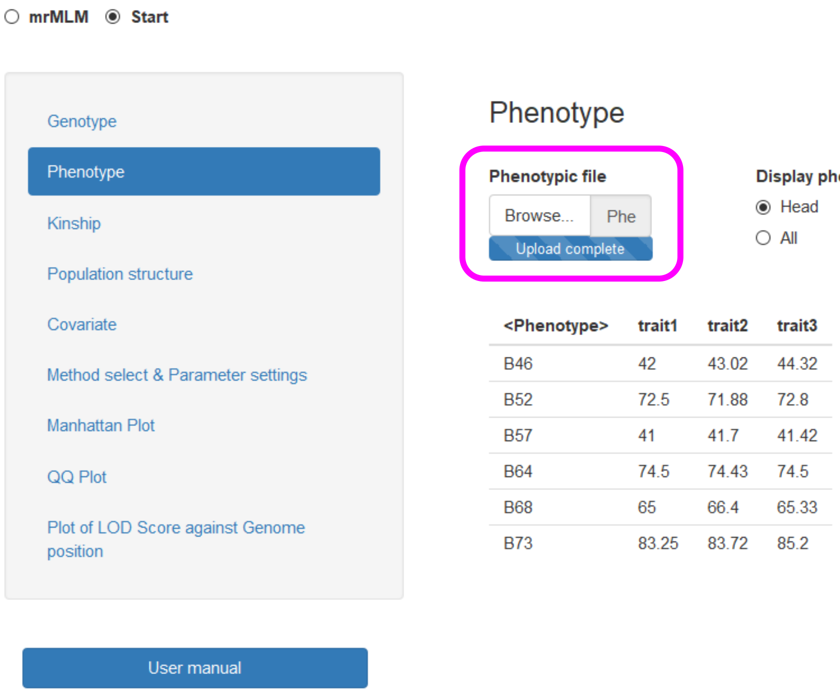


**Figure 3. Input Phenotypic dataset**

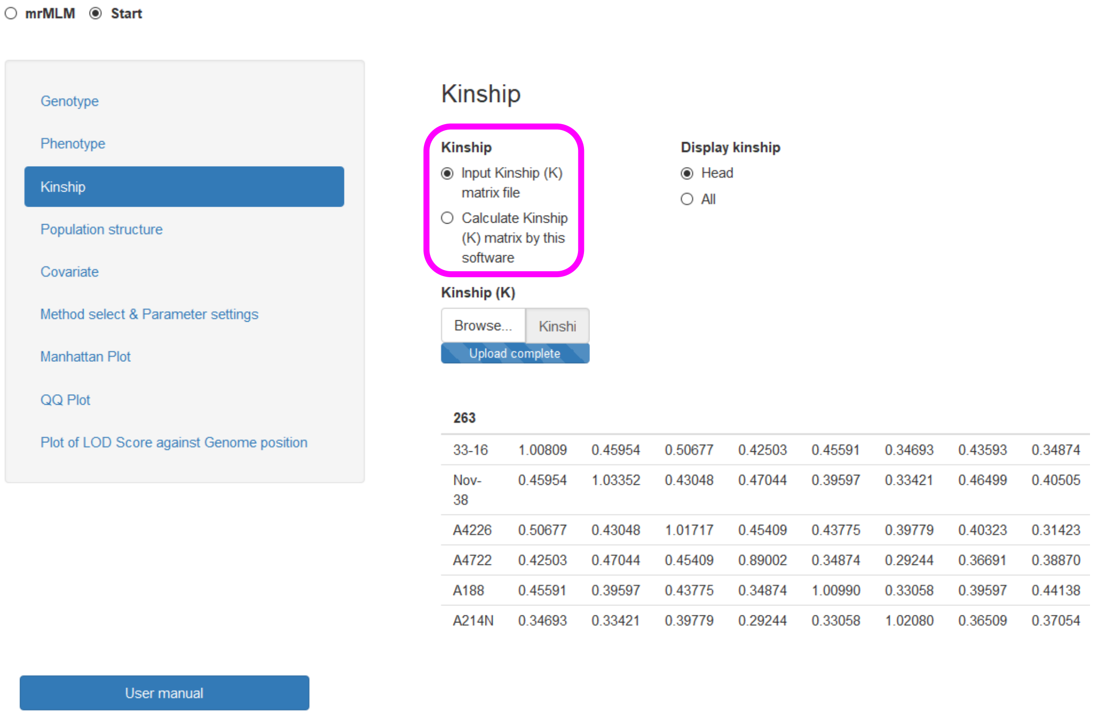


**Figure 4. Input kinship dataset**

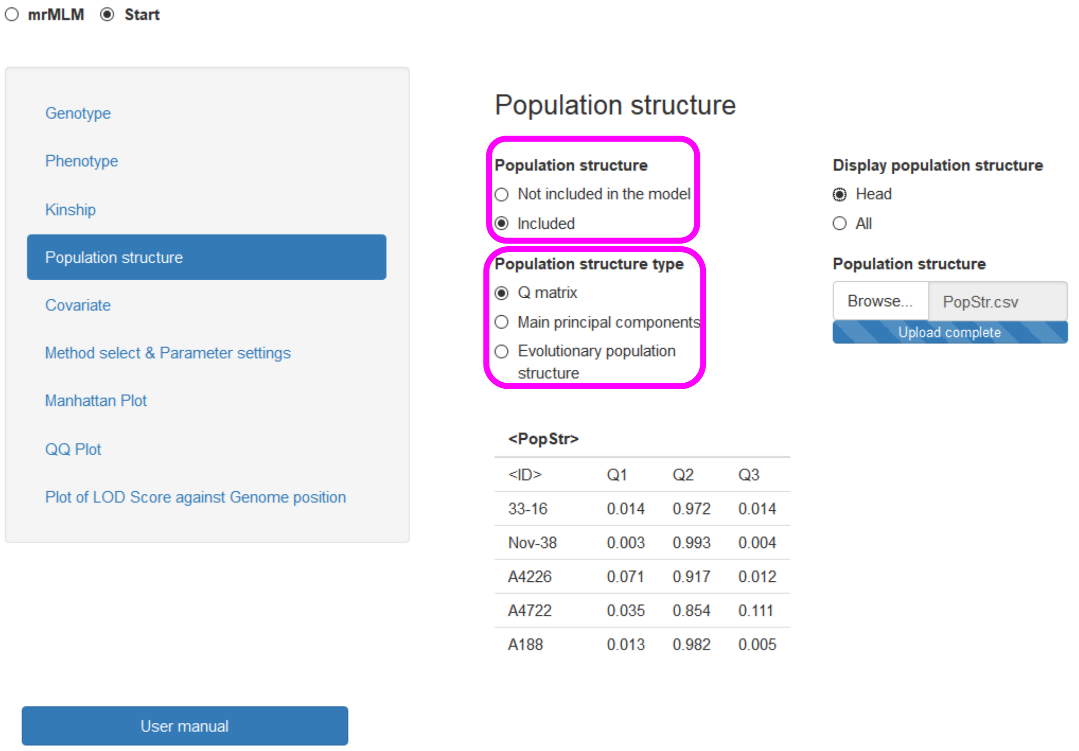


**Figure 5. Input Population Structure dataset**

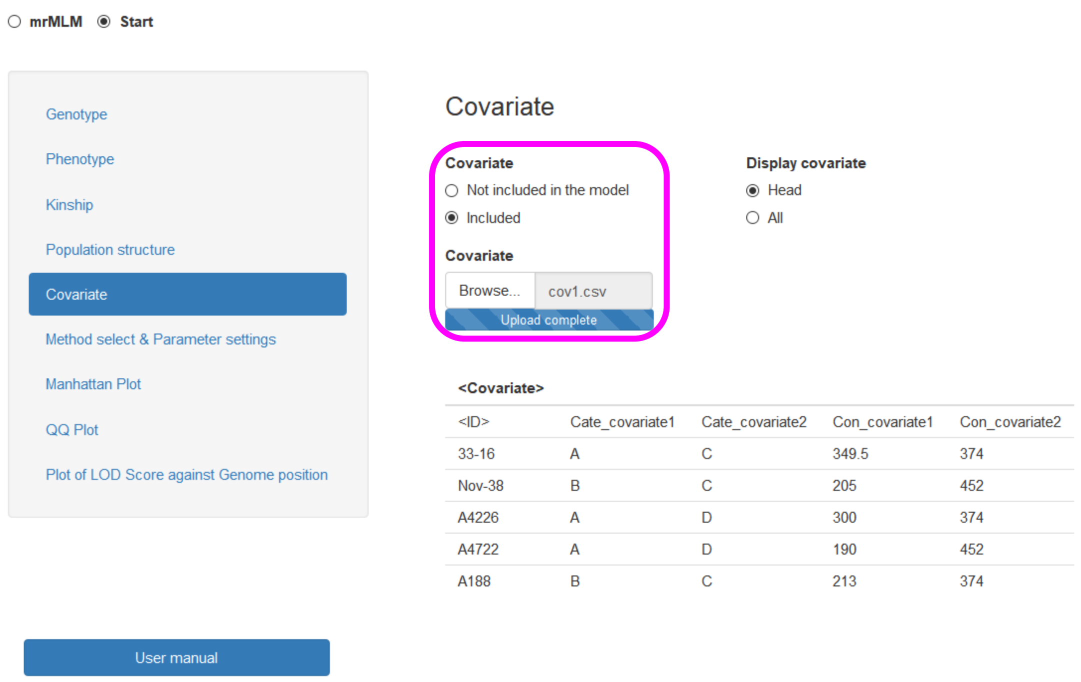


**Figure 6. Input Covariate dataset**

#### 3.3 Method select & Parameter setting (Fig 7)

**Method selection:** There are six multi-locus GWAS methods available in the mrMLM.GUI. Users may select one to six methods.

**Search radius of candidate gene (kb) (mrMLM & FASTmrMLM):** This parameter is only for mrMLM and FASTmrMLM, indicating Search Radius (kb) in search of potentially associated QTN. If users set it as 20 kb, only one potentially associated QTN within the radius of 20 kb may be selected into multi-locus model.

**Likelihood function (FASTmrEMMA):** This parameter is only for FASTmrEMMA, including restricted maximum likelihood (REML) and maximum likelihood (ML).

**No. of potentially associated variables selected by LARS (pLARmEB):** This parameter is only for pLARmEB. If users set it as 50, 50 potentially associated variables can be selected from each chromosome. Users may change this number in real data analysis in order to obtain the best results.

**Bootstrap (pLARmEB):** This parameter is only for pLARmEB, including **FALSE** & **TRUE**. **FALSE** indicates only the analysis of real dataset; **TRUE** indicates the analyses of both real and four resampling datasets.

**Save path:** Save path in your computer in order to output the results in this path.

**Traits analyzed:** Traits analyzed may be from number $n_1$ to number $n_2$. For example, "1:3" indicates that users analyze the first to third traits.

**Draw plot (All the methods):** Including **FALSE** and **TRUE**. **FALSE** indicates no figure output; **TRUE** indicates the output of figures, including the Manhattan, QQ and LOD score against genome position.

**Plot Resolution (All the methods):** Including **Low** and **High** for all the figure files. Their parameters are showed at Page 17.

**Plot format (All the methods):** Including *.jpeg, *.png, *.tiff and *.pdf for all the figure files.

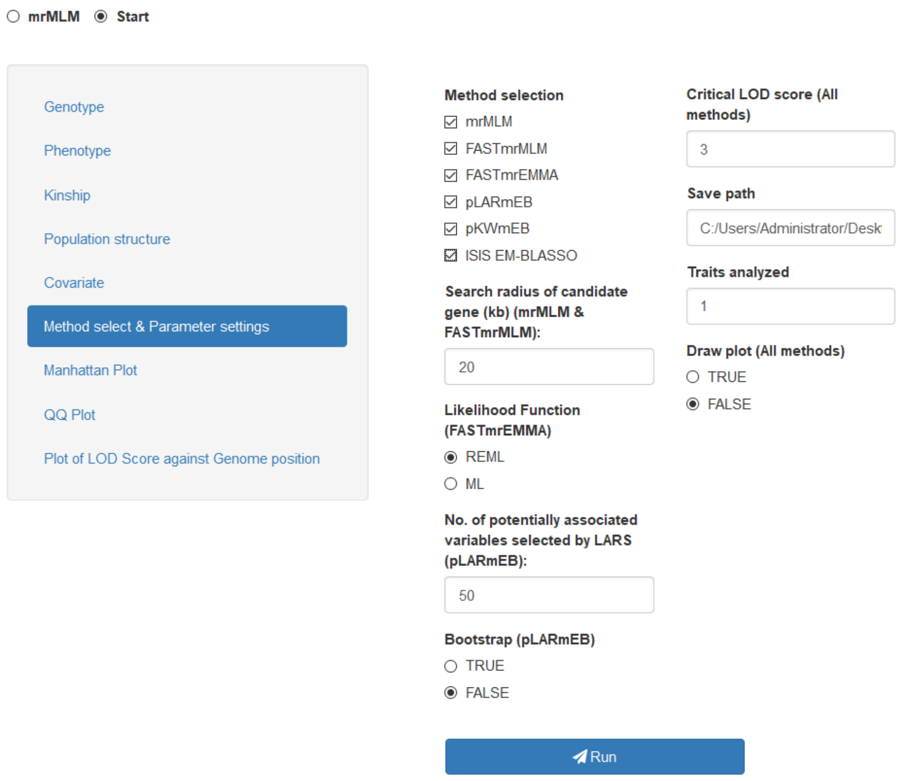


**Figure 7. Method select & Parameter setting**

##### 3.4 Run the software (Fig 8)

After uploading all the needed files and setting all the parameters, users can run the software. The result files will be saved to the path that users set up.

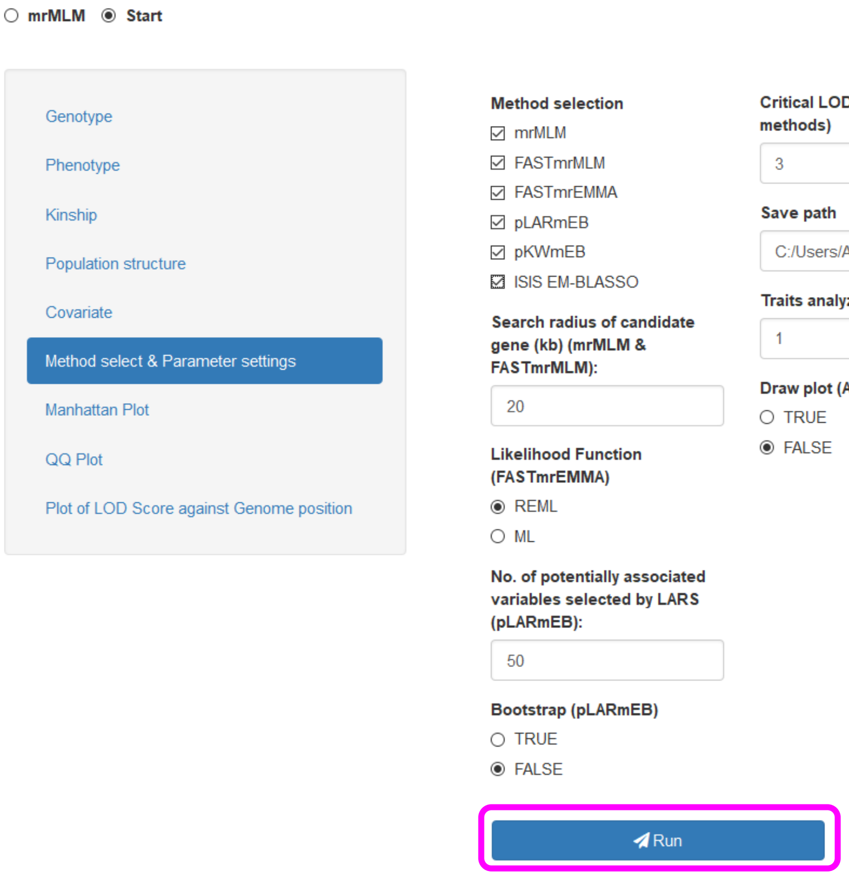


**Figure 8. Run the software mrMLM.GUI**

##### 3.5 Re-draw the plot according to user's requirement

Once users run the software, users can obtain a result file, named **resultforplot.xlsx,** which is used to redraw the plot.

###### 3.5.1 Manhattan plot

In the independent dialog window of Manhattan plot module, users can preview the Manhattan plot. Before saving this Figure, please set up the related parameters: **width** and **height** [with the unit of pixel (px)], **word resolution** [with the unit of 1/72 inch, being pixels per inch (ppi)], and **figure resolution** [with the unit of pixels per inch (ppi)]. Users may set up the colors for the adjacent chromosomes, with a drop-down option. The critical value for $-\log_{10}$(P-value) is defaulted as the value of $0.05/m_e$, where $m_e$ is the effective number of markers (please see Wang et al. *Scientific Reports* 2016,

6: 19444). Use “**Save Manhattan plot**” button to choose a path and to save the Figure, with four frequently used image formats: *.png, *.tiff , *.jpeg and *.pdf (**Fig 9**).

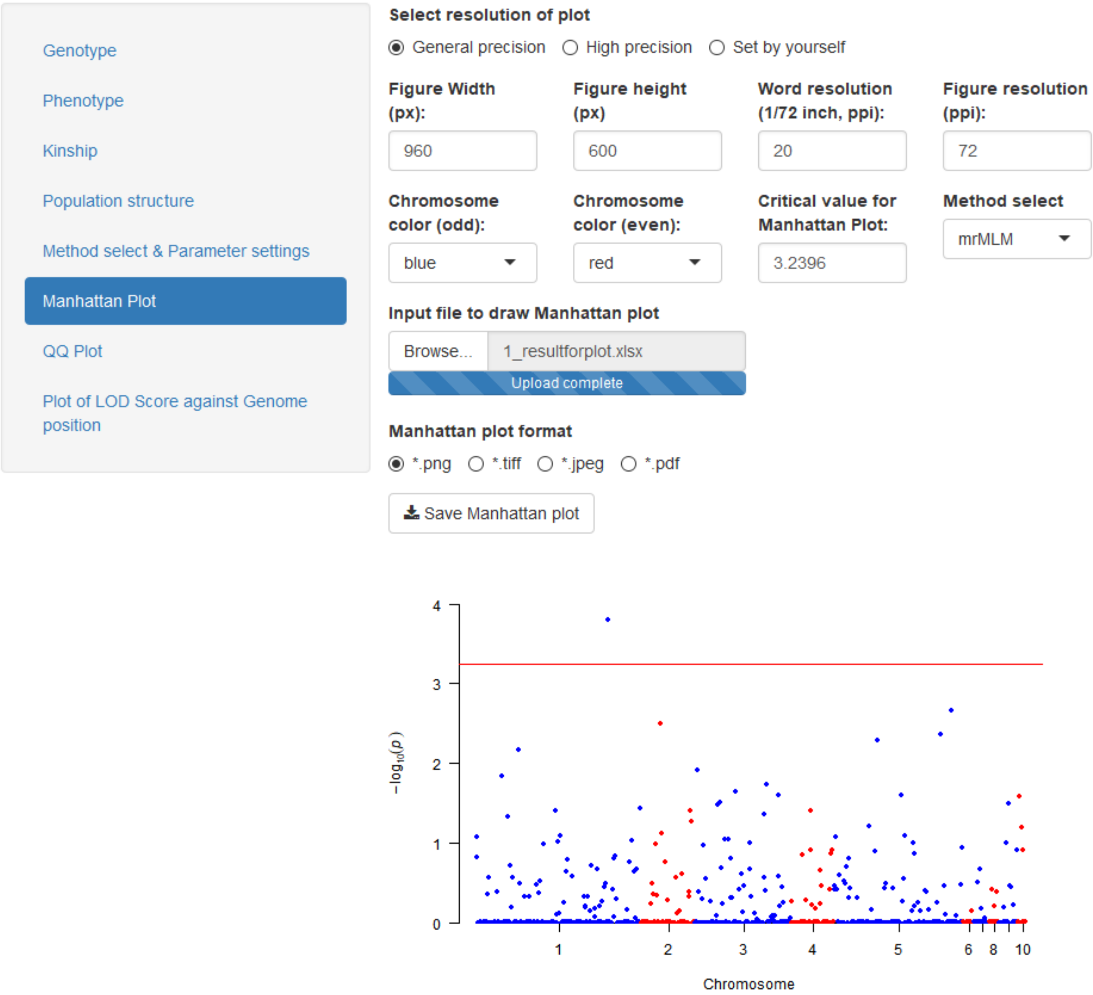


**Figure 9. Manhattan plot module**

##### 3.5.2 QQ plot

In the independent dialog window of **QQ plot** module, users can redraw the QQ plot. The parameter settings are the same as those in the Manhattan plot. Use “**Save QQ plot**” button to choose a path and to save the Figure, with four frequently used image formats: *.png, *.tiff , *.jpeg and *.pdf (**Fig 10**).

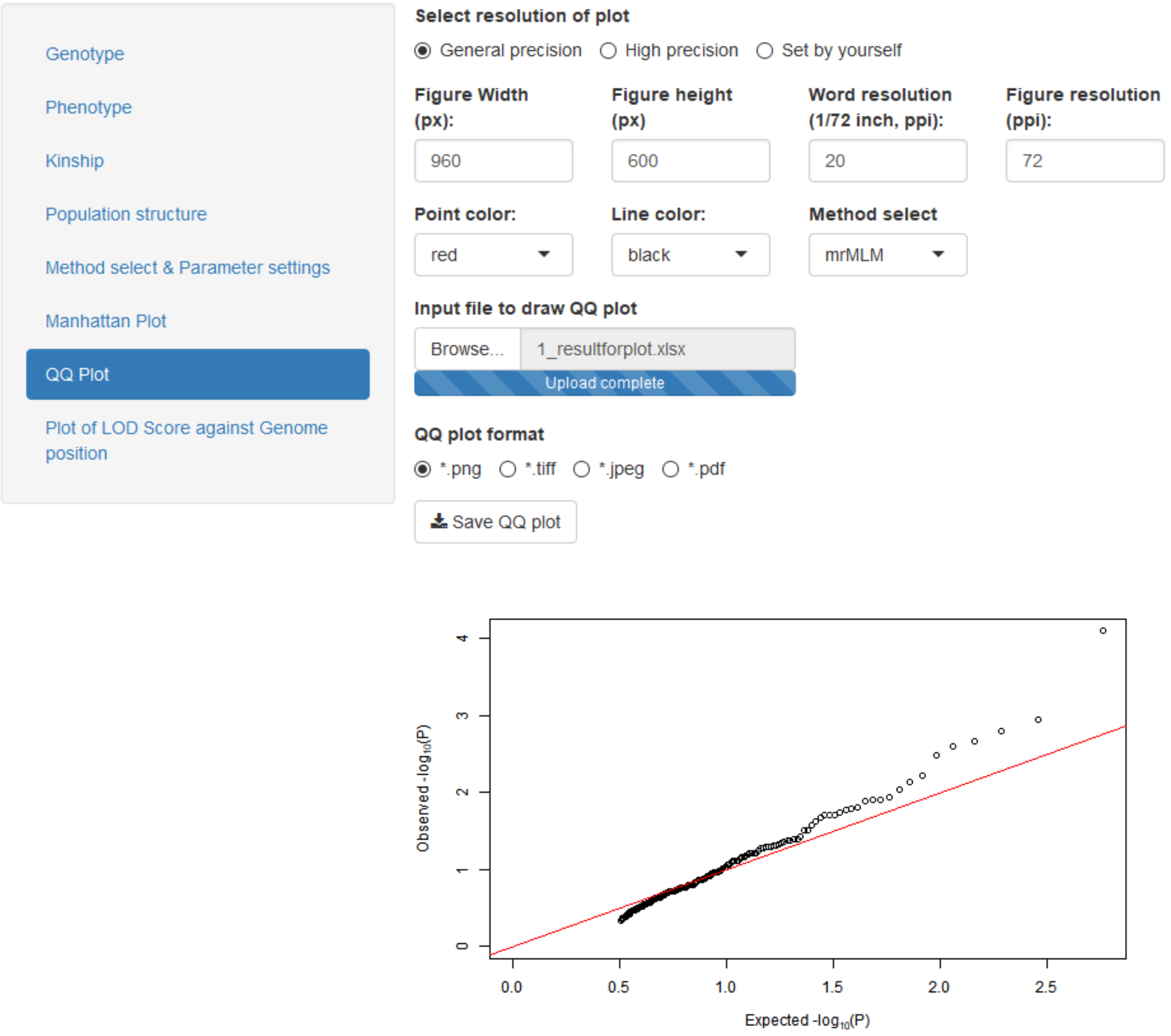


**Figure 10. QQ plot module**

##### 3.5.3 Plot of LOD score against genome position

In the "**Plot of LOD Score against Genome Position**" module, users can redraw the plot. The parameter settings are the same as those in the Manhattan plot. Users may set up the color of LOD line. Use the "**Save plot**" button to choose a path and to save the Figure, with four frequently used image formats: *.png, *.tiff, *.jpeg and *.pdf (**Fig 11**).

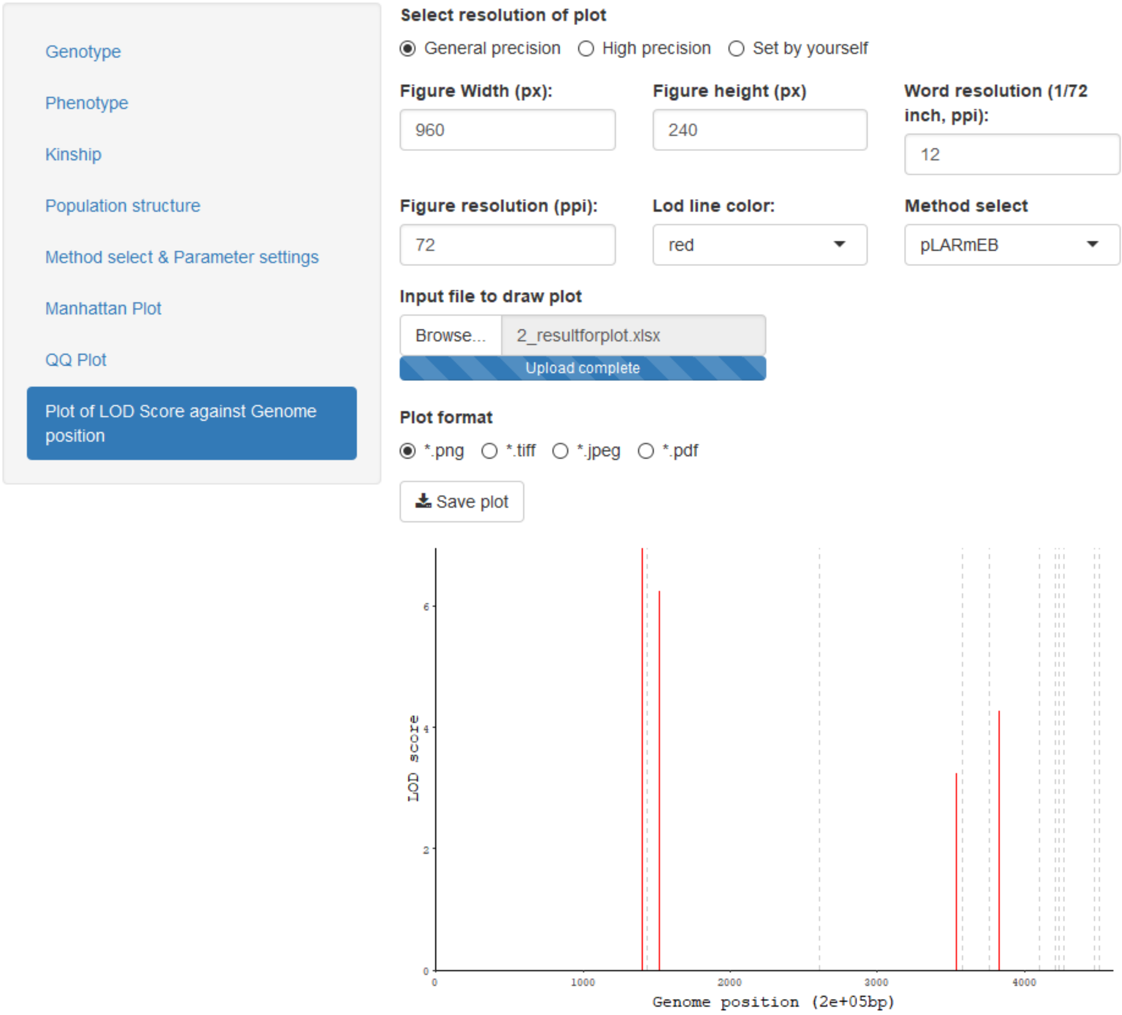


**Figure 11. Plot of LOD score against genome position (bp)**

#### 4. Result

At the work directory of user's R package, there will have two "results" files for the trait of interest, "1_intermediate result.csv" and "1_Final result.csv".

In the **intermediate result** from the mrMLM method, the result file includes: Trait ID, Trait name, method, reference sequence number (rs#, marker name), chromosome, marker's position (bp) in the chromosome, SNP effect ( $\gamma_k$ , Effect), -$\log_{10}$(P), genotype for code 1.

In the **Final result** from the mrMLM method, the result file includes: Trait ID, Trait name, method, reference sequence number (rs#, marker names), chromosome, marker's position (bp) in the chromosome, QTN effect, LOD score, -$\log_{10}$(P), the proportion of phenotypic variance explained by significant QTN ($r^2$), minor allelic frequency, genotype for code 1, residual error variance, and total phenotypic variance.

In the **plot results**, there are ten sheets, including "Manhattan mrMLM", "qq mrMLM", "Manhattan FASTmrMLM", "qq FASTmrMLM", "Manhattan FASTmrEMMA", "qq FASTmrEMMA", "Plot pLARmEB", "Manhattan pKWmEB", "qq pKWmEB", "Plot ISIS EM-BLASSO". These plot results will be saved to the ten sheets if users select all the methods. Users may upload this file into mrMLM.GUI in order to adjust all the figures based on user's opinions.

#### 5. References


1 Zhang Yuan-Ming, Mao Yongcai, Xie Chongqing, Howie Smith, Luo Lang, Xu Shizhong. Mapping quantitative trait loci using naturally occurring genetic variance among commercial inbred lines of maize (*Zea mays* L.). ***Genetics*** 2005, **169**: 2267–2275. DOI: 10.1534/genetics.104.033217

2 Wang Shi-Bo, Feng Jian-Ying, Ren Wen-Long, Huang Bo, Zhou Ling, Wen Yang-Jun, Zhang Jin, Jim M. Dunwell, Xu Shizhong*, Zhang Yuan-Ming*. Improving power and accuracy of genome-wide association studies via a multi-locus mixed linear model methodology. ***Scientific Reports*** 2016, **6**: 19444. DOI: 10.1038/srep19444

3 Tamba Cox Lwaka, Ni Yuan-Li, Zhang Yuan-Ming*. Iterative sure independence screening EM-Bayesian LASSO algorithm for multi-locus genome-wide association studies. ***PLoS Computational Biology*** 2017, **13**(1): e1005357, DOI: 10.1371/journal.pcbi.1005357

4 Zhang Jin[#], Feng Jian-Ying[#], Ni Yuan-Li, Wen Yang-Jun, Niu Yuan, Tamba Cox Lwaka, Yue Chao, Song Qi-Jian, Zhang Yuan-Ming*. pLARmEB: Integration of least angle regression with empirical Bayes for multi-locus genome-wide association studies. ***Heredity*** 2017, **118**: 517–524. DOI:10.1038/hdy.2017.8

5 Ren Wen-Long[#], Wen Yang-Jun[#], Jim M. Dunwell, Zhang Yuan-Ming*. pKWmEB: Integration of Kruskal-Wallis test with empirical Bayes under polygenic background control for multi-locus genome-wide association study. ***Heredity*** 2018, **120**: 208–218. https://doi.org/10.1038/s41437-017-0007-4

6 Wen Yang-Jun, Zhang Hanwen, Ni Yuan-Li, Huang Bo, Zhang Jin, Feng Jian-Ying, Wang Shi-Bo, Jim M. Dunwell, Zhang Yuan-Ming*, Wu Rongling*. Methodological implementation of mixed linear models in multi-locus genome-wide association studies. ***Briefings in Bioinformatics*** 2018, **19**(4): 700–712. https://doi.org/10.1093/bib/bbw145

7 Tamba Cox Lwaka, Zhang Yuan-Ming*. A fast mrMLM algorithm for multi-locus genome-wide association studies. *bioRxiv* 2018; doi: https://doi.org/10.1101/341784, online (June 7, 2018)

8 Zhang Ya-Wen, Tamba Cox Lwaka, Wen Yang-Jun, Li Pei, Ren Wen-Long, Ni Yuan-Li, Gao Jun, Zhang Yuan-Ming*. mrMLM v4.0: An R platform for multi-locus genome-wide association studies. ***Genomics, Proteomics & Bioinformatics***, resubmission
